# Transcriptional landscape of cardiac-specific *Gpx4* deletion recapitulates human cardiomyopathy

**DOI:** 10.64898/2026.03.27.714934

**Authors:** Alexandra M. Wiley, Xiaoyun Guo, Yi Chen, Eric Evangelista, Melissa A. Krueger, Qinghang Liu, Libin Xu, Sina A. Gharib, Rheem A. Totah

## Abstract

Glutathione peroxidase 4 (GPX4) is an antioxidant enzyme important for the reduction of toxic lipid peroxide products. Previous studies revealed the importance of mouse *Gpx4* in protecting cardiomyocytes from ferroptosis and, subsequently, the development of cardiovascular disease. In this paper, we investigate the transcriptional consequences of cardiac-specific deletion of *Gpx4* in mice and compare this response with that observed in human cardiomyopathy. The findings in this study highlight the importance of GPX4 in maintaining both structural and functional stability of the heart and identify key pathway changes resulting from excessive ferroptosis in cardiac tissue. By overlapping common transcriptional programs perturbed in this animal model and human cardiomyopathy, our findings identify putative mechanisms through which ferroptosis contributes to the development and progression of heart disease. These studies may help guide future cardiovascular therapeutics targeting ferroptosis-dependent pathways.

## Introduction

Lipid peroxidation is a free radical-mediated process where reactive oxygen species (ROS) react with polyunsaturated fatty acids (PUFAs) to generate a wide array of oxidation products, such as hydroperoxides (LOOH) and reactive lipid electrophiles. Lipid peroxidation plays an important role in the pathogenesis of cardiac remodeling and heart failure (HF) since the heart relies mostly on fatty acids (FA) as a source of energy (1). In patients with congestive HF, lipid peroxidation levels are significantly elevated and directly proportional to HF severity (2,3).

Glutathione peroxidase 4 (GPX4) is a selenium-dependent antioxidant enzyme that utilizes glutathione to reduce lipid hydroperoxides, preventing excessive lipid peroxidation and decreasing cellular oxidative damage (4). Experimental evidence in mice suggests that Gpx4 plays a key role in ischemia/reperfusion (I/R) injury, and that dosing mice with liproxstatin-1, an inhibitor of lipid peroxidation, protects the heart by restoring Gpx4 levels (5,6). Recently, GPX4 was identified as a key suppressor of ferroptosis (7,8), a recently defined form of cell death characterized by an iron-dependent accumulation of lipid peroxidation products, leading to irreparable lipid damage and membrane permeabilization. Emerging evidence suggests that ferroptosis plays an important role in the pathogenesis of HF induced by pressure overload, ischemic injury, and cardiomyopathy (9,10).

Among the other glutathione peroxidase isoforms, GPX4 is unique as it is the only peroxidase capable of reducing oxidized fatty acids and cholesterol hydroperoxides. Ablation of murine *Gpx4*, but not other *Gpx* isoforms, leads to embryonic lethality, highlighting an indispensable role for Gpx4 (11,12). Conditional deletion of *Gpx4* in neurons results in rapid onset of motor neuron degradation and paralysis in mice (13). Overexpression of *Gpx4* attenuates, whereas heterodeletion of *Gpx4* exacerbates, doxorubicin-induced cardiomyopathy and myocardial I/R injury (14,15). Moreover, *Gpx4* is markedly downregulated in the heart under pathological conditions such as myocardial infarction, pressure overload, and doxorubicin-induced cardiomyopathy (14–17).

By generating a cardiomyocyte-specific *Gpx4* knockout (KO) mouse model, we recently identified a key role for Gpx4 in myocardial homeostasis and remodeling (18). Importantly, ablation of *Gpx4* in the adult mouse heart induced dilated cardiomyopathy, which was rescued by ferroptosis inhibitors (18). Given that ablation of *Gpx4* is sufficient to induce ferroptosis in the heart *in vivo*, generation of a cardiac-specific *Gpx4* KO represents a novel model of ferroptosis-mediated cardiomyopathy.

Previous studies have highlighted the contribution of ferroptosis to various cardiovascular diseases (CVDs). However, the mechanisms through which ferroptosis contributes to HF remain poorly understood. In this paper, we leveraged our *Gpx4* conditional KO model to investigate ferroptosis-mediated pathways leading to HF and translated our findings in a more clinically relevant context by comparing them with transcriptional programs altered in human heart tissue from patients with cardiomyopathy.

## Methods

### Cardiomyocyte-specific Gpx4 KO mice

*Gpx4*-floxed mice were provided by Dr. Qitao Ran at the University of Texas Health Science Center at San Antonio (11). Cardiomyocyte-specific *Gpx4* KO mice were generated by crossing *Gpx4*-floxed mice with tamoxifen-inducible □MHC-MerCreMer mice (19). Mice were treated with tamoxifen (50 mg per kg body weight, i.p. for 5 days) to induce *Gpx4* gene deletion. Echocardiography and histology were used to evaluate the cardiac phenotype resulting from *Gpx4* gene deletion.

### Adult cardiac ventricular tissue

Surgical residual ventricular tissues from eighteen individuals (N = 18: 14 males, 4 females) with non-ischemic cardiomyopathy undergoing LVAD or transplant procedures were obtained from the University of Washington Medical Center. Since the tissue collected is anonymous, the Institutional Review Board at the University of Washington determined the samples were not human research (NHR) and waived the need for ethical review and informed consent. This policy is in accordance with Office for Human Research Protections guidelines (www.hhs.gov/ohrp/policy/cdebiol.html). Ventricular tissue was flash-frozen in liquid nitrogen immediately upon receipt and stored at −80 °C. Upon thawing, tissue was washed with phosphate buffered saline and immediately processed. Ventricular tissue from seventeen individuals (N = 12: 7 males, 5 females) with no known cardiac or cardiovascular health issues were obtained from BioIVT (Westbury, NY) and were used as the control group.

### Total RNA isolation

Heart tissue underwent manual homogenization with two sterile razor blades before being resuspended in 1mL TRIzol for 5 min at room temperature and placed in the Precellys24 (Bertin Instruments, Rockville, MD) at 6,800 RPM for 3 x 30 s with 60 s delay between cycles. Total RNA was then extracted using the PureLink RNA Isolation Kit (#12183018A, Invitrogen; Carlsbad, CA) by following the manufacturer’s Phase Separation with TRIzol Reagent protocol.

### RNA-sequencing (RNA-seq)

Following total RNA isolation, samples were shipped to Novogene Corporation Inc. (Sacramento, CA) where they underwent bulk mRNA-sequencing utilizing Illumina PE150 technology. Mouse and human samples were aligned to their respective reference genomes.

### Bioinformatics and pathway analyses

For each species, low-abundance genes were filtered if the maximum raw counts were less than 10. Differential gene expression analysis between conditions was performed using DESeq2 (R statistical environment) (20). Benjamini-Hochberg adjusted P-value (Padj) < 0.01 was used to designate significance and correct for multiple hypothesis testing.

To identify biological pathways altered by either cardiac-specific *Gpx4* KO in mice or human cardiomyopathy, we applied Gene Set Enrichment Analysis (GSEA) (21,22) using rank-ordered genes based on their DESeq2 test statistic and utilizing two Molecular Signature Database (MSigDB) categories: Hallmark and Canonical Pathways (23,24). We applied an FDR < 0.05 threshold for designating significant gene set enrichment. Network visualization of GSEA results was performed using the “enrichment map” feature in Cytoscape 3.10.2 (25).

### Comparison of murine and human data

Mouse Genome Database (26) was surveyed to identify mouse and human gene homologs and enable a more accurate examination of interspecies gene names. Common differentially expressed genes between the two datasets (Padj < 0.01) and common gene sets (FDR < 0.05) were identified and explored.

## Results

### Gpx4 knockout elicits profound transcriptional changes in murine heart

To assess the global effects of knocking out cardiomyocyte-specific *Gpx4* in mice, we performed principal component analysis (PCA) that showed distinct separation between wild type (WT) and KO mice indicating widespread transcriptional changes as a result of ablating *Gpx4* in the heart (**Supplementary Figure S1**). Next, we identified over 3,800 transcripts that were differentially expressed between *Gpx4* KO and WT mice (Figure 1A). Among these DEGs, we observed a number of genes well-known to be associated with an increased risk of CVD, including *Tnnt3, Nppb, Acta1*, and *Egln1*, as well as a number of ferroptosis-related genes, such as *Iscu, Sat1*, and *Cybb* (**Figure 1B**).

**Figure 1.**
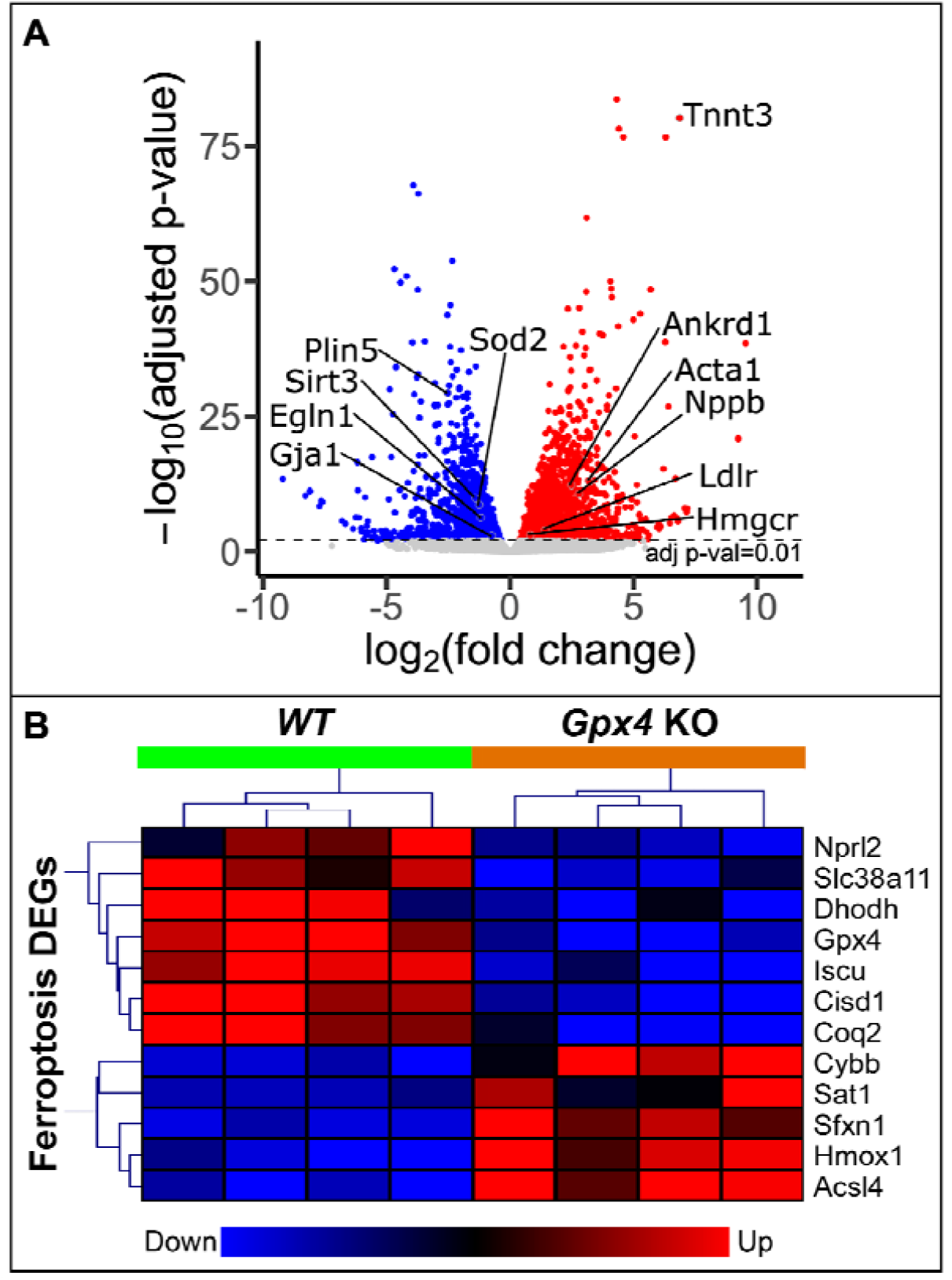
Identified DEGs with a Padj < 0.01 between WT and *Gpx4* KO mice. (**A**) Volcano plot of all the DEGs highlighting representative genes involved in cardiovascular disease and (**B**) a heatmap depiction of the expression patterns of ferroptosis-related DEGs.

To further elucidate the transcriptional consequences of knocking out *Gpx4* in mouse hearts, we applied GSEA to systematically identify altered pathways. We identified 788 upregulated pathways and 143 downregulated gene sets in *Gpx4* KO (FDR < 0.05). **Figure 2** depicts a volcano plot representation of the diverse pathways altered following *Gpx4* KO in mouse hearts. The most prominent upregulated processes in *Gpx4* KO mice were related to the immune response, development and remodeling (evidenced by extracellular matrix (ECM) organization), cell cycle, and regulated cell death. In contrast, significantly downregulated pathways mapped to oxidative phosphorylation (OXPHOS), the tricarboxylic acid (TCA) cycle, and ferroptosis regulation, among others. Interestingly, lipid-associated pathways had heterogeneous expression patterns, with many processes in fatty acid and lipid metabolism being downregulated, while both sphingolipid and cholesterol biosynthesis were upregulated. To better capture this diversity, we applied network analysis to the lipid-associated GSEA results to identify biological modules with distinct expression patterns (**Figure 3**).

**Figure 2.**
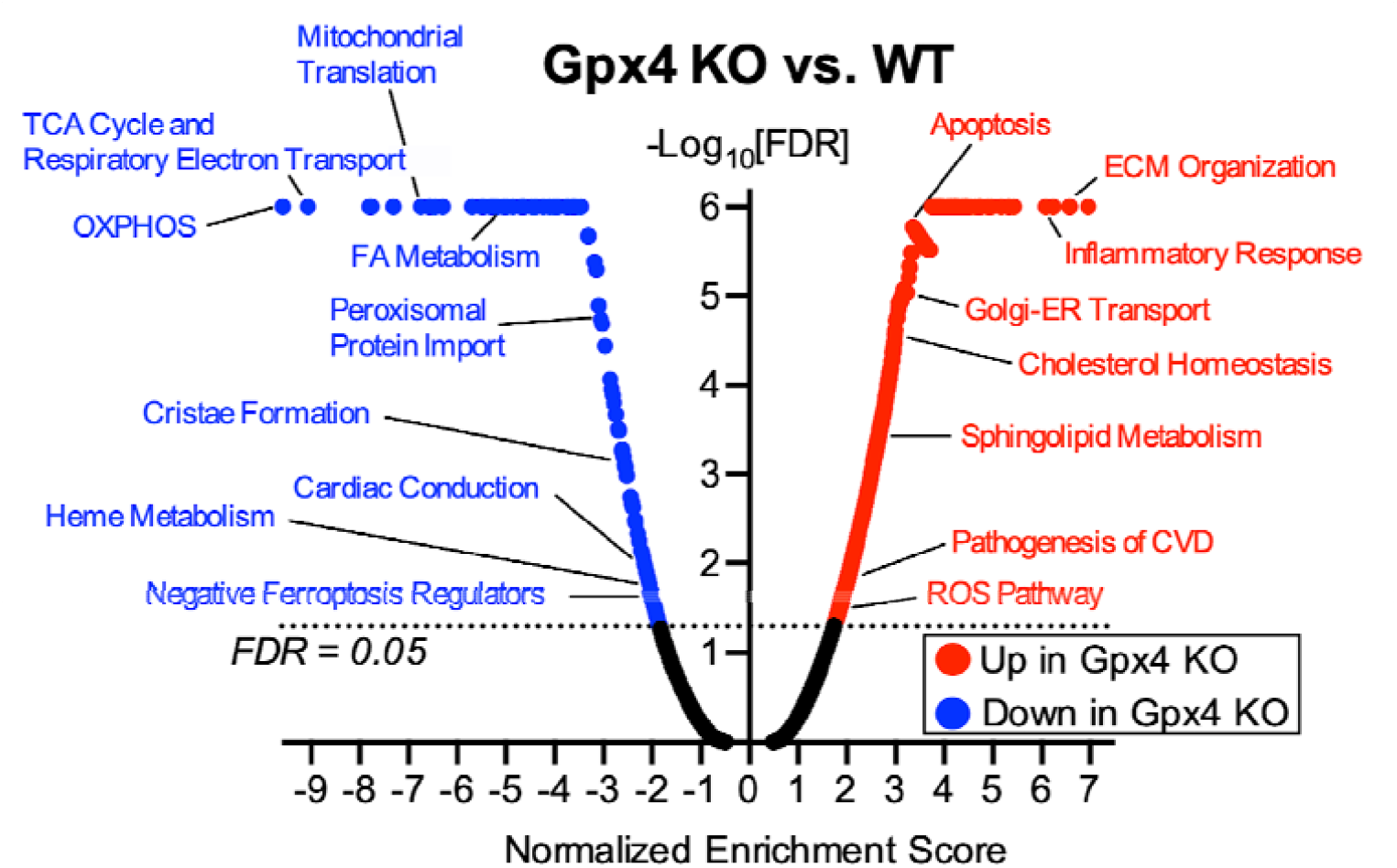
Volcano plot with labels encompassing pathways altered due to cardiomyocyte-specific *Gpx4* KO. The highlighted pathways are representative of the vast quantity of changes observed following *Gpx4* KO and further represent important pathways in heart function and ferroptosis. A full list of altered gene sets is included in Supplementary Table S1. TCA – tricarboxylic acid, OXPHOS – oxidative phosphorylation, FA – fatty acid, AA – amino acid, ECM – extracellular matrix, ER – endoplasmic reticulum, CVD – cardiovascular disease, ROS – reactive oxygen species

**Figure 3.**
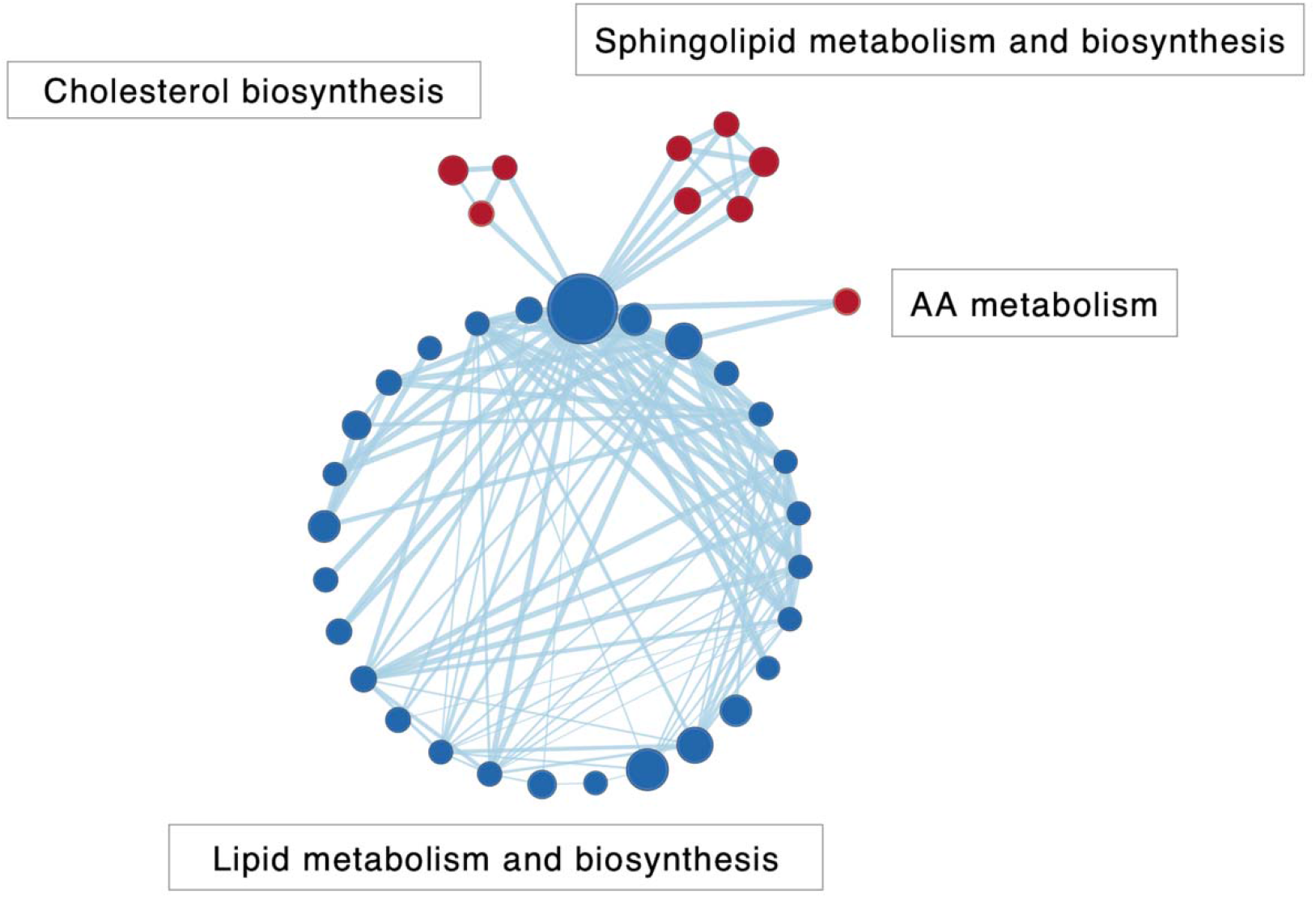
Network-based biological module demonstrating changes in lipid metabolism due to *Gpx4* KO. Red and blue nodes indicate upregulated and downregulated pathways, respectively, with connecting lines demonstrating greater than 50% gene overlap between nodes.

### Transcriptomic analysis of cardiac tissue from cardiomyopathy vs controls

Given the dramatic pathophysiological, structural, and transcriptional alterations observed in the hearts of *Gpx4* KO mice, we surveyed the transcriptome of human hearts with cardiomyopathy to place our findings in a more clinically relevant context. Thus, we performed a similar bulk RNA-seq analysis on ventricular cardiac tissue collected from 18 patients diagnosed with cardiomyopathy (CM) and 12 control subjects free of cardiovascular disease diagnoses (**Table 1**). PCA of this dataset showed clear separation between diseased patients and controls (**Supplementary Figure S2**), implying widespread transcriptomic differences between the two groups.

**Table 1.**
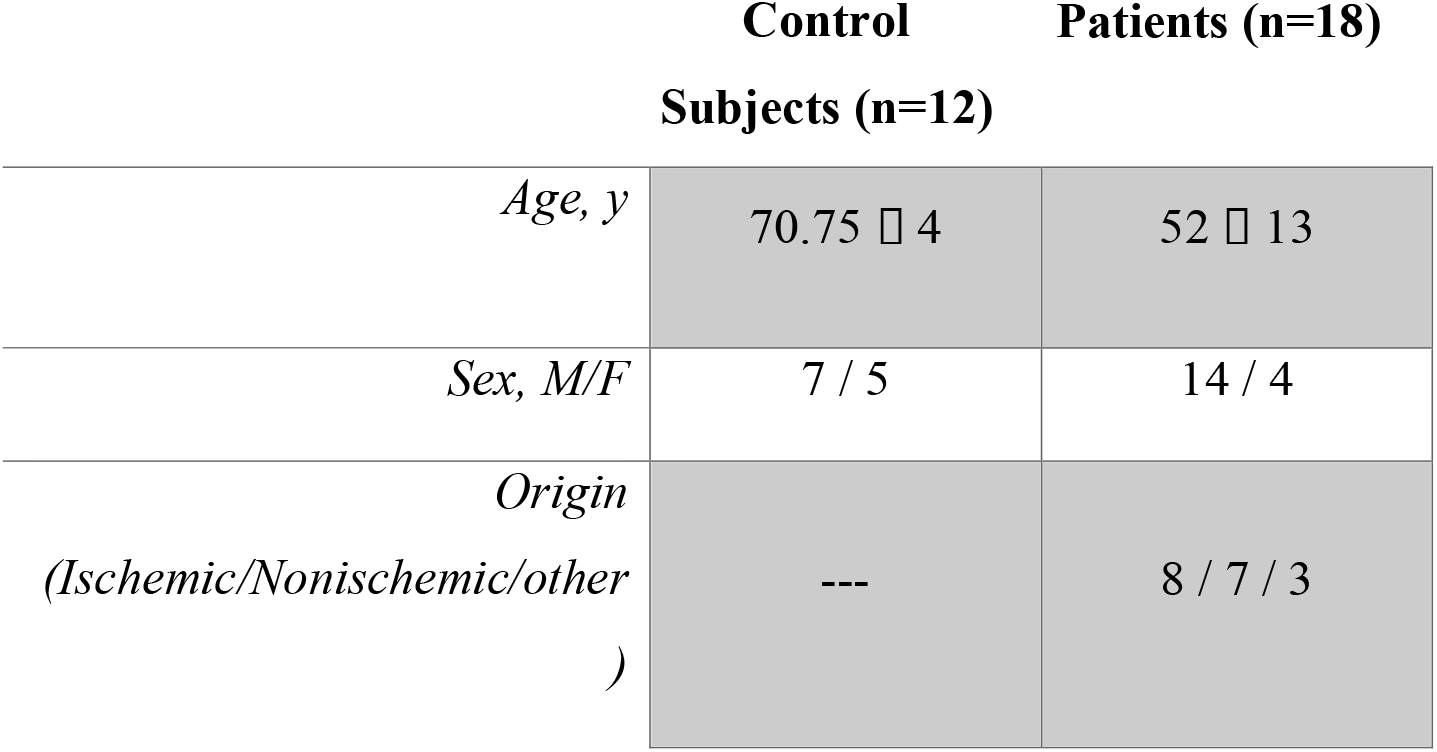
Known patient characteristics.

We identified over 6,000 DEGs between human CM and controls (Padj < 0.01). Of note, *GPX4* was decreased by 16% in CM with a P-value of 0.027. We next applied GSEA to reveal altered pathways due to CM. Although over half of the observed DEGs were downregulated, there were no significant downregulated gene sets observed, indicating that changes in the downregulated DEGs did not map to biologically coherent pathways. However, 527 gene sets were upregulated in CM (FDR < 0.05). A network-based summary of the GSEA results revealed the emergence of several biological modules composed of pathways involved in the immune response, development and remodeling, and the cell cycle among others. (**Figure 4**).

**Figure 4.**
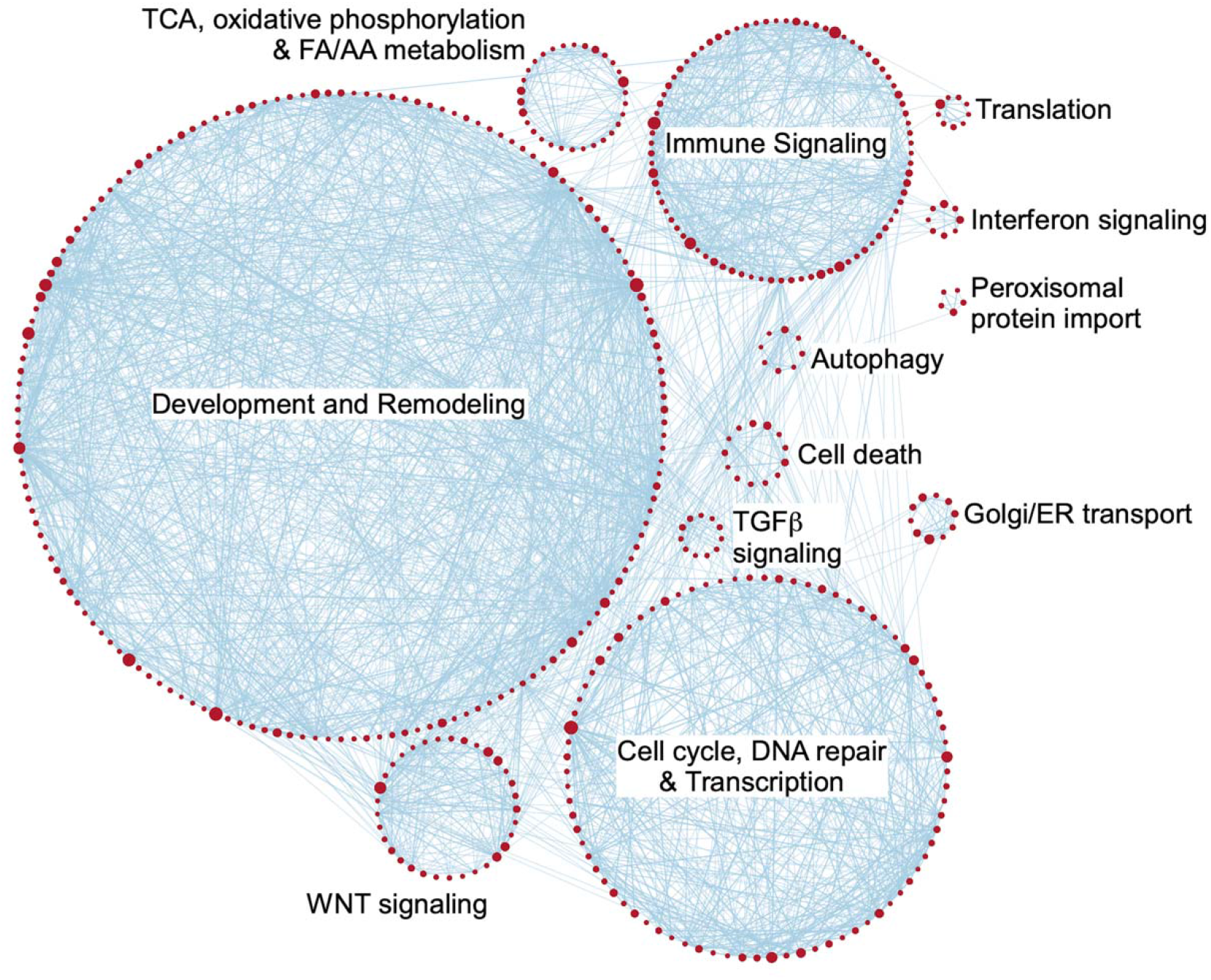
Network based visual depiction of GSEA results in human cardiomyopathy tissues compared to controls. The size of a node is proportional to the number of genes present in the pathway, while node connectivity between the gene set pairs demonstrates greater than 50% overlap among member genes. A full list of altered gene sets is included in Supplementary Table S2. TCA – tricarboxylic acid, FA – fatty acid, AA – amino acid, ER – endoplasmic reticulum

### Comparison of transcriptomic alterations due to cardiac-specific Gpx4 KO and human cardiomyopathy

We next compared the transcriptional changes between the human and murine datasets to assess how well the *Gpx4* KO model recapitulates features of human cardiomyopathy. We found ∼1,100 overlapping DEGs commonly altered in both *Gpx4* KO and human cardiomyopathy (Padj < 0.01). Within these, we noted significant downregulation of genes located on the mitochondrial chromosome, pointing to a common dysregulation of mitochondria in both *Gpx4* KO and human CM (**Figure 5**).

**Figure 5.**
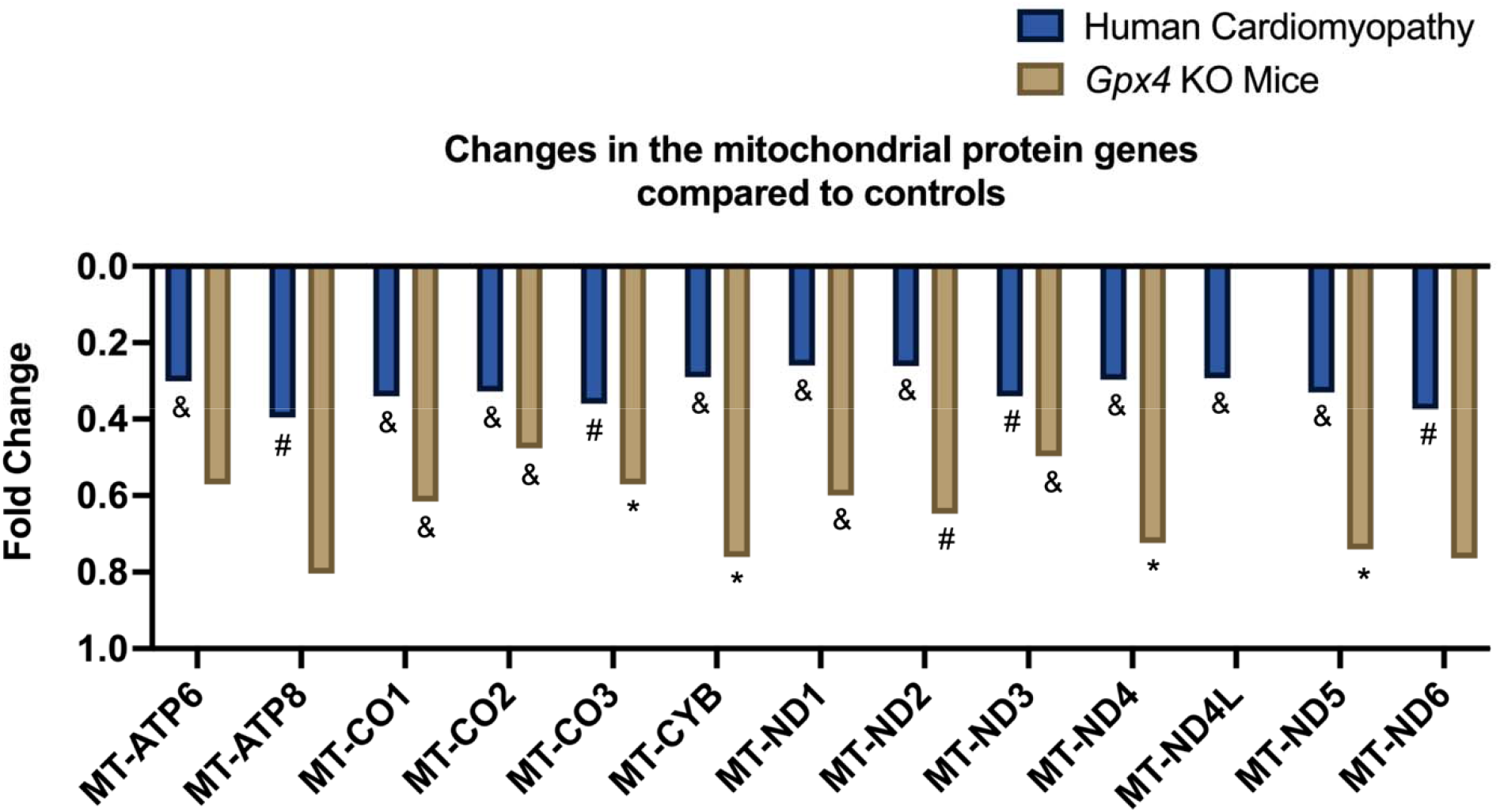
Changes in the mitochondrial protein genes. compared to controls following DESeq2 analysis. * Padj < 0.05, # Padj < 0.01, & Padj < 0.001.

More generally, when comparing pathway enrichment of the entire transcriptome using GSEA, we noted up to ∼40% concordance in pathway overlap (FDR < 0.05), suggesting that KO of *Gpx4* in mouse cardiomyocytes and human CM activate similar cellular compensatory mechanisms. Some of the most prominent similar enriched pathways included those involved in development and remodeling, activation of the immune system, and cell death (**Supplementary Table S3**). However, we also observed discordances between the two datasets in processes related to energy and lipid metabolism (**Supplementary Table S4**).

## Discussion

In this study, we compared a novel mouse conditional knockout model to human heart tissue to define the complex role of *Gpx4* in the pathogenesis of cardiomyopathy beyond its known regulation of ferroptosis. In the murine model, we found many differentially altered genes and pathways following *Gpx4* KO, indicating that GPX4 plays a central role in regulating cardiovascular homeostasis. Furthermore, there was significant overlap in DEGs and pathway changes between murine *Gpx4* KO and human cardiomyopathy, suggesting that targeting GPX4 may be a viable future treatment strategy to combat CM and CVD.

A transcriptomic analysis of the cardiovascular consequences of knocking out cardiac-specific *Gpx4* in mice identified a significant number of DEGs and pathway changes compared with wild-type controls, suggesting that GPX4 may play a broader role than simply suppressing ferroptosis in cardiac tissue. Following a DESeq2 analysis, we identified roughly 3,800 DEGs (Padj < 0.01), many of which suggested increased cardiac dysfunction, such as increased expression of *Tnnt3, Nppb, Ankrd1*, and *Hmgcr*. Additionally, the *Gpx4* KO mice had a decreased expression of *Plin5, Sirt3, Gja1*, and *Egln1*, further supporting a CVD phenotype. Additionally, we noted the upregulation of genes *Sfxn1, Hmox1, Sat1, Acsl4*, and *Cybb*, which were previously reported to increase in the presence of ferroptosis (27,28), as well as a decrease in genes that are thought to be protective against ferroptosis, such as *Gpx4, Iscu, Cisd1*, and *Coq2* (27,29) (**Figure 1B**).

Our pathway analysis revealed profound changes within the cardiomyocyte transcriptome consistent with widespread cellular dysfunction in cardiomyopathy. With *Gpx4* KO, we observed increases in ECM organization, apoptosis, and reactive oxygen species (ROS) generation (**Figure 2**). ECM organization is a hallmark of cardiac remodeling and a signature of CVD pathophysiology (30). Additionally, ROS accumulation is an amplifier for both apoptotic and ferroptotic cell death (31), and an increase in ROS concentrations in the myocardium has been associated with both the development and progression of CVD (32). Furthermore, we detected decreases in crucial pathways with roles in energy and lipid metabolism, as well as mitochondrial regulation in the *Gpx4* KO mice compared to the WT controls (**Figure 2**). These dysregulations suggest the cardiomyocytes of the *Gpx4* KO mice may be struggling to meet the same energy requirements as healthy. Since the heart relies heavily on FAs to maintain functionality (1,33–35). Recent reports have suggested that when cells undergo ferroptosis, and the antioxidant defense is dysfunctional, as is the case with *Gpx4* suppression, the cells can undergo metabolic switching, relying predominantly on the glycolysis pathway to support vital biological functions instead of OXPHOS (36). As we observed upregulated pathways involved in glycolysis with downregulated pathways crucial to pyruvate metabolism, the TCA cycle, and OXPHOS following *Gpx4* KO (**Supplementary Figure S1**), it appears that this metabolic switching could be occurring within the mouse cardiomyocytes. The exception to the widespread decrease in FA metabolism lies in the observed upregulation of cholesterol and sphingolipid metabolism and biosynthesis (**Figure 3**), warranting further experimental investigation. Taken together, the observed pathway alterations due to *Gpx4* KO are suggestive of increased cardiac dysfunction and cardiovascular disease.

Since the *Gpx4* KO mice exhibit distinct pathophysiological changes in the heart (18) leading to profound transcriptomic perturbations indicative of cardiac dysfunction, we extended our analysis to human samples to elucidate shared programs between the *Gpx4* KO model and human CM. We observed a significant number of overlapping DEGs and pathways between the two, suggesting a key link between GPX4 suppression and the development of CM across species. By comparing biological modules that emerged following network analysis of enriched pathways in the *Gpx4* KO and CM datasets, we found upregulation of pathways imperative to cardiovascular development and remodeling, Golgi transport, cell death, and the immune response. Beyond the well-recognized importance of cardiovascular development and remodeling, and cell death in CM, increased transport within the Golgi apparatus indicates dysfunction in protein processing and lipid homeostasis (37), both of which are important for cardiomyocyte homeostasis and heart function. The shared upregulation of these modules further supports that suppression of GPX4 leads to a cascade of molecular dysregulations similar to those observed in human CM.

Additionally, we observed similar downregulation of the 13 mitochondrial gene-encoded proteins that instruct the cell to produce protein subunits essential for OXPHOS (38) in both datasets (**Figure 5**). Notably, OXPHOS is under dual control by both the mitochondrial and nuclear genomes, of which the latter has majority control (39). While we found an upregulation of OXPHOS in the CM dataset, it is interesting that there was significant downregulation in the mtDNA OXPHOS genes, suggesting that the mitochondrial-encoded OXPHOS genes are downregulated in both *Gpx4* KO and CM. This finding points to a potentially common theme whereby mitochondrial dysregulation is a critical factor in both the *Gpx4* KO model and human CM. In terms of the *Gpx4* KO, the mitochondria are rich in membrane-bound unsaturated phospholipids, making this organelle a primary target for lipid peroxidation (40), so this dysregulation is not surprising. Of note, when taking both the mitochondrial and nuclear OXPHOS genes into consideration in the CM dataset, OXPHOS is primarily upregulated, suggesting overall differences in the OXPHOS regulation between the two disease models.

Several discordant results were also observed between the murine model and human CM. Differences were detected between the *Gpx4* KO and human cardiomyocyte pathway changes in the TCA cycle, OXPHOS, and the metabolism of FAs, which were downregulated following *Gpx4* KO but upregulated in CM. The current metabolic switching hypothesis, previously presented, could provide an explanation for this phenomenon. Other heart cell types may also contribute to ferroptosis and are not captured in the cardiomyocyte specific KO mouse model.

Another set of key differences fall under changes in mitochondrial translation and biogenesis, and in the peroxisome, which are decreased in *Gpx4* KO mice but increased in human CM. As both the mitochondria and peroxisomes are essential for FA oxidation, they are imperative organelles for proper cardiac function (41,42). In the case of the *Gpx4* KO, it has been suggested that as peroxisome expression decreases, ferroptosis susceptibility increases (43). Alternatively, within human CM, it is possible that both the mitochondrial and peroxisomal pathways are being upregulated in a feedback mechanism to overcome the contractility issues associated with CM (41,42). Taken together, we speculate that when mouse cardiomyocytes undergo ferroptosis, they may be unable to meet the ATP demands of the heart and thus undergo heart failure, while in human CM, the heart structure is unable to meet the functional demands, rather than the energy demands. However, this speculation needs to be further validated experimentally. Although lipid metabolism changes are discordant between the *Gpx4* KO and CM datasets, there are similar trends of upregulation in cholesterol and sphingolipid metabolism and biosynthesis. This finding suggests that these lipid species react similarly with the suppression of Gpx4 and CM, and further experimental efforts are needed to determine the significance of these lipid species during ferroptosis.

There were several limitations in our study. The human heart tissues used in this analysis were at end-stage heart failure, whereas the *Gpx4* KO mice, while exhibiting dilated cardiomyopathy (18), were not at an equivalent end-stage disease state. Furthermore, the murine model captures one potential path to cardiomyopathy, while many such pathophysiologic perturbations occur in human CM. This may also partially explain why, despite substantial overlaps in DEGs and enriched pathways, we also found significant discrepancies between human CM and the animal model. Nevertheless, using genetically modified mouse models, despite their limitations, has made important contributions to our understanding of key molecular mechanisms in human heart disease (44).

In conclusion, using an agnostic approach, we demonstrate many similarities in the transcriptional landscape between human cardiomyopathy and conditional deletion of *Gpx4* in mice, while also highlighting several differences in cardiac energetics. The observed upregulation in cardiomyocyte remodeling and cell death, paired with downregulation of energy and lipid metabolism, sheds further light on the importance of GPX4 in the development and progression of CVD and identifies GPX4 as a potential therapeutic target to combat heart disease.

## Supporting information

table of differentially expressed genes

