## Supplementary material for "Transcriptional landscape of cardiac-specific *Gpx4* deletion recapitulates human cardiomyopathy": table of differentially expressed genes

**Supplementary Table S1.** List of differentially altered gene sets in *Gpx4 KO* mouse hearts (FDR < 0.05).

| **Gene Set Name** | **Normalized Enrichment Score (NES)** | **FDR** |
| --- | --- | --- |
| HALLMARK_EPITHELIAL_MESENCHYMAL_TRANSITION | 6.98 | 0.00E+00 |
| HALLMARK_TNFA_SIGNALING_VIA_NFKB | 6.63 | 0.00E+00 |
| REACTOME_EXTRACELLULAR_MATRIX_ORGANIZATION | 6.59 | 0.00E+00 |
| REACTOME_NEUTROPHIL_DEGRANULATION | 6.12 | 0.00E+00 |
| HALLMARK_INFLAMMATORY_RESPONSE | 6.09 | 0.00E+00 |
| REACTOME_SIGNALING_BY_RECEPTOR_TYROSINE_KINASES | 5.33 | 0.00E+00 |
| NABA_MATRISOME_ASSOCIATED | 5.24 | 0.00E+00 |
| REACTOME_SIGNALING_BY_INTERLEUKINS | 5.22 | 0.00E+00 |
| REACTOME_HEMOSTASIS | 5.19 | 0.00E+00 |
| HALLMARK_ALLOGRAFT_REJECTION | 5.14 | 0.00E+00 |
| REACTOME_RHO_GTPASE_EFFECTORS | 4.90 | 0.00E+00 |
| KEGG_REGULATION_OF_ACTIN_CYTOSKELETON | 4.82 | 0.00E+00 |
| HALLMARK_APICAL_JUNCTION | 4.75 | 0.00E+00 |
| WP_TYROBP_CAUSAL_NETWORK_IN_MICROGLIA | 4.63 | 0.00E+00 |
| REACTOME_INTERLEUKIN_10_SIGNALING | 4.55 | 0.00E+00 |
| REACTOME_COLLAGEN_FORMATION | 4.53 | 0.00E+00 |
| KEGG_CYTOKINE_CYTOKINE_RECEPTOR_INTERACTION | 4.53 | 0.00E+00 |
| HALLMARK_IL2_STAT5_SIGNALING | 4.48 | 0.00E+00 |
| KEGG_FOCAL_ADHESION | 4.43 | 0.00E+00 |
| REACTOME_DEGRADATION_OF_THE_EXTRACELLULAR_MATRIX | 4.42 | 0.00E+00 |
| REACTOME_IMMUNOREGULATORY_INTERACTIONS_BETWEEN_A_LYMPHOID_AND_A_NON_LYMPHOID_CELL | 4.40 | 0.00E+00 |
| REACTOME_CELL_SURFACE_INTERACTIONS_AT_THE_VASCULAR_WALL | 4.31 | 0.00E+00 |
| REACTOME_ELASTIC_FIBRE_FORMATION | 4.30 | 0.00E+00 |
| REACTOME_INTEGRIN_CELL_SURFACE_INTERACTIONS | 4.27 | 0.00E+00 |
| PID_INTEGRIN1_PATHWAY | 4.24 | 0.00E+00 |
| WP_BURN_WOUND_HEALING | 4.22 | 0.00E+00 |
| HALLMARK_IL6_JAK_STAT3_SIGNALING | 4.21 | 0.00E+00 |
| REACTOME_CELL_CYCLE_MITOTIC | 4.18 | 0.00E+00 |
| REACTOME_INTERLEUKIN_4_AND_INTERLEUKIN_13_SIGNALING | 4.17 | 0.00E+00 |
| REACTOME_COLLAGEN_BIOSYNTHESIS_AND_MODIFYING_ENZYMES | 4.17 | 0.00E+00 |
| WP_FOCAL_ADHESION | 4.16 | 0.00E+00 |
| KEGG_LEUKOCYTE_TRANSENDOTHELIAL_MIGRATION | 4.13 | 0.00E+00 |
| HALLMARK_KRAS_SIGNALING_UP | 4.13 | 0.00E+00 |
| REACTOME_ECM_PROTEOGLYCANS | 4.09 | 0.00E+00 |
| WP_MICROGLIA_PATHOGEN_PHAGOCYTOSIS_PATHWAY | 4.09 | 0.00E+00 |
| WP_PLATELETMEDIATED_INTERACTIONS_WITH_VASCULAR_AND_CIRCULATING_CELLS | 4.09 | 0.00E+00 |
| PID_CXCR4_PATHWAY | 4.06 | 0.00E+00 |
| REACTOME_COLLAGEN_DEGRADATION | 3.96 | 0.00E+00 |
| NABA_CORE_MATRISOME | 3.91 | 0.00E+00 |
| HALLMARK_E2F_TARGETS | 3.90 | 0.00E+00 |
| HALLMARK_COMPLEMENT | 3.89 | 0.00E+00 |
| REACTOME_EPH_EPHRIN_SIGNALING | 3.89 | 0.00E+00 |
| REACTOME_ASSEMBLY_OF_COLLAGEN_FIBRILS_AND_OTHER_MULTIMERIC_STRUCTURES | 3.87 | 0.00E+00 |
| NABA_ECM_REGULATORS | 3.84 | 0.00E+00 |
| REACTOME_SYNDECAN_INTERACTIONS | 3.83 | 0.00E+00 |
| HALLMARK_G2M_CHECKPOINT | 3.83 | 0.00E+00 |
| HALLMARK_ANDROGEN_RESPONSE | 3.82 | 0.00E+00 |
| KEGG_CHEMOKINE_SIGNALING_PATHWAY | 3.79 | 0.00E+00 |
| REACTOME_PLATELET_ACTIVATION_SIGNALING_AND_AGGREGATION | 3.78 | 0.00E+00 |
| WP_VEGFAVEGFR2_SIGNALING | 3.72 | 0.00E+00 |
| WP_NETWORK_MAP_OF_SARSCOV2_SIGNALING_PATHWAY | 3.71 | 0.00E+00 |
| HALLMARK_INTERFERON_GAMMA_RESPONSE | 3.70 | 0.00E+00 |
| WP_LDL_INFLUENCE_ON_CD14_AND_TLR4 | 3.67 | 0.00E+00 |
| WP_RETINOBLASTOMA_GENE_IN_CANCER | 3.66 | 0.00E+00 |
| REACTOME_MITOTIC_G1_PHASE_AND_G1_S_TRANSITION | 3.65 | 0.00E+00 |
| WP_TYPE_I_COLLAGEN_SYNTHESIS_IN_THE_CONTEXT_OF_OSTEOGENESIS_IMPERFECTA | 3.63 | 0.00E+00 |
| PID_AVB3_INTEGRIN_PATHWAY | 3.61 | 0.00E+00 |
| REACTOME_CELL_CYCLE_CHECKPOINTS | 3.61 | 0.00E+00 |
| WP_IL18_SIGNALING_PATHWAY | 3.60 | 0.00E+00 |
| WP_TOLLLIKE_RECEPTOR_SIGNALING_PATHWAY | 3.57 | 0.00E+00 |
| WP_FIBRIN_COMPLEMENT_RECEPTOR_3_SIGNALING_PATHWAY | 3.57 | 0.00E+00 |
| REACTOME_MOLECULES_ASSOCIATED_WITH_ELASTIC_FIBRES | 3.56 | 0.00E+00 |
| PID_AMB2_NEUTROPHILS_PATHWAY | 3.56 | 0.00E+00 |
| WP_IL1_AND_MEGAKARYOCYTES_IN_OBESITY | 3.56 | 0.00E+00 |
| PID_PDGFRB_PATHWAY | 3.56 | 0.00E+00 |
| KEGG_TOLL_LIKE_RECEPTOR_SIGNALING_PATHWAY | 3.56 | 0.00E+00 |
| REACTOME_SIGNALING_BY_ROBO_RECEPTORS | 3.52 | 0.00E+00 |
| REACTOME_RAC2_GTPASE_CYCLE | 3.51 | 0.00E+00 |
| REACTOME_RAC1_GTPASE_CYCLE | 3.51 | 0.00E+00 |
| REACTOME_RHO_GTPASE_CYCLE | 3.49 | 0.00E+00 |
| KEGG_CELL_ADHESION_MOLECULES_CAMS | 3.49 | 0.00E+00 |
| REACTOME_SARS_COV_INFECTIONS | 3.48 | 0.00E+00 |
| WP_CHEMOKINE_SIGNALING_PATHWAY | 3.47 | 0.00E+00 |
| WP_REGULATION_OF_ACTIN_CYTOSKELETON | 3.47 | 0.00E+00 |
| REACTOME_RHO_GTPASES_ACTIVATE_FORMINS | 3.45 | 0.00E+00 |
| HALLMARK_P53_PATHWAY | 3.45 | 0.00E+00 |
| REACTOME_NON_INTEGRIN_MEMBRANE_ECM_INTERACTIONS | 3.44 | 0.00E+00 |
| WP_EBOLA_VIRUS_INFECTION_IN_HOST | 3.44 | 0.00E+00 |
| KEGG_ENDOCYTOSIS | 3.44 | 0.00E+00 |
| REACTOME_PARASITE_INFECTION | 3.44 | 0.00E+00 |
| KEGG_FC_GAMMA_R_MEDIATED_PHAGOCYTOSIS | 3.43 | 0.00E+00 |
| WP_NEURAL_CREST_CELL_MIGRATION_DURING_DEVELOPMENT | 3.42 | 0.00E+00 |
| SA_MMP_CYTOKINE_CONNECTION | 3.40 | 0.00E+00 |
| NABA_COLLAGENS | 3.40 | 0.00E+00 |
| REACTOME_FCGAMMA_RECEPTOR_FCGR_DEPENDENT_PHAGOCYTOSIS | 3.40 | 0.00E+00 |
| REACTOME_COLLAGEN_CHAIN_TRIMERIZATION | 3.39 | 0.00E+00 |
| KEGG_ECM_RECEPTOR_INTERACTION | 3.39 | 0.00E+00 |
| WP_MEASLES_VIRUS_INFECTION | 3.39 | 0.00E+00 |
| REACTOME_MITOTIC_METAPHASE_AND_ANAPHASE | 3.38 | 0.00E+00 |
| WP_SPINAL_CORD_INJURY | 3.37 | 0.00E+00 |
| WP_COMPLEMENT_SYSTEM_IN_NEURONAL_DEVELOPMENT_AND_PLASTICITY | 3.37 | 0.00E+00 |
| HALLMARK_MTORC1_SIGNALING | 3.35 | 0.00E+00 |
| REACTOME_REGULATION_OF_EXPRESSION_OF_SLITS_AND_ROBOS | 3.35 | 0.00E+00 |
| KEGG_HEMATOPOIETIC_CELL_LINEAGE | 3.35 | 0.00E+00 |
| REACTOME_LEISHMANIA_INFECTION | 3.34 | 0.00E+00 |
| REACTOME_ASPARAGINE_N_LINKED_GLYCOSYLATION | 3.32 | 0.00E+00 |
| HALLMARK_APOPTOSIS | 3.32 | 0.00E+00 |
| PID_SYNDECAN_1_PATHWAY | 3.30 | 0.00E+00 |
| REACTOME_RESPONSE_TO_ELEVATED_PLATELET_CYTOSOLIC_CA2 | 3.30 | 0.00E+00 |
| WP_OVERVIEW_OF_PROINFLAMMATORY_AND_PROFIBROTIC_MEDIATORS | 3.30 | 0.00E+00 |
| REACTOME_ANTIGEN_PROCESSING_CROSS_PRESENTATION | 3.29 | 0.00E+00 |
| REACTOME_TRAFFICKING_AND_PROCESSING_OF_ENDOSOMAL_TLR | 3.28 | 0.00E+00 |
| PID_INTEGRIN_A4B1_PATHWAY | 3.27 | 0.00E+00 |
| REACTOME_RESOLUTION_OF_SISTER_CHROMATID_COHESION | 3.27 | 0.00E+00 |
| PID_UPA_UPAR_PATHWAY | 3.26 | 0.00E+00 |
| PID_INTEGRIN3_PATHWAY | 3.24 | 0.00E+00 |
| WP_MALIGNANT_PLEURAL_MESOTHELIOMA | 3.23 | 0.00E+00 |
| REACTOME_MHC_CLASS_II_ANTIGEN_PRESENTATION | 3.22 | 0.00E+00 |
| WP_G1_TO_S_CELL_CYCLE_CONTROL | 3.21 | 0.00E+00 |
| HALLMARK_MYC_TARGETS_V1 | 3.21 | 0.00E+00 |
| PID_SYNDECAN_4_PATHWAY | 3.21 | 0.00E+00 |
| WP_TROP2_REGULATORY_SIGNALING | 3.21 | 0.00E+00 |
| REACTOME_SEPARATION_OF_SISTER_CHROMATIDS | 3.19 | 0.00E+00 |
| SIG_REGULATION_OF_THE_ACTIN_CYTOSKELETON_BY_RHO_GTPASES | 3.19 | 0.00E+00 |
| REACTOME_CHEMOKINE_RECEPTORS_BIND_CHEMOKINES | 3.17 | 0.00E+00 |
| WP_HEPATITIS_B_INFECTION | 3.17 | 1.23E-05 |
| HALLMARK_MITOTIC_SPINDLE | 3.17 | 1.22E-05 |
| REACTOME_INTRA_GOLGI_AND_RETROGRADE_GOLGI_TO_ER_TRAFFIC | 3.17 | 1.21E-05 |
| WP_MIRNA_TARGETS_IN_ECM_AND_MEMBRANE_RECEPTORS | 3.16 | 1.20E-05 |
| REACTOME_HOST_INTERACTIONS_OF_HIV_FACTORS | 3.16 | 1.19E-05 |
| NABA_SECRETED_FACTORS | 3.15 | 1.18E-05 |
| WP_NEURAL_CREST_CELL_MIGRATION_IN_CANCER | 3.15 | 1.17E-05 |
| WP_COMPLEMENT_SYSTEM | 3.14 | 1.16E-05 |
| REACTOME_GLYCOSAMINOGLYCAN_METABOLISM | 3.14 | 1.15E-05 |
| KEGG_PATHOGENIC_ESCHERICHIA_COLI_INFECTION | 3.13 | 1.14E-05 |
| PID_AURORA_B_PATHWAY | 3.13 | 1.14E-05 |
| WP_PATHOGENIC_ESCHERICHIA_COLI_INFECTION | 3.13 | 1.13E-05 |
| BIOCARTA_LAIR_PATHWAY | 3.12 | 1.12E-05 |
| KEGG_CELL_CYCLE | 3.11 | 1.11E-05 |
| WP_CELLS_AND_MOLECULES_INVOLVED_IN_LOCAL_ACUTE_INFLAMMATORY_RESPONSE | 3.11 | 1.10E-05 |
| REACTOME_SIGNALING_BY_MET | 3.11 | 1.09E-05 |
| WP_CELL_CYCLE | 3.11 | 1.08E-05 |
| WP_PI3KAKT_SIGNALING_PATHWAY | 3.10 | 1.08E-05 |
| WP_LUNG_FIBROSIS | 3.10 | 1.07E-05 |
| REACTOME_GOLGI_TO_ER_RETROGRADE_TRANSPORT | 3.09 | 1.06E-05 |
| HALLMARK_MYOGENESIS | 3.09 | 1.05E-05 |
| KEGG_P53_SIGNALING_PATHWAY | 3.08 | 1.04E-05 |
| KEGG_PATHWAYS_IN_CANCER | 3.07 | 1.04E-05 |
| WP_NEUROINFLAMMATION_AND_GLUTAMATERGIC_SIGNALING | 3.06 | 1.03E-05 |
| KEGG_LEISHMANIA_INFECTION | 3.04 | 1.02E-05 |
| REACTOME_COPI_DEPENDENT_GOLGI_TO_ER_RETROGRADE_TRAFFIC | 3.04 | 1.01E-05 |
| BIOCARTA_CELLCYCLE_PATHWAY | 3.04 | 1.01E-05 |
| REACTOME_INTERFERON_SIGNALING | 3.04 | 1.00E-05 |
| REACTOME_RRNA_PROCESSING | 3.04 | 9.93E-06 |
| HALLMARK_HYPOXIA | 3.03 | 9.87E-06 |
| REACTOME_GPVI_MEDIATED_ACTIVATION_CASCADE | 3.03 | 9.80E-06 |
| WP_G13_SIGNALING_PATHWAY | 3.03 | 9.73E-06 |
| WP_SMALL_CELL_LUNG_CANCER | 3.01 | 9.67E-06 |
| WP_INTEGRINMEDIATED_CELL_ADHESION | 3.01 | 9.60E-06 |
| REACTOME_SEMAPHORIN_INTERACTIONS | 3.00 | 9.54E-06 |
| PID_RAC1_REG_PATHWAY | 2.99 | 9.47E-06 |
| REACTOME_M_PHASE | 2.99 | 9.41E-06 |
| BIOCARTA_MONOCYTE_PATHWAY | 2.98 | 9.35E-06 |
| REACTOME_CROSSLINKING_OF_COLLAGEN_FIBRILS | 2.98 | 9.29E-06 |
| HALLMARK_COAGULATION | 2.96 | 9.23E-06 |
| PID_RAC1_PATHWAY | 2.95 | 9.17E-06 |
| WP_PROSTAGLANDIN_SIGNALING | 2.95 | 9.11E-06 |
| WP_INFLAMMATORY_RESPONSE_PATHWAY | 2.95 | 9.05E-06 |
| HALLMARK_CHOLESTEROL_HOMEOSTASIS | 2.95 | 9.00E-06 |
| KEGG_LYSOSOME | 2.94 | 8.94E-06 |
| REACTOME_HIV_INFECTION | 2.93 | 2.70E-05 |
| PID_INTEGRIN_A9B1_PATHWAY | 2.92 | 2.69E-05 |
| HALLMARK_PROTEIN_SECRETION | 2.91 | 2.67E-05 |
| HALLMARK_ESTROGEN_RESPONSE_EARLY | 2.90 | 2.65E-05 |
| REACTOME_RAC3_GTPASE_CYCLE | 2.90 | 2.64E-05 |
| REACTOME_KINESINS | 2.90 | 2.62E-05 |
| REACTOME_G2_M_CHECKPOINTS | 2.90 | 2.61E-05 |
| REACTOME_SRP_DEPENDENT_COTRANSLATIONAL_PROTEIN_TARGETING_TO_MEMBRANE | 2.89 | 3.47E-05 |
| REACTOME_CLEC7A_DECTIN_1_SIGNALING | 2.89 | 3.45E-05 |
| REACTOME_S_PHASE | 2.89 | 3.43E-05 |
| REACTOME_MET_PROMOTES_CELL_MOTILITY | 2.89 | 3.41E-05 |
| WP_SELECTIVE_EXPRESSION_OF_CHEMOKINE_RECEPTORS_DURING_TCELL_POLARIZATION | 2.88 | 3.39E-05 |
| REACTOME_MET_ACTIVATES_PTK2_SIGNALING | 2.87 | 3.37E-05 |
| REACTOME_SARS_COV_1_HOST_INTERACTIONS | 2.86 | 3.35E-05 |
| NABA_ECM_GLYCOPROTEINS | 2.86 | 3.33E-05 |
| WP_HEPATITIS_C_AND_HEPATOCELLULAR_CARCINOMA | 2.86 | 3.31E-05 |
| WP_PHOTODYNAMIC_THERAPYINDUCED_NFKB_SURVIVAL_SIGNALING | 2.85 | 3.30E-05 |
| WP_TOLLLIKE_RECEPTOR_SIGNALING_RELATED_TO_MYD88 | 2.85 | 4.10E-05 |
| WP_PANCREATIC_ADENOCARCINOMA_PATHWAY | 2.85 | 4.08E-05 |
| PID_ATF2_PATHWAY | 2.85 | 4.87E-05 |
| NABA_ECM_AFFILIATED | 2.84 | 4.85E-05 |
| WP_TGFBETA_SIGNALING_PATHWAY | 2.84 | 4.82E-05 |
| REACTOME_TNFR2_NON_CANONICAL_NF_KB_PATHWAY | 2.83 | 4.79E-05 |
| WP_DNA_DAMAGE_RESPONSE | 2.83 | 4.77E-05 |
| KEGG_T_CELL_RECEPTOR_SIGNALING_PATHWAY | 2.83 | 4.74E-05 |
| HALLMARK_TGF_BETA_SIGNALING | 2.81 | 7.08E-05 |
| REACTOME_EPHB_MEDIATED_FORWARD_SIGNALING | 2.81 | 7.04E-05 |
| REACTOME_C_TYPE_LECTIN_RECEPTORS_CLRS | 2.81 | 7.01E-05 |
| PID_TCR_PATHWAY | 2.81 | 8.54E-05 |
| REACTOME_DISSOLUTION_OF_FIBRIN_CLOT | 2.80 | 8.50E-05 |
| REACTOME_MAPK_FAMILY_SIGNALING_CASCADES | 2.80 | 8.45E-05 |
| REACTOME_SARS_COV_2_INFECTION | 2.79 | 9.92E-05 |
| KEGG_NATURAL_KILLER_CELL_MEDIATED_CYTOTOXICITY | 2.79 | 9.87E-05 |
| WP_MIRNA_REGULATION_OF_DNA_DAMAGE_RESPONSE | 2.79 | 9.82E-05 |
| BIOCARTA_NKT_PATHWAY | 2.79 | 9.77E-05 |
| REACTOME_EUKARYOTIC_TRANSLATION_INITIATION | 2.78 | 9.72E-05 |
| KEGG_CHRONIC_MYELOID_LEUKEMIA | 2.78 | 9.67E-05 |
| REACTOME_INTERFERON_ALPHA_BETA_SIGNALING | 2.77 | 1.11E-04 |
| BIOCARTA_G1_PATHWAY | 2.77 | 1.10E-04 |
| WP_EXTRAFOLLICULAR_AND_FOLLICULAR_B_CELL_ACTIVATION_BY_SARSCOV2 | 2.76 | 1.10E-04 |
| PID_AVB3_OPN_PATHWAY | 2.76 | 1.09E-04 |
| REACTOME_INTERLEUKIN_1_FAMILY_SIGNALING | 2.76 | 1.09E-04 |
| WP_MICROTUBULE_CYTOSKELETON_REGULATION | 2.76 | 1.08E-04 |
| PID_E2F_PATHWAY | 2.75 | 1.22E-04 |
| REACTOME_ABERRANT_REGULATION_OF_MITOTIC_G1_S_TRANSITION_IN_CANCER_DUE_TO_RB1_DEFECTS | 2.75 | 1.28E-04 |
| PID_ER_NONGENOMIC_PATHWAY | 2.75 | 1.28E-04 |
| BIOCARTA_ERYTH_PATHWAY | 2.73 | 1.55E-04 |
| WP_PROTEASOME_DEGRADATION | 2.73 | 1.62E-04 |
| WP_INTERACTIONS_BETWEEN_IMMUNE_CELLS_AND_MICRORNAS_IN_TUMOR_MICROENVIRONMENT | 2.73 | 1.68E-04 |
| REACTOME_DAP12_INTERACTIONS | 2.72 | 1.74E-04 |
| REACTOME_TCR_SIGNALING | 2.72 | 1.73E-04 |
| REACTOME_SCF_SKP2_MEDIATED_DEGRADATION_OF_P27_P21 | 2.71 | 2.00E-04 |
| REACTOME_TOLL_LIKE_RECEPTOR_CASCADES | 2.71 | 1.99E-04 |
| KEGG_EPITHELIAL_CELL_SIGNALING_IN_HELICOBACTER_PYLORI_INFECTION | 2.71 | 2.12E-04 |
| PID_INTEGRIN_CS_PATHWAY | 2.70 | 2.18E-04 |
| REACTOME_PROGRAMMED_CELL_DEATH | 2.69 | 2.31E-04 |
| PID_FOXM1_PATHWAY | 2.69 | 2.43E-04 |
| KEGG_JAK_STAT_SIGNALING_PATHWAY | 2.69 | 2.49E-04 |
| REACTOME_DOWNSTREAM_SIGNALING_EVENTS_OF_B_CELL_RECEPTOR_BCR | 2.69 | 2.47E-04 |
| REACTOME_SIGNALING_BY_GPCR | 2.69 | 2.46E-04 |
| REACTOME_INFLAMMASOMES | 2.69 | 2.45E-04 |
| REACTOME_RHOA_GTPASE_CYCLE | 2.68 | 2.71E-04 |
| WP_NONSMALL_CELL_LUNG_CANCER | 2.67 | 2.69E-04 |
| REACTOME_CLATHRIN_MEDIATED_ENDOCYTOSIS | 2.67 | 2.81E-04 |
| REACTOME_PURINERGIC_SIGNALING_IN_LEISHMANIASIS_INFECTION | 2.67 | 2.93E-04 |
| WP_PHOTODYNAMIC_THERAPYINDUCED_AP1_SURVIVAL_SIGNALING | 2.66 | 3.11E-04 |
| REACTOME_MITOTIC_SPINDLE_CHECKPOINT | 2.66 | 3.16E-04 |
| WP_MIRNA_ROLE_IN_IMMUNE_RESPONSE_IN_SEPSIS | 2.66 | 3.15E-04 |
| REACTOME_CYCLIN_D_ASSOCIATED_EVENTS_IN_G1 | 2.65 | 3.32E-04 |
| REACTOME_SIGNALING_BY_VEGF | 2.65 | 3.31E-04 |
| REACTOME_SYNTHESIS_OF_DNA | 2.65 | 3.48E-04 |
| REACTOME_DNA_REPLICATION | 2.65 | 3.53E-04 |
| PID_P38_ALPHA_BETA_PATHWAY | 2.65 | 3.52E-04 |
| REACTOME_SPHINGOLIPID_METABOLISM | 2.65 | 3.50E-04 |
| HALLMARK_MYC_TARGETS_V2 | 2.65 | 3.49E-04 |
| WP_MAMMARY_GLAND_DEVELOPMENT_PATHWAY_PUBERTY_STAGE_2_OF_4 | 2.64 | 3.47E-04 |
| BIOCARTA_PLATELETAPP_PATHWAY | 2.64 | 3.52E-04 |
| WP_ARYL_HYDROCARBON_RECEPTOR_PATHWAY_WP2586 | 2.64 | 3.51E-04 |
| REACTOME_APC_C_MEDIATED_DEGRADATION_OF_CELL_CYCLE_PROTEINS | 2.64 | 3.55E-04 |
| REACTOME_INTERLEUKIN_12_FAMILY_SIGNALING | 2.63 | 3.72E-04 |
| WP_TGFBETA_RECEPTOR_SIGNALING_IN_SKELETAL_DYSPLASIAS | 2.63 | 3.71E-04 |
| BIOCARTA_LYMPHOCYTE_PATHWAY | 2.63 | 3.81E-04 |
| REACTOME_REGULATION_OF_INSULIN_LIKE_GROWTH_FACTOR_IGF_TRANSPORT_AND_UPTAKE_BY_INSULIN_LIKE_GROWTH_FACTOR_BINDING_PROTEINS_IGFBPS | 2.63 | 3.92E-04 |
| WP_VITAMIN_D_RECEPTOR_PATHWAY | 2.62 | 4.02E-04 |
| REACTOME_REGULATION_OF_RUNX3_EXPRESSION_AND_ACTIVITY | 2.62 | 4.36E-04 |
| REACTOME_SARS_COV_1_INFECTION | 2.61 | 4.76E-04 |
| PID_GLYPICAN_1PATHWAY | 2.61 | 4.98E-04 |
| WP_TYPE_II_INTERFERON_SIGNALING | 2.61 | 5.08E-04 |
| BIOCARTA_BLYMPHOCYTE_PATHWAY | 2.61 | 5.06E-04 |
| REACTOME_PCP_CE_PATHWAY | 2.60 | 5.16E-04 |
| PID_ERBB1_DOWNSTREAM_PATHWAY | 2.60 | 5.14E-04 |
| PID_P53_DOWNSTREAM_PATHWAY | 2.60 | 5.30E-04 |
| REACTOME_GENE_AND_PROTEIN_EXPRESSION_BY_JAK_STAT_SIGNALING_AFTER_INTERLEUKIN_12_STIMULATION | 2.59 | 5.62E-04 |
| REACTOME_MITOTIC_PROMETAPHASE | 2.59 | 5.71E-04 |
| REACTOME_ORC1_REMOVAL_FROM_CHROMATIN | 2.59 | 5.86E-04 |
| KEGG_HYPERTROPHIC_CARDIOMYOPATHY_HCM | 2.59 | 5.84E-04 |
| PID_PI3KCI_PATHWAY | 2.58 | 6.28E-04 |
| REACTOME_DECTIN_1_MEDIATED_NONCANONICAL_NF_KB_SIGNALING | 2.58 | 6.31E-04 |
| REACTOME_SIGNALING_BY_PDGF | 2.58 | 6.29E-04 |
| WP_SPHINGOLIPID_METABOLISM_INTEGRATED_PATHWAY | 2.57 | 6.43E-04 |
| WP_HIPPOMERLIN_SIGNALING_DYSREGULATION | 2.57 | 6.52E-04 |
| REACTOME_SIGNALING_BY_THE_B_CELL_RECEPTOR_BCR | 2.57 | 6.66E-04 |
| WP_DEVELOPMENT_OF_PULMONARY_DENDRITIC_CELLS_AND_MACROPHAGE_SUBSETS | 2.56 | 6.92E-04 |
| PID_NFAT_TFPATHWAY | 2.56 | 7.39E-04 |
| REACTOME_DEATH_RECEPTOR_SIGNALING | 2.56 | 7.36E-04 |
| KEGG_GLIOMA | 2.55 | 7.72E-04 |
| REACTOME_NCAM1_INTERACTIONS | 2.55 | 7.91E-04 |
| PID_INTEGRIN5_PATHWAY | 2.55 | 8.05E-04 |
| REACTOME_DISEASES_OF_IMMUNE_SYSTEM | 2.55 | 8.13E-04 |
| HALLMARK_APICAL_SURFACE | 2.54 | 9.02E-04 |
| REACTOME_THE_ROLE_OF_GTSE1_IN_G2_M_PROGRESSION_AFTER_G2_CHECKPOINT | 2.54 | 8.99E-04 |
| REACTOME_SARS_COV_2_HOST_INTERACTIONS | 2.53 | 9.12E-04 |
| WP_PROSTAGLANDIN_SYNTHESIS_AND_REGULATION | 2.53 | 9.09E-04 |
| REACTOME_RRNA_MODIFICATION_IN_THE_NUCLEUS_AND_CYTOSOL | 2.53 | 9.06E-04 |
| BIOCARTA_P53_PATHWAY | 2.53 | 9.29E-04 |
| REACTOME_RHO_GTPASES_ACTIVATE_WASPS_AND_WAVES | 2.53 | 9.37E-04 |
| WP_MAMMARY_GLAND_DEVELOPMENT_PATHWAY_INVOLUTION_STAGE_4_OF_4 | 2.53 | 9.44E-04 |
| REACTOME_METABOLISM_OF_FAT_SOLUBLE_VITAMINS | 2.53 | 9.46E-04 |
| WP_OXIDATIVE_DAMAGE_RESPONSE | 2.52 | 9.48E-04 |
| PID_FCER1_PATHWAY | 2.52 | 1.01E-03 |
| WP_SARSCOV2_INNATE_IMMUNITY_EVASION_AND_CELLSPECIFIC_IMMUNE_RESPONSE | 2.52 | 1.01E-03 |
| REACTOME_FC_EPSILON_RECEPTOR_FCERI_SIGNALING | 2.51 | 1.08E-03 |
| REACTOME_TRANSCRIPTIONAL_REGULATION_BY_RUNX3 | 2.51 | 1.08E-03 |
| PID_NECTIN_PATHWAY | 2.50 | 1.10E-03 |
| REACTOME_BETA_CATENIN_INDEPENDENT_WNT_SIGNALING | 2.50 | 1.11E-03 |
| WP_TGFBETA_RECEPTOR_SIGNALING | 2.50 | 1.15E-03 |
| HALLMARK_ESTROGEN_RESPONSE_LATE | 2.50 | 1.16E-03 |
| WP_RAS_SIGNALING | 2.49 | 1.20E-03 |
| PID_IL8_CXCR2_PATHWAY | 2.48 | 1.29E-03 |
| REACTOME_SYNTHESIS_OF_UDP_N_ACETYL_GLUCOSAMINE | 2.48 | 1.29E-03 |
| REACTOME_CDC42_GTPASE_CYCLE | 2.48 | 1.29E-03 |
| REACTOME_KERATAN_SULFATE_KERATIN_METABOLISM | 2.47 | 1.31E-03 |
| HALLMARK_UV_RESPONSE_DN | 2.47 | 1.32E-03 |
| WP_APOPTOSISRELATED_NETWORK_DUE_TO_ALTERED_NOTCH3_IN_OVARIAN_CANCER | 2.46 | 1.42E-03 |
| REACTOME_INTERLEUKIN_12_SIGNALING | 2.46 | 1.42E-03 |
| KEGG_AMINO_SUGAR_AND_NUCLEOTIDE_SUGAR_METABOLISM | 2.46 | 1.42E-03 |
| WP_SPHINGOLIPID_METABOLISM_IN_SENESCENCE | 2.46 | 1.45E-03 |
| KEGG_B_CELL_RECEPTOR_SIGNALING_PATHWAY | 2.46 | 1.49E-03 |
| BIOCARTA_LYM_PATHWAY | 2.46 | 1.50E-03 |
| PID_FRA_PATHWAY | 2.45 | 1.58E-03 |
| WP_MACROPHAGE_MARKERS | 2.45 | 1.60E-03 |
| REACTOME_SARS_COV_1_ACTIVATES_MODULATES_INNATE_IMMUNE_RESPONSES | 2.45 | 1.66E-03 |
| PID_DELTA_NP63_PATHWAY | 2.44 | 1.69E-03 |
| PID_SYNDECAN_2_PATHWAY | 2.44 | 1.73E-03 |
| REACTOME_SIGNALING_BY_EGFR | 2.44 | 1.72E-03 |
| REACTOME_HEDGEHOG_ON_STATE | 2.43 | 1.75E-03 |
| REACTOME_INFLUENZA_INFECTION | 2.43 | 1.75E-03 |
| KEGG_PROSTATE_CANCER | 2.43 | 1.74E-03 |
| WP_OVERVIEW_OF_NANOPARTICLE_EFFECTS | 2.43 | 1.74E-03 |
| REACTOME_THE_ROLE_OF_NEF_IN_HIV_1_REPLICATION_AND_DISEASE_PATHOGENESIS | 2.43 | 1.73E-03 |
| WP_TCELL_RECEPTOR_SIGNALING_PATHWAY | 2.43 | 1.74E-03 |
| REACTOME_SIGNALING_BY_HEDGEHOG | 2.43 | 1.75E-03 |
| REACTOME_SIGNALING_BY_CSF1_M_CSF_IN_MYELOID_CELLS | 2.43 | 1.76E-03 |
| REACTOME_DISEASES_OF_GLYCOSYLATION | 2.43 | 1.77E-03 |
| REACTOME_REGULATION_OF_MRNA_STABILITY_BY_PROTEINS_THAT_BIND_AU_RICH_ELEMENTS | 2.42 | 1.86E-03 |
| REACTOME_EGFR_DOWNREGULATION | 2.42 | 1.85E-03 |
| REACTOME_P75_NTR_RECEPTOR_MEDIATED_SIGNALLING | 2.42 | 1.84E-03 |
| REACTOME_CROSS_PRESENTATION_OF_SOLUBLE_EXOGENOUS_ANTIGENS_ENDOSOMES | 2.42 | 1.94E-03 |
| PID_INTEGRIN2_PATHWAY | 2.42 | 1.94E-03 |
| PID_IL23_PATHWAY | 2.41 | 1.95E-03 |
| BIOCARTA_NEUTROPHIL_PATHWAY | 2.41 | 1.98E-03 |
| WP_SPHINGOLIPID_PATHWAY | 2.41 | 1.98E-03 |
| WP_FOCAL_ADHESION_PI3KAKTMTORSIGNALING_PATHWAY | 2.41 | 2.00E-03 |
| WP_T_CELL_MODULATION_IN_PANCREATIC_CANCER | 2.41 | 2.00E-03 |
| KEGG_ADHERENS_JUNCTION | 2.41 | 2.02E-03 |
| WP_REGULATORY_CIRCUITS_OF_THE_STAT3_SIGNALING_PATHWAY | 2.41 | 2.06E-03 |
| WP_EGFEGFR_SIGNALING_PATHWAY | 2.40 | 2.14E-03 |
| REACTOME_DISEASES_OF_SIGNAL_TRANSDUCTION_BY_GROWTH_FACTOR_RECEPTORS_AND_SECOND_MESSENGERS | 2.40 | 2.14E-03 |
| REACTOME_CLASS_A_1_RHODOPSIN_LIKE_RECEPTORS | 2.40 | 2.15E-03 |
| WP_AGERAGE_PATHWAY | 2.40 | 2.16E-03 |
| REACTOME_ANTIVIRAL_MECHANISM_BY_IFN_STIMULATED_GENES | 2.39 | 2.20E-03 |
| REACTOME_VISUAL_PHOTOTRANSDUCTION | 2.39 | 2.27E-03 |
| BIOCARTA_GRANULOCYTES_PATHWAY | 2.39 | 2.28E-03 |
| REACTOME_RHOG_GTPASE_CYCLE | 2.39 | 2.31E-03 |
| REACTOME_ROS_AND_RNS_PRODUCTION_IN_PHAGOCYTES | 2.39 | 2.32E-03 |
| REACTOME_INTERLEUKIN_1_PROCESSING | 2.38 | 2.43E-03 |
| REACTOME_DEGRADATION_OF_AXIN | 2.38 | 2.44E-03 |
| KEGG_N_GLYCAN_BIOSYNTHESIS | 2.38 | 2.44E-03 |
| PID_CD8_TCR_PATHWAY | 2.38 | 2.44E-03 |
| REACTOME_EPH_EPHRIN_MEDIATED_REPULSION_OF_CELLS | 2.37 | 2.64E-03 |
| KEGG_PRION_DISEASES | 2.37 | 2.65E-03 |
| REACTOME_AUF1_HNRNP_D0_BINDS_AND_DESTABILIZES_MRNA | 2.37 | 2.65E-03 |
| REACTOME_DNA_REPLICATION_PRE_INITIATION | 2.37 | 2.65E-03 |
| REACTOME_NONSENSE_MEDIATED_DECAY_NMD | 2.36 | 2.76E-03 |
| REACTOME_DRUG_MEDIATED_INHIBITION_OF_CDK4_CDK6_ACTIVITY | 2.36 | 2.76E-03 |
| WP_FAS_LIGAND_PATHWAY_AND_STRESS_INDUCTION_OF_HEAT_SHOCK_PROTEINS | 2.36 | 2.79E-03 |
| REACTOME_APOPTOSIS | 2.36 | 2.79E-03 |
| PID_PLK1_PATHWAY | 2.36 | 2.80E-03 |
| REACTOME_DECTIN_2_FAMILY | 2.36 | 2.80E-03 |
| KEGG_SPHINGOLIPID_METABOLISM | 2.35 | 2.88E-03 |
| REACTOME_EUKARYOTIC_TRANSLATION_ELONGATION | 2.35 | 2.91E-03 |
| PID_EPHA2_FWD_PATHWAY | 2.35 | 2.90E-03 |
| WP_GLIOBLASTOMA_SIGNALING_PATHWAYS | 2.35 | 2.97E-03 |
| REACTOME_DAP12_SIGNALING | 2.35 | 2.96E-03 |
| BIOCARTA_FIBRINOLYSIS_PATHWAY | 2.35 | 2.96E-03 |
| REACTOME_CROSS_PRESENTATION_OF_PARTICULATE_EXOGENOUS_ANTIGENS_PHAGOSOMES | 2.35 | 3.04E-03 |
| REACTOME_PEPTIDE_LIGAND_BINDING_RECEPTORS | 2.34 | 3.06E-03 |
| REACTOME_CYCLIN_A_CDK2_ASSOCIATED_EVENTS_AT_S_PHASE_ENTRY | 2.34 | 3.13E-03 |
| KEGG_MAPK_SIGNALING_PATHWAY | 2.34 | 3.14E-03 |
| REACTOME_ACTIVATION_OF_RAC1_DOWNSTREAM_OF_NMDARS | 2.34 | 3.14E-03 |
| REACTOME_SWITCHING_OF_ORIGINS_TO_A_POST_REPLICATIVE_STATE | 2.34 | 3.17E-03 |
| PID_IL12_2PATHWAY | 2.34 | 3.16E-03 |
| REACTOME_SIGNALING_BY_TGF_BETA_RECEPTOR_COMPLEX | 2.34 | 3.16E-03 |
| REACTOME_LAMININ_INTERACTIONS | 2.34 | 3.19E-03 |
| REACTOME_SCF_BETA_TRCP_MEDIATED_DEGRADATION_OF_EMI1 | 2.34 | 3.18E-03 |
| PID_THROMBIN_PAR1_PATHWAY | 2.34 | 3.19E-03 |
| WP_B_CELL_RECEPTOR_SIGNALING_PATHWAY | 2.33 | 3.26E-03 |
| REACTOME_SIGNALING_BY_TGFB_FAMILY_MEMBERS | 2.33 | 3.30E-03 |
| PID_CMYB_PATHWAY | 2.33 | 3.34E-03 |
| SIG_CHEMOTAXIS | 2.33 | 3.36E-03 |
| REACTOME_HEDGEHOG_LIGAND_BIOGENESIS | 2.32 | 3.40E-03 |
| REACTOME_INTERLEUKIN_1_SIGNALING | 2.32 | 3.40E-03 |
| WP_NUCLEAR_RECEPTORS_METAPATHWAY | 2.32 | 3.41E-03 |
| REACTOME_MAPK6_MAPK4_SIGNALING | 2.32 | 3.60E-03 |
| REACTOME_ANTIMICROBIAL_PEPTIDES | 2.31 | 3.64E-03 |
| PID_ECADHERIN_NASCENT_AJ_PATHWAY | 2.31 | 3.64E-03 |
| REACTOME_FCERI_MEDIATED_NF_KB_ACTIVATION | 2.31 | 3.81E-03 |
| WP_INFLAMMATORY_BOWEL_DISEASE_SIGNALING | 2.31 | 3.82E-03 |
| WP_GENES_ASSOCIATED_WITH_THE_DEVELOPMENT_OF_RHEUMATOID_ARTHRITIS | 2.31 | 3.81E-03 |
| REACTOME_POST_CHAPERONIN_TUBULIN_FOLDING_PATHWAY | 2.31 | 3.82E-03 |
| REACTOME_SARS_COV_2_ACTIVATES_MODULATES_INNATE_AND_ADAPTIVE_IMMUNE_RESPONSES | 2.30 | 3.89E-03 |
| REACTOME_SIGNALING_BY_NTRKS | 2.30 | 3.90E-03 |
| WP_UREA_CYCLE_AND_METABOLISM_OF_AMINO_GROUPS | 2.30 | 4.00E-03 |
| BIOCARTA_NKCELLS_PATHWAY | 2.29 | 4.07E-03 |
| BIOCARTA_MSP_PATHWAY | 2.29 | 4.11E-03 |
| REACTOME_METABOLISM_OF_CARBOHYDRATES | 2.29 | 4.10E-03 |
| WP_SELENIUM_MICRONUTRIENT_NETWORK | 2.29 | 4.16E-03 |
| PID_CDC42_PATHWAY | 2.29 | 4.26E-03 |
| REACTOME_CELL_CELL_COMMUNICATION | 2.29 | 4.26E-03 |
| REACTOME_CHONDROITIN_SULFATE_DERMATAN_SULFATE_METABOLISM | 2.28 | 4.26E-03 |
| PID_ERBB1_INTERNALIZATION_PATHWAY | 2.28 | 4.26E-03 |
| WP_ARYL_HYDROCARBON_RECEPTOR_PATHWAY_WP2873 | 2.28 | 4.27E-03 |
| HALLMARK_GLYCOLYSIS | 2.28 | 4.44E-03 |
| KEGG_SMALL_CELL_LUNG_CANCER | 2.28 | 4.51E-03 |
| REACTOME_TRANSPORT_TO_THE_GOLGI_AND_SUBSEQUENT_MODIFICATION | 2.28 | 4.56E-03 |
| REACTOME_RHOU_GTPASE_CYCLE | 2.28 | 4.59E-03 |
| REACTOME_ONCOGENIC_MAPK_SIGNALING | 2.27 | 4.65E-03 |
| WP_LTF_DANGER_SIGNAL_RESPONSE_PATHWAY | 2.27 | 4.65E-03 |
| REACTOME_P130CAS_LINKAGE_TO_MAPK_SIGNALING_FOR_INTEGRINS | 2.26 | 5.00E-03 |
| REACTOME_THE_NLRP3_INFLAMMASOME | 2.26 | 5.00E-03 |
| REACTOME_INTERLEUKIN_RECEPTOR_SHC_SIGNALING | 2.26 | 5.21E-03 |
| REACTOME_DEGRADATION_OF_BETA_CATENIN_BY_THE_DESTRUCTION_COMPLEX | 2.26 | 5.23E-03 |
| BIOCARTA_HIVNEF_PATHWAY | 2.25 | 5.26E-03 |
| KEGG_O_GLYCAN_BIOSYNTHESIS | 2.25 | 5.33E-03 |
| KEGG_NON_SMALL_CELL_LUNG_CANCER | 2.25 | 5.38E-03 |
| WP_CYTOPLASMIC_RIBOSOMAL_PROTEINS | 2.25 | 5.40E-03 |
| REACTOME_BASIGIN_INTERACTIONS | 2.25 | 5.40E-03 |
| REACTOME_TRANS_GOLGI_NETWORK_VESICLE_BUDDING | 2.24 | 5.54E-03 |
| REACTOME_GENERATION_OF_SECOND_MESSENGER_MOLECULES | 2.24 | 5.55E-03 |
| REACTOME_DEFECTIVE_CFTR_CAUSES_CYSTIC_FIBROSIS | 2.24 | 5.71E-03 |
| REACTOME_RHO_GTPASES_ACTIVATE_IQGAPS | 2.23 | 5.85E-03 |
| HALLMARK_ANGIOGENESIS | 2.23 | 5.84E-03 |
| WP_PROLACTIN_SIGNALING_PATHWAY | 2.23 | 5.83E-03 |
| REACTOME_DISEASES_ASSOCIATED_WITH_GLYCOSAMINOGLYCAN_METABOLISM | 2.23 | 5.84E-03 |
| WP_MYD88_DISTINCT_INPUTOUTPUT_PATHWAY | 2.23 | 5.89E-03 |
| BIOCARTA_BTG2_PATHWAY | 2.23 | 6.08E-03 |
| WP_SYNTHESIS_OF_CERAMIDES_AND_1DEOXYCERAMIDES | 2.23 | 6.15E-03 |
| WP_FGF23_SIGNALING_IN_HYPOPHOSPHATEMIC_RICKETS_AND_RELATED_DISORDERS | 2.22 | 6.49E-03 |
| REACTOME_DETOXIFICATION_OF_REACTIVE_OXYGEN_SPECIES | 2.22 | 6.54E-03 |
| WP_TNFRELATED_WEAK_INDUCER_OF_APOPTOSIS_TWEAK_SIGNALING_PATHWAY | 2.22 | 6.57E-03 |
| REACTOME_REGULATION_OF_RAS_BY_GAPS | 2.21 | 6.67E-03 |
| WP_CELL_INTERACTIONS_OF_THE_PANCREATIC_CANCER_MICROENVIRONMENT | 2.21 | 6.68E-03 |
| REACTOME_RESPONSE_OF_EIF2AK4_GCN2_TO_AMINO_ACID_DEFICIENCY | 2.21 | 6.71E-03 |
| PID_AJDISS_2PATHWAY | 2.21 | 6.84E-03 |
| WP_TCELL_ANTIGEN_RECEPTOR_TCR_PATHWAY_DURING_STAPHYLOCOCCUS_AUREUS_INFECTION | 2.21 | 6.85E-03 |
| WP_PHYSICOCHEMICAL_FEATURES_AND_TOXICITYASSOCIATED_PATHWAYS | 2.21 | 6.86E-03 |
| WP_SPHINGOLIPID_METABOLISM_OVERVIEW | 2.21 | 6.92E-03 |
| REACTOME_CARGO_RECOGNITION_FOR_CLATHRIN_MEDIATED_ENDOCYTOSIS | 2.21 | 6.95E-03 |
| REACTOME_INTERLEUKIN_3_INTERLEUKIN_5_AND_GM_CSF_SIGNALING | 2.20 | 6.99E-03 |
| REACTOME_WNT5A_DEPENDENT_INTERNALIZATION_OF_FZD2_FZD5_AND_ROR2 | 2.20 | 6.99E-03 |
| REACTOME_RECYCLING_PATHWAY_OF_L1 | 2.20 | 7.15E-03 |
| WP_CCL18_SIGNALING_PATHWAY | 2.20 | 7.14E-03 |
| WP_HEMATOPOIETIC_STEM_CELL_DIFFERENTIATION | 2.20 | 7.15E-03 |
| WP_CHOLESTEROL_BIOSYNTHESIS_PATHWAY | 2.20 | 7.21E-03 |
| WP_MAPK_SIGNALING_PATHWAY | 2.20 | 7.21E-03 |
| KEGG_FC_EPSILON_RI_SIGNALING_PATHWAY | 2.19 | 7.44E-03 |
| KEGG_PROTEASOME | 2.19 | 7.56E-03 |
| WP_HOSTPATHOGEN_INTERACTION_OF_HUMAN_CORONAVIRUSES_INTERFERON_INDUCTION | 2.19 | 7.58E-03 |
| REACTOME_UCH_PROTEINASES | 2.19 | 7.58E-03 |
| REACTOME_BINDING_AND_UPTAKE_OF_LIGANDS_BY_SCAVENGER_RECEPTORS | 2.19 | 7.60E-03 |
| REACTOME_G0_AND_EARLY_G1 | 2.18 | 7.71E-03 |
| REACTOME_G1_S_DNA_DAMAGE_CHECKPOINTS | 2.18 | 7.81E-03 |
| HALLMARK_UV_RESPONSE_UP | 2.18 | 7.80E-03 |
| WP_TYPE_I_INTERFERON_INDUCTION_AND_SIGNALING_DURING_SARSCOV2_INFECTION | 2.18 | 7.79E-03 |
| REACTOME_SCAVENGING_BY_CLASS_A_RECEPTORS | 2.18 | 7.87E-03 |
| REACTOME_NEF_MEDIATED_CD8_DOWN_REGULATION | 2.18 | 7.86E-03 |
| REACTOME_TP53_REGULATES_TRANSCRIPTION_OF_CELL_CYCLE_GENES | 2.18 | 7.87E-03 |
| REACTOME_THE_CANONICAL_RETINOID_CYCLE_IN_RODS_TWILIGHT_VISION | 2.18 | 7.86E-03 |
| REACTOME_METABOLISM_OF_POLYAMINES | 2.18 | 7.84E-03 |
| SA_REG_CASCADE_OF_CYCLIN_EXPR | 2.18 | 7.98E-03 |
| WP_MIRNA_REGULATION_OF_P53_PATHWAY_IN_PROSTATE_CANCER | 2.18 | 8.01E-03 |
| REACTOME_SIGNALING_BY_PTK6 | 2.17 | 8.12E-03 |
| PID_ARF6_TRAFFICKING_PATHWAY | 2.17 | 8.17E-03 |
| WP_STRIATED_MUSCLE_CONTRACTION_PATHWAY | 2.17 | 8.16E-03 |
| WP_CHOLESTEROL_SYNTHESIS_DISORDERS | 2.17 | 8.25E-03 |
| REACTOME_DEUBIQUITINATION | 2.17 | 8.24E-03 |
| REACTOME_INTERLEUKIN_2_FAMILY_SIGNALING | 2.17 | 8.29E-03 |
| REACTOME_DEGRADATION_OF_GLI1_BY_THE_PROTEASOME | 2.17 | 8.32E-03 |
| REACTOME_POTENTIAL_THERAPEUTICS_FOR_SARS | 2.17 | 8.30E-03 |
| BIOCARTA_CCR5_PATHWAY | 2.17 | 8.28E-03 |
| REACTOME_ACTIVATION_OF_NIMA_KINASES_NEK9_NEK6_NEK7 | 2.17 | 8.38E-03 |
| WP_EGFR_TYROSINE_KINASE_INHIBITOR_RESISTANCE | 2.16 | 8.48E-03 |
| KEGG_VIBRIO_CHOLERAE_INFECTION | 2.16 | 8.50E-03 |
| REACTOME_ACTIVATION_OF_AMPK_DOWNSTREAM_OF_NMDARS | 2.16 | 8.51E-03 |
| PID_TXA2PATHWAY | 2.16 | 8.57E-03 |
| PID_FAK_PATHWAY | 2.16 | 8.67E-03 |
| REACTOME_UB_SPECIFIC_PROCESSING_PROTEASES | 2.16 | 8.68E-03 |
| WP_IMMUNE_RESPONSE_TO_TUBERCULOSIS | 2.16 | 8.70E-03 |
| HALLMARK_UNFOLDED_PROTEIN_RESPONSE | 2.16 | 8.75E-03 |
| REACTOME_NUCLEAR_EVENTS_MEDIATED_BY_NFE2L2 | 2.16 | 8.75E-03 |
| REACTOME_NUCLEAR_ENVELOPE_NE_REASSEMBLY | 2.15 | 9.01E-03 |
| REACTOME_RHO_GTPASES_ACTIVATE_PAKS | 2.15 | 9.08E-03 |
| WP_GENETIC_CAUSES_OF_PORTOSINUSOIDAL_VASCULAR_DISEASE | 2.15 | 9.30E-03 |
| REACTOME_IRAK4_DEFICIENCY_TLR2_4 | 2.15 | 9.35E-03 |
| KEGG_MELANOMA | 2.14 | 9.54E-03 |
| PID_RB_1PATHWAY | 2.14 | 9.52E-03 |
| WP_PROSURVIVAL_SIGNALING_OF_NEUROPROTECTIN_D1 | 2.14 | 9.54E-03 |
| REACTOME_SIGNALING_BY_NOTCH4 | 2.14 | 9.60E-03 |
| REACTOME_HIV_LIFE_CYCLE | 2.14 | 9.60E-03 |
| REACTOME_NEGATIVE_REGULATION_OF_NOTCH4_SIGNALING | 2.14 | 9.74E-03 |
| REACTOME_EXTRA_NUCLEAR_ESTROGEN_SIGNALING | 2.14 | 9.74E-03 |
| REACTOME_SIGNALING_BY_HIPPO | 2.14 | 9.82E-03 |
| REACTOME_MITOTIC_G2_G2_M_PHASES | 2.13 | 9.88E-03 |
| REACTOME_SIGNALING_BY_WNT | 2.13 | 9.93E-03 |
| WP_ALLOGRAFT_REJECTION | 2.13 | 1.03E-02 |
| WP_BLADDER_CANCER | 2.13 | 1.03E-02 |
| BIOCARTA_NPP1_PATHWAY | 2.12 | 1.05E-02 |
| PID_AP1_PATHWAY | 2.12 | 1.05E-02 |
| REACTOME_DISEASES_OF_MITOTIC_CELL_CYCLE | 2.12 | 1.06E-02 |
| REACTOME_APC_C_CDH1_MEDIATED_DEGRADATION_OF_CDC20_AND_OTHER_APC_C_CDH1_TARGETED_PROTEINS_IN_LATE_MITOSIS_EARLY_G1 | 2.12 | 1.07E-02 |
| WP_IMMUNE_INFILTRATION_IN_PANCREATIC_CANCER | 2.12 | 1.07E-02 |
| WP_P53_TRANSCRIPTIONAL_GENE_NETWORK | 2.12 | 1.07E-02 |
| REACTOME_RUNX1_REGULATES_TRANSCRIPTION_OF_GENES_INVOLVED_IN_DIFFERENTIATION_OF_KERATINOCYTES | 2.12 | 1.08E-02 |
| BIOCARTA_VITCB_PATHWAY | 2.12 | 1.08E-02 |
| PID_PTP1B_PATHWAY | 2.12 | 1.10E-02 |
| REACTOME_ASYMMETRIC_LOCALIZATION_OF_PCP_PROTEINS | 2.12 | 1.10E-02 |
| BIOCARTA_HSP27_PATHWAY | 2.11 | 1.10E-02 |
| WP_NRF2_PATHWAY | 2.11 | 1.10E-02 |
| PID_ECADHERIN_STABILIZATION_PATHWAY | 2.11 | 1.14E-02 |
| KEGG_PANCREATIC_CANCER | 2.11 | 1.14E-02 |
| WP_2Q13_COPY_NUMBER_VARIATION_SYNDROME | 2.11 | 1.14E-02 |
| REACTOME_TRANSLOCATION_OF_SLC2A4_GLUT4_TO_THE_PLASMA_MEMBRANE | 2.11 | 1.16E-02 |
| PID_FGF_PATHWAY | 2.11 | 1.16E-02 |
| BIOCARTA_INTRINSIC_PATHWAY | 2.10 | 1.19E-02 |
| REACTOME_SUMOYLATION_OF_DNA_REPLICATION_PROTEINS | 2.10 | 1.20E-02 |
| KEGG_RIBOSOME | 2.10 | 1.21E-02 |
| KEGG_ANTIGEN_PROCESSING_AND_PRESENTATION | 2.09 | 1.23E-02 |
| WP_NCRNAS_INVOLVED_IN_STAT3_SIGNALING_IN_HEPATOCELLULAR_CARCINOMA | 2.09 | 1.27E-02 |
| WP_ACUTE_VIRAL_MYOCARDITIS | 2.09 | 1.29E-02 |
| KEGG_STEROID_BIOSYNTHESIS | 2.09 | 1.30E-02 |
| REACTOME_RHOF_GTPASE_CYCLE | 2.08 | 1.32E-02 |
| REACTOME_CELL_DEATH_SIGNALLING_VIA_NRAGE_NRIF_AND_NADE | 2.08 | 1.32E-02 |
| REACTOME_DEX_H_BOX_HELICASES_ACTIVATE_TYPE_I_IFN_AND_INFLAMMATORY_CYTOKINES_PRODUCTION | 2.08 | 1.32E-02 |
| REACTOME_REGULATION_OF_SIGNALING_BY_CBL | 2.08 | 1.32E-02 |
| REACTOME_GLYCOSPHINGOLIPID_METABOLISM | 2.08 | 1.32E-02 |
| WP_MIRNAS_INVOLVED_IN_DNA_DAMAGE_RESPONSE | 2.08 | 1.35E-02 |
| WP_PLURIPOTENT_STEM_CELL_DIFFERENTIATION_PATHWAY | 2.07 | 1.36E-02 |
| KEGG_CYTOSOLIC_DNA_SENSING_PATHWAY | 2.07 | 1.38E-02 |
| REACTOME_HOMOLOGY_DIRECTED_REPAIR | 2.07 | 1.39E-02 |
| PID_ARF_3PATHWAY | 2.07 | 1.39E-02 |
| WP_APOPTOSIS | 2.07 | 1.39E-02 |
| REACTOME_NGF_STIMULATED_TRANSCRIPTION | 2.07 | 1.39E-02 |
| PID_TOLL_ENDOGENOUS_PATHWAY | 2.07 | 1.39E-02 |
| REACTOME_RHOC_GTPASE_CYCLE | 2.07 | 1.39E-02 |
| REACTOME_NRAGE_SIGNALS_DEATH_THROUGH_JNK | 2.07 | 1.40E-02 |
| BIOCARTA_UCALPAIN_PATHWAY | 2.07 | 1.40E-02 |
| KEGG_COMPLEMENT_AND_COAGULATION_CASCADES | 2.07 | 1.40E-02 |
| REACTOME_TRIF_MEDIATED_PROGRAMMED_CELL_DEATH | 2.07 | 1.42E-02 |
| REACTOME_G_ALPHA_I_SIGNALLING_EVENTS | 2.07 | 1.42E-02 |
| REACTOME_TGF_BETA_RECEPTOR_SIGNALING_ACTIVATES_SMADS | 2.07 | 1.41E-02 |
| PID_CASPASE_PATHWAY | 2.06 | 1.42E-02 |
| REACTOME_GAP_JUNCTION_TRAFFICKING_AND_REGULATION | 2.06 | 1.42E-02 |
| REACTOME_RHOJ_GTPASE_CYCLE | 2.06 | 1.43E-02 |
| SIG_PIP3_SIGNALING_IN_CARDIAC_MYOCTES | 2.06 | 1.43E-02 |
| KEGG_NEUROTROPHIN_SIGNALING_PATHWAY | 2.06 | 1.43E-02 |
| WP_PARKINUBIQUITIN_PROTEASOMAL_SYSTEM_PATHWAY | 2.06 | 1.48E-02 |
| WP_CORTICOTROPINRELEASING_HORMONE_SIGNALING_PATHWAY | 2.05 | 1.49E-02 |
| REACTOME_TLR3_MEDIATED_TICAM1_DEPENDENT_PROGRAMMED_CELL_DEATH | 2.05 | 1.50E-02 |
| SA_G1_AND_S_PHASES | 2.05 | 1.51E-02 |
| WP_IL3_SIGNALING_PATHWAY | 2.05 | 1.54E-02 |
| REACTOME_SELENOAMINO_ACID_METABOLISM | 2.05 | 1.54E-02 |
| WP_TLR4_SIGNALING_AND_TOLERANCE | 2.05 | 1.56E-02 |
| BIOCARTA_PKC_PATHWAY | 2.04 | 1.56E-02 |
| KEGG_PRIMARY_IMMUNODEFICIENCY | 2.04 | 1.57E-02 |
| REACTOME_NEGATIVE_REGULATION_OF_FLT3 | 2.04 | 1.58E-02 |
| WP_APOPTOSIS_MODULATION_AND_SIGNALING | 2.04 | 1.58E-02 |
| KEGG_VIRAL_MYOCARDITIS | 2.04 | 1.60E-02 |
| WP_TP53_NETWORK | 2.04 | 1.60E-02 |
| REACTOME_SIGNALING_BY_BRAF_AND_RAF1_FUSIONS | 2.04 | 1.62E-02 |
| REACTOME_INFECTION_WITH_MYCOBACTERIUM_TUBERCULOSIS | 2.04 | 1.63E-02 |
| REACTOME_REGULATION_OF_TLR_BY_ENDOGENOUS_LIGAND | 2.04 | 1.64E-02 |
| WP_CYTOKINES_AND_INFLAMMATORY_RESPONSE | 2.04 | 1.64E-02 |
| REACTOME_RETINOID_CYCLE_DISEASE_EVENTS | 2.03 | 1.64E-02 |
| REACTOME_FACTORS_INVOLVED_IN_MEGAKARYOCYTE_DEVELOPMENT_AND_PLATELET_PRODUCTION | 2.03 | 1.66E-02 |
| REACTOME_P75NTR_REGULATES_AXONOGENESIS | 2.03 | 1.67E-02 |
| REACTOME_TRANSCRIPTIONAL_REGULATION_BY_RUNX2 | 2.03 | 1.67E-02 |
| REACTOME_CYCLIN_A_B1_B2_ASSOCIATED_EVENTS_DURING_G2_M_TRANSITION | 2.03 | 1.67E-02 |
| PID_ECADHERIN_KERATINOCYTE_PATHWAY | 2.03 | 1.68E-02 |
| BIOCARTA_SRCRPTP_PATHWAY | 2.03 | 1.68E-02 |
| REACTOME_O_LINKED_GLYCOSYLATION_OF_MUCINS | 2.03 | 1.69E-02 |
| REACTOME_SEMA4D_IN_SEMAPHORIN_SIGNALING | 2.03 | 1.68E-02 |
| BIOCARTA_FAS_PATHWAY | 2.03 | 1.70E-02 |
| WP_NANOPARTICLE_TRIGGERED_REGULATED_NECROSIS | 2.02 | 1.72E-02 |
| WP_HEPATOCYTE_GROWTH_FACTOR_RECEPTOR_SIGNALING | 2.02 | 1.73E-02 |
| BIOCARTA_SODD_PATHWAY | 2.02 | 1.73E-02 |
| WP_EPITHELIAL_TO_MESENCHYMAL_TRANSITION_IN_COLORECTAL_CANCER | 2.02 | 1.73E-02 |
| BIOCARTA_FREE_PATHWAY | 2.02 | 1.74E-02 |
| REACTOME_MET_INTERACTS_WITH_TNS_PROTEINS | 2.02 | 1.76E-02 |
| PID_RHOA_PATHWAY | 2.02 | 1.79E-02 |
| WP_CYTOSOLIC_DNASENSING_PATHWAY | 2.01 | 1.80E-02 |
| KEGG_AXON_GUIDANCE | 2.01 | 1.80E-02 |
| WP_CHROMOSOMAL_AND_MICROSATELLITE_INSTABILITY_IN_COLORECTAL_CANCER | 2.01 | 1.81E-02 |
| REACTOME_NCAM_SIGNALING_FOR_NEURITE_OUT_GROWTH | 2.01 | 1.82E-02 |
| REACTOME_CONSTITUTIVE_SIGNALING_BY_ABERRANT_PI3K_IN_CANCER | 2.01 | 1.83E-02 |
| REACTOME_RHO_GTPASES_ACTIVATE_ROCKS | 2.01 | 1.84E-02 |
| REACTOME_HCMV_INFECTION | 2.01 | 1.84E-02 |
| REACTOME_ACTIVATION_OF_THE_PRE_REPLICATIVE_COMPLEX | 2.01 | 1.85E-02 |
| REACTOME_ROLE_OF_LAT2_NTAL_LAB_ON_CALCIUM_MOBILIZATION | 2.00 | 1.89E-02 |
| KEGG_TGF_BETA_SIGNALING_PATHWAY | 2.00 | 1.89E-02 |
| WP_HOSTPATHOGEN_INTERACTION_OF_HUMAN_CORONAVIRUSES_APOPTOSIS | 2.00 | 1.91E-02 |
| REACTOME_CLEC7A_INFLAMMASOME_PATHWAY | 2.00 | 1.93E-02 |
| REACTOME_CHONDROITIN_SULFATE_BIOSYNTHESIS | 2.00 | 1.93E-02 |
| REACTOME_PYROPTOSIS | 2.00 | 1.93E-02 |
| REACTOME_STABILIZATION_OF_P53 | 2.00 | 1.93E-02 |
| REACTOME_RHO_GTPASES_ACTIVATE_NADPH_OXIDASES | 2.00 | 1.93E-02 |
| WP_METABOLISM_OF_SPHINGOLIPIDS_IN_ER_AND_GOLGI_APPARATUS | 2.00 | 1.94E-02 |
| PID_RHOA_REG_PATHWAY | 2.00 | 1.94E-02 |
| REACTOME_GOLGI_ASSOCIATED_VESICLE_BIOGENESIS | 2.00 | 1.95E-02 |
| BIOCARTA_EIF2_PATHWAY | 2.00 | 1.96E-02 |
| WP_GASTRIC_CANCER_NETWORK_1 | 1.99 | 1.96E-02 |
| KEGG_NOD_LIKE_RECEPTOR_SIGNALING_PATHWAY | 1.99 | 2.03E-02 |
| KEGG_GLYCOSPHINGOLIPID_BIOSYNTHESIS_GLOBO_SERIES | 1.99 | 2.04E-02 |
| WP_HYPOTHESIZED_PATHWAYS_IN_PATHOGENESIS_OF_CARDIOVASCULAR_DISEASE | 1.99 | 2.04E-02 |
| BIOCARTA_NPC_PATHWAY | 1.99 | 2.04E-02 |
| WP_HYPERTROPHY_MODEL | 1.99 | 2.04E-02 |
| REACTOME_RUNX3_REGULATES_IMMUNE_RESPONSE_AND_CELL_MIGRATION | 1.99 | 2.04E-02 |
| REACTOME_CELL_EXTRACELLULAR_MATRIX_INTERACTIONS | 1.98 | 2.14E-02 |
| REACTOME_ACTIVATION_OF_THE_MRNA_UPON_BINDING_OF_THE_CAP_BINDING_COMPLEX_AND_EIFS_AND_SUBSEQUENT_BINDING_TO_43S | 1.98 | 2.16E-02 |
| REACTOME_SIGNALING_BY_MODERATE_KINASE_ACTIVITY_BRAF_MUTANTS | 1.98 | 2.16E-02 |
| WP_OSTEOCLAST_SIGNALING | 1.98 | 2.16E-02 |
| BIOCARTA_PAR1_PATHWAY | 1.97 | 2.19E-02 |
| REACTOME_DNA_REPLICATION_INITIATION | 1.97 | 2.22E-02 |
| BIOCARTA_P38MAPK_PATHWAY | 1.97 | 2.22E-02 |
| KEGG_OOCYTE_MEIOSIS | 1.97 | 2.23E-02 |
| WP_THYROID_STIMULATING_HORMONE_TSH_SIGNALING_PATHWAY | 1.97 | 2.22E-02 |
| REACTOME_AMINO_ACID_TRANSPORT_ACROSS_THE_PLASMA_MEMBRANE | 1.97 | 2.23E-02 |
| REACTOME_ANTIGEN_ACTIVATES_B_CELL_RECEPTOR_BCR_LEADING_TO_GENERATION_OF_SECOND_MESSENGERS | 1.97 | 2.23E-02 |
| WP_PATHOGENESIS_OF_SARSCOV2_MEDIATED_BY_NSP9NSP10_COMPLEX | 1.97 | 2.23E-02 |
| REACTOME_TP53_REGULATES_TRANSCRIPTION_OF_GENES_INVOLVED_IN_G2_CELL_CYCLE_ARREST | 1.97 | 2.25E-02 |
| PID_LYSOPHOSPHOLIPID_PATHWAY | 1.97 | 2.25E-02 |
| REACTOME_COPI_MEDIATED_ANTEROGRADE_TRANSPORT | 1.97 | 2.26E-02 |
| WP_CANONICAL_AND_NONCANONICAL_TGFB_SIGNALING | 1.96 | 2.27E-02 |
| REACTOME_HEDGEHOG_OFF_STATE | 1.96 | 2.30E-02 |
| KEGG_SYSTEMIC_LUPUS_ERYTHEMATOSUS | 1.96 | 2.31E-02 |
| REACTOME_REGULATION_OF_PTEN_STABILITY_AND_ACTIVITY | 1.96 | 2.30E-02 |
| KEGG_TIGHT_JUNCTION | 1.96 | 2.31E-02 |
| WP_MAMMARY_GLAND_DEVELOPMENT_PATHWAY_EMBRYONIC_DEVELOPMENT_STAGE_1_OF_4 | 1.96 | 2.30E-02 |
| PID_TCR_JNK_PATHWAY | 1.96 | 2.31E-02 |
| REACTOME_UNFOLDED_PROTEIN_RESPONSE_UPR | 1.96 | 2.33E-02 |
| BIOCARTA_CDMAC_PATHWAY | 1.96 | 2.34E-02 |
| KEGG_APOPTOSIS | 1.96 | 2.34E-02 |
| HALLMARK_PI3K_AKT_MTOR_SIGNALING | 1.95 | 2.36E-02 |
| WP_HAIR_FOLLICLE_DEVELOPMENT_ORGANOGENESIS_PART_2_OF_3 | 1.95 | 2.42E-02 |
| PID_TGFBR_PATHWAY | 1.95 | 2.42E-02 |
| REACTOME_INTRACELLULAR_SIGNALING_BY_SECOND_MESSENGERS | 1.95 | 2.42E-02 |
| WP_GASTRIN_SIGNALING_PATHWAY | 1.95 | 2.43E-02 |
| REACTOME_REGULATION_OF_RUNX2_EXPRESSION_AND_ACTIVITY | 1.95 | 2.47E-02 |
| PID_MYC_ACTIV_PATHWAY | 1.95 | 2.47E-02 |
| REACTOME_O_LINKED_GLYCOSYLATION | 1.95 | 2.47E-02 |
| REACTOME_L1CAM_INTERACTIONS | 1.94 | 2.48E-02 |
| BIOCARTA_CARDIACEGF_PATHWAY | 1.94 | 2.55E-02 |
| WP_FOXP3_IN_COVID19 | 1.94 | 2.56E-02 |
| WP_ACTIVATION_OF_NLRP3_INFLAMMASOME_BY_SARSCOV2 | 1.94 | 2.56E-02 |
| SA_PROGRAMMED_CELL_DEATH | 1.94 | 2.56E-02 |
| REACTOME_RUNX1_REGULATES_TRANSCRIPTION_OF_GENES_INVOLVED_IN_DIFFERENTIATION_OF_MYELOID_CELLS | 1.94 | 2.56E-02 |
| KEGG_SPLICEOSOME | 1.93 | 2.59E-02 |
| WP_METASTATIC_BRAIN_TUMOR | 1.93 | 2.60E-02 |
| REACTOME_DEGRADATION_OF_DVL | 1.93 | 2.61E-02 |
| PID_BCR_5PATHWAY | 1.93 | 2.61E-02 |
| BIOCARTA_BAD_PATHWAY | 1.93 | 2.62E-02 |
| WP_PURINERGIC_SIGNALING | 1.93 | 2.63E-02 |
| BIOCARTA_TNFR1_PATHWAY | 1.93 | 2.65E-02 |
| REACTOME_NUCLEOTIDE_BINDING_DOMAIN_LEUCINE_RICH_REPEAT_CONTAINING_RECEPTOR_NLR_SIGNALING_PATHWAYS | 1.93 | 2.66E-02 |
| WP_MELANOMA | 1.93 | 2.66E-02 |
| REACTOME_GRB2_SOS_PROVIDES_LINKAGE_TO_MAPK_SIGNALING_FOR_INTEGRINS | 1.93 | 2.66E-02 |
| BIOCARTA_TCR_PATHWAY | 1.93 | 2.67E-02 |
| PID_CXCR3_PATHWAY | 1.92 | 2.70E-02 |
| KEGG_INTESTINAL_IMMUNE_NETWORK_FOR_IGA_PRODUCTION | 1.92 | 2.71E-02 |
| REACTOME_FOLDING_OF_ACTIN_BY_CCT_TRIC | 1.92 | 2.73E-02 |
| REACTOME_PTK6_PROMOTES_HIF1A_STABILIZATION | 1.92 | 2.75E-02 |
| REACTOME_ACTIVATION_OF_NMDA_RECEPTORS_AND_POSTSYNAPTIC_EVENTS | 1.92 | 2.77E-02 |
| REACTOME_IRON_UPTAKE_AND_TRANSPORT | 1.92 | 2.78E-02 |
| REACTOME_GPCR_LIGAND_BINDING | 1.92 | 2.78E-02 |
| REACTOME_NEUROTRANSMITTER_RECEPTORS_AND_POSTSYNAPTIC_SIGNAL_TRANSMISSION | 1.92 | 2.79E-02 |
| REACTOME_STRIATED_MUSCLE_CONTRACTION | 1.92 | 2.80E-02 |
| WP_APOE_AND_MIR146_IN_INFLAMMATION_AND_ATHEROSCLEROSIS | 1.92 | 2.81E-02 |
| WP_NRP1TRIGGERED_SIGNALING_PATHWAYS_IN_PANCREATIC_CANCER | 1.91 | 2.84E-02 |
| REACTOME_SENESCENCE_ASSOCIATED_SECRETORY_PHENOTYPE_SASP | 1.91 | 2.86E-02 |
| REACTOME_RAF_INDEPENDENT_MAPK1_3_ACTIVATION | 1.91 | 2.86E-02 |
| REACTOME_SIGNAL_TRANSDUCTION_BY_L1 | 1.91 | 2.86E-02 |
| BIOCARTA_IL1R_PATHWAY | 1.91 | 2.86E-02 |
| BIOCARTA_AMI_PATHWAY | 1.91 | 2.86E-02 |
| KEGG_COLORECTAL_CANCER | 1.91 | 2.91E-02 |
| WP_GASTRIC_CANCER_NETWORK_2 | 1.90 | 2.94E-02 |
| REACTOME_INTEGRIN_SIGNALING | 1.90 | 2.94E-02 |
| BIOCARTA_PRION_PATHWAY | 1.90 | 2.94E-02 |
| REACTOME_REGULATION_OF_IFNA_IFNB_SIGNALING | 1.90 | 2.97E-02 |
| WP_NITRIC_OXIDE_METABOLISM_IN_CYSTIC_FIBROSIS | 1.90 | 3.01E-02 |
| REACTOME_NEF_MEDIATES_DOWN_MODULATION_OF_CELL_SURFACE_RECEPTORS_BY_RECRUITING_THEM_TO_CLATHRIN_ADAPTERS | 1.90 | 3.03E-02 |
| REACTOME_INTERACTIONS_OF_REV_WITH_HOST_CELLULAR_PROTEINS | 1.90 | 3.05E-02 |
| PID_ARF6_DOWNSTREAM_PATHWAY | 1.89 | 3.07E-02 |
| WP_EXTRACELLULAR_VESICLES_IN_THE_CROSSTALK_OF_CARDIAC_CELLS | 1.89 | 3.07E-02 |
| BIOCARTA_S1P_PATHWAY | 1.89 | 3.07E-02 |
| REACTOME_PTK6_REGULATES_RHO_GTPASES_RAS_GTPASE_AND_MAP_KINASES | 1.89 | 3.09E-02 |
| REACTOME_CD28_DEPENDENT_VAV1_PATHWAY | 1.89 | 3.12E-02 |
| REACTOME_ER_TO_GOLGI_ANTEROGRADE_TRANSPORT | 1.89 | 3.12E-02 |
| REACTOME_SEMA3A_PAK_DEPENDENT_AXON_REPULSION | 1.89 | 3.14E-02 |
| BIOCARTA_RNA_PATHWAY | 1.89 | 3.15E-02 |
| REACTOME_DEPOLYMERISATION_OF_THE_NUCLEAR_LAMINA | 1.89 | 3.15E-02 |
| PID_ILK_PATHWAY | 1.89 | 3.15E-02 |
| REACTOME_HEPARAN_SULFATE_HEPARIN_HS_GAG_METABOLISM | 1.89 | 3.16E-02 |
| REACTOME_SIGNALING_BY_ALK_IN_CANCER | 1.89 | 3.17E-02 |
| REACTOME_SIGNAL_REGULATORY_PROTEIN_FAMILY_INTERACTIONS | 1.88 | 3.22E-02 |
| REACTOME_REGULATED_NECROSIS | 1.88 | 3.23E-02 |
| BIOCARTA_MCM_PATHWAY | 1.88 | 3.24E-02 |
| BIOCARTA_GSK3_PATHWAY | 1.88 | 3.28E-02 |
| REACTOME_NEGATIVE_REGULATION_OF_THE_PI3K_AKT_NETWORK | 1.88 | 3.28E-02 |
| BIOCARTA_PROTEASOME_PATHWAY | 1.88 | 3.28E-02 |
| REACTOME_TP53_REGULATES_TRANSCRIPTION_OF_GENES_INVOLVED_IN_G1_CELL_CYCLE_ARREST | 1.88 | 3.29E-02 |
| REACTOME_CD163_MEDIATING_AN_ANTI_INFLAMMATORY_RESPONSE | 1.88 | 3.29E-02 |
| WP_THYMIC_STROMAL_LYMPHOPOIETIN_TSLP_SIGNALING_PATHWAY | 1.88 | 3.28E-02 |
| WP_DNA_REPLICATION | 1.88 | 3.29E-02 |
| REACTOME_CONDENSATION_OF_PROMETAPHASE_CHROMOSOMES | 1.88 | 3.29E-02 |
| REACTOME_TRANSCRIPTIONAL_REGULATION_BY_THE_AP_2_TFAP2_FAMILY_OF_TRANSCRIPTION_FACTORS | 1.88 | 3.32E-02 |
| REACTOME_AGGREPHAGY | 1.87 | 3.35E-02 |
| REACTOME_G1_S_SPECIFIC_TRANSCRIPTION | 1.87 | 3.35E-02 |
| REACTOME_DERMATAN_SULFATE_BIOSYNTHESIS | 1.87 | 3.38E-02 |
| BIOCARTA_NFKB_PATHWAY | 1.87 | 3.41E-02 |
| BIOCARTA_EFP_PATHWAY | 1.87 | 3.42E-02 |
| REACTOME_PHOSPHORYLATION_OF_EMI1 | 1.87 | 3.45E-02 |
| BIOCARTA_TCYTOTOXIC_PATHWAY | 1.87 | 3.45E-02 |
| WP_COMPLEMENT_AND_COAGULATION_CASCADES | 1.86 | 3.51E-02 |
| REACTOME_REGULATION_OF_KIT_SIGNALING | 1.86 | 3.51E-02 |
| PID_ATR_PATHWAY | 1.86 | 3.61E-02 |
| REACTOME_ANCHORING_FIBRIL_FORMATION | 1.86 | 3.63E-02 |
| KEGG_DILATED_CARDIOMYOPATHY | 1.86 | 3.65E-02 |
| REACTOME_RHOB_GTPASE_CYCLE | 1.86 | 3.65E-02 |
| PID_CD8_TCR_DOWNSTREAM_PATHWAY | 1.85 | 3.67E-02 |
| REACTOME_SYNAPTIC_ADHESION_LIKE_MOLECULES | 1.85 | 3.67E-02 |
| REACTOME_CHK1_CHK2_CDS1_MEDIATED_INACTIVATION_OF_CYCLIN_B_CDK1_COMPLEX | 1.85 | 3.69E-02 |
| REACTOME_FORMATION_OF_TUBULIN_FOLDING_INTERMEDIATES_BY_CCT_TRIC | 1.85 | 3.70E-02 |
| WP_INTERACTIONS_OF_NATURAL_KILLER_CELLS_IN_PANCREATIC_CANCER | 1.85 | 3.74E-02 |
| REACTOME_INSULIN_RECEPTOR_RECYCLING | 1.85 | 3.73E-02 |
| REACTOME_NUCLEAR_IMPORT_OF_REV_PROTEIN | 1.85 | 3.74E-02 |
| REACTOME_CELL_JUNCTION_ORGANIZATION | 1.85 | 3.78E-02 |
| BIOCARTA_ACH_PATHWAY | 1.85 | 3.77E-02 |
| REACTOME_E2F_MEDIATED_REGULATION_OF_DNA_REPLICATION | 1.85 | 3.79E-02 |
| BIOCARTA_INFLAM_PATHWAY | 1.85 | 3.79E-02 |
| REACTOME_BIOSYNTHESIS_OF_THE_N_GLYCAN_PRECURSOR_DOLICHOL_LIPID_LINKED_OLIGOSACCHARIDE_LLO_AND_TRANSFER_TO_A_NASCENT_PROTEIN | 1.84 | 3.82E-02 |
| WP_SEROTONIN_AND_ANXIETYRELATED_EVENTS | 1.84 | 3.82E-02 |
| REACTOME_SIGNALING_BY_NOTCH | 1.84 | 3.83E-02 |
| BIOCARTA_THELPER_PATHWAY | 1.84 | 3.85E-02 |
| REACTOME_INHIBITION_OF_REPLICATION_INITIATION_OF_DAMAGED_DNA_BY_RB1_E2F1 | 1.84 | 3.85E-02 |
| REACTOME_GAB1_SIGNALOSOME | 1.84 | 3.85E-02 |
| WP_PRIMARY_FOCAL_SEGMENTAL_GLOMERULOSCLEROSIS_FSGS | 1.84 | 3.85E-02 |
| REACTOME_FCERI_MEDIATED_CA_2_MOBILIZATION | 1.84 | 3.87E-02 |
| REACTOME_SIGNALING_BY_SCF_KIT | 1.84 | 3.92E-02 |
| REACTOME_KERATAN_SULFATE_BIOSYNTHESIS | 1.84 | 3.96E-02 |
| REACTOME_NS1_MEDIATED_EFFECTS_ON_HOST_PATHWAYS | 1.83 | 4.01E-02 |
| REACTOME_RHOH_GTPASE_CYCLE | 1.83 | 4.07E-02 |
| BIOCARTA_GCR_PATHWAY | 1.83 | 4.07E-02 |
| REACTOME_TRAFFICKING_OF_AMPA_RECEPTORS | 1.83 | 4.11E-02 |
| PID_IL4_2PATHWAY | 1.83 | 4.14E-02 |
| REACTOME_HCMV_LATE_EVENTS | 1.82 | 4.18E-02 |
| REACTOME_NEGATIVE_REGULATION_OF_MET_ACTIVITY | 1.82 | 4.17E-02 |
| WP_MICRORNAS_IN_CARDIOMYOCYTE_HYPERTROPHY | 1.82 | 4.17E-02 |
| PID_IL8_CXCR1_PATHWAY | 1.82 | 4.21E-02 |
| BIOCARTA_NOS1_PATHWAY | 1.82 | 4.22E-02 |
| BIOCARTA_CHEMICAL_PATHWAY | 1.82 | 4.24E-02 |
| BIOCARTA_RANMS_PATHWAY | 1.82 | 4.25E-02 |
| WP_OSTEOPONTIN_SIGNALING | 1.82 | 4.25E-02 |
| REACTOME_NEF_MEDIATED_CD4_DOWN_REGULATION | 1.82 | 4.28E-02 |
| REACTOME_KERATAN_SULFATE_DEGRADATION | 1.82 | 4.28E-02 |
| REACTOME_TRANSMISSION_ACROSS_CHEMICAL_SYNAPSES | 1.82 | 4.28E-02 |
| PID_ALPHA_SYNUCLEIN_PATHWAY | 1.82 | 4.31E-02 |
| REACTOME_WNT5A_DEPENDENT_INTERNALIZATION_OF_FZD4 | 1.81 | 4.36E-02 |
| REACTOME_DISEASES_ASSOCIATED_WITH_N_GLYCOSYLATION_OF_PROTEINS | 1.81 | 4.36E-02 |
| REACTOME_RESPONSE_OF_MTB_TO_PHAGOCYTOSIS | 1.81 | 4.40E-02 |
| BIOCARTA_RANKL_PATHWAY | 1.81 | 4.40E-02 |
| REACTOME_DEFECTIVE_B4GALT7_CAUSES_EDS_PROGEROID_TYPE | 1.81 | 4.40E-02 |
| WP_OMEGA9_FATTY_ACID_SYNTHESIS | 1.81 | 4.41E-02 |
| REACTOME_SMAD2_SMAD3_SMAD4_HETEROTRIMER_REGULATES_TRANSCRIPTION | 1.81 | 4.41E-02 |
| PID_GMCSF_PATHWAY | 1.81 | 4.40E-02 |
| SA_B_CELL_RECEPTOR_COMPLEXES | 1.81 | 4.40E-02 |
| SIG_PIP3_SIGNALING_IN_B_LYMPHOCYTES | 1.81 | 4.40E-02 |
| WP_MRNA_VACCINE_ACTIVATION_OF_DENDRITIC_CELL_AND_INDUCTION_OF_IFN1 | 1.81 | 4.40E-02 |
| REACTOME_G_ALPHA_12_13_SIGNALLING_EVENTS | 1.81 | 4.40E-02 |
| WP_HEAD_AND_NECK_SQUAMOUS_CELL_CARCINOMA | 1.81 | 4.41E-02 |
| WP_MECHANOREGULATION_AND_PATHOLOGY_OF_YAPTAZ_VIA_HIPPO_AND_NONHIPPO_MECHANISMS | 1.81 | 4.40E-02 |
| BIOCARTA_SPPA_PATHWAY | 1.81 | 4.41E-02 |
| REACTOME_NUCLEAR_PORE_COMPLEX_NPC_DISASSEMBLY | 1.81 | 4.42E-02 |
| KEGG_BLADDER_CANCER | 1.81 | 4.43E-02 |
| REACTOME_INTERLEUKIN_6_FAMILY_SIGNALING | 1.81 | 4.45E-02 |
| BIOCARTA_TEL_PATHWAY | 1.81 | 4.46E-02 |
| REACTOME_ACTIVATION_OF_MATRIX_METALLOPROTEINASES | 1.80 | 4.47E-02 |
| WP_WNT_SIGNALING | 1.80 | 4.56E-02 |
| REACTOME_PLATELET_AGGREGATION_PLUG_FORMATION | 1.80 | 4.59E-02 |
| BIOCARTA_TID_PATHWAY | 1.80 | 4.62E-02 |
| WP_OLIGODENDROCYTE_SPECIFICATION_AND_DIFFERENTIATION_LEADING_TO_MYELIN_COMPONENTS_FOR_CNS | 1.80 | 4.62E-02 |
| HALLMARK_REACTIVE_OXYGEN_SPECIES_PATHWAY | 1.80 | 4.61E-02 |
| PID_S1P_S1P2_PATHWAY | 1.80 | 4.64E-02 |
| WP_SIGNAL_TRANSDUCTION_THROUGH_IL1R | 1.80 | 4.65E-02 |
| REACTOME_TRANSLATION_OF_SARS_COV_2_STRUCTURAL_PROTEINS | 1.80 | 4.64E-02 |
| KEGG_PROGESTERONE_MEDIATED_OOCYTE_MATURATION | 1.80 | 4.65E-02 |
| REACTOME_PTEN_REGULATION | 1.79 | 4.67E-02 |
| REACTOME_RHO_GTPASES_ACTIVATE_CIT | 1.79 | 4.69E-02 |
| REACTOME_INTERLEUKIN_37_SIGNALING | 1.79 | 4.73E-02 |
| KEGG_WNT_SIGNALING_PATHWAY | 1.79 | 4.78E-02 |
| WP_KIT_RECEPTOR_SIGNALING_PATHWAY | 1.79 | 4.79E-02 |
| REACTOME_SPHINGOLIPID_DE_NOVO_BIOSYNTHESIS | 1.79 | 4.81E-02 |
| REACTOME_SUPPRESSION_OF_PHAGOSOMAL_MATURATION | 1.79 | 4.81E-02 |
| REACTOME_LDL_CLEARANCE | 1.79 | 4.83E-02 |
| REACTOME_RHOD_GTPASE_CYCLE | 1.79 | 4.86E-02 |
| REACTOME_TRANSPORT_OF_CONNEXONS_TO_THE_PLASMA_MEMBRANE | 1.78 | 4.88E-02 |
| REACTOME_RUNX3_REGULATES_P14_ARF | 1.78 | 4.92E-02 |
| WP_DNA_IRDOUBLE_STRAND_BREAKS_AND_CELLULAR_RESPONSE_VIA_ATM | 1.78 | 4.91E-02 |
| WP_RESOLVIN_E1_AND_RESOLVIN_D1_SIGNALING_PATHWAYS_PROMOTING_INFLAMMATION_RESOLUTION | 1.78 | 4.91E-02 |
| PID_MYC_REPRESS_PATHWAY | 1.78 | 4.94E-02 |
| REACTOME_MITOTIC_PROPHASE | 1.78 | 4.97E-02 |
| HALLMARK_OXIDATIVE_PHOSPHORYLATION | -9.42 | 0.00E+00 |
| REACTOME_THE_CITRIC_ACID_TCA_CYCLE_AND_RESPIRATORY_ELECTRON_TRANSPORT | -9.08 | 0.00E+00 |
| REACTOME_RESPIRATORY_ELECTRON_TRANSPORT | -7.89 | 0.00E+00 |
| REACTOME_RESPIRATORY_ELECTRON_TRANSPORT_ATP_SYNTHESIS_BY_CHEMIOSMOTIC_COUPLING_AND_HEAT_PRODUCTION_BY_UNCOUPLING_PROTEINS | -7.79 | 0.00E+00 |
| WP_ELECTRON_TRANSPORT_CHAIN_OXPHOS_SYSTEM_IN_MITOCHONDRIA | -7.38 | 0.00E+00 |
| REACTOME_MITOCHONDRIAL_TRANSLATION | -6.75 | 0.00E+00 |
| REACTOME_COMPLEX_I_BIOGENESIS | -6.57 | 0.00E+00 |
| KEGG_OXIDATIVE_PHOSPHORYLATION | -6.55 | 0.00E+00 |
| KEGG_PARKINSONS_DISEASE | -6.53 | 0.00E+00 |
| WP_MITOCHONDRIAL_COMPLEX_I_ASSEMBLY_MODEL_OXPHOS_SYSTEM | -6.28 | 0.00E+00 |
| WP_OXIDATIVE_PHOSPHORYLATION | -5.74 | 0.00E+00 |
| WP_NONALCOHOLIC_FATTY_LIVER_DISEASE | -5.44 | 0.00E+00 |
| HALLMARK_ADIPOGENESIS | -5.41 | 0.00E+00 |
| REACTOME_PROTEIN_LOCALIZATION | -5.25 | 0.00E+00 |
| KEGG_ALZHEIMERS_DISEASE | -5.23 | 0.00E+00 |
| REACTOME_MITOCHONDRIAL_PROTEIN_IMPORT | -5.21 | 0.00E+00 |
| HALLMARK_FATTY_ACID_METABOLISM | -5.16 | 0.00E+00 |
| REACTOME_PYRUVATE_METABOLISM_AND_CITRIC_ACID_TCA_CYCLE | -5.07 | 0.00E+00 |
| REACTOME_MITOCHONDRIAL_FATTY_ACID_BETA_OXIDATION | -4.95 | 0.00E+00 |
| KEGG_HUNTINGTONS_DISEASE | -4.93 | 0.00E+00 |
| REACTOME_CITRIC_ACID_CYCLE_TCA_CYCLE | -4.72 | 0.00E+00 |
| REACTOME_FATTY_ACID_METABOLISM | -4.68 | 0.00E+00 |
| KEGG_VALINE_LEUCINE_AND_ISOLEUCINE_DEGRADATION | -4.65 | 0.00E+00 |
| WP_LEUCINE_ISOLEUCINE_AND_VALINE_METABOLISM | -4.35 | 0.00E+00 |
| KEGG_CARDIAC_MUSCLE_CONTRACTION | -4.31 | 0.00E+00 |
| WP_TCA_CYCLE_AKA_KREBS_OR_CITRIC_ACID_CYCLE | -4.28 | 0.00E+00 |
| WP_FATTY_ACID_BETAOXIDATION | -4.06 | 0.00E+00 |
| REACTOME_BRANCHED_CHAIN_AMINO_ACID_CATABOLISM | -4.00 | 0.00E+00 |
| KEGG_CITRATE_CYCLE_TCA_CYCLE | -3.98 | 0.00E+00 |
| KEGG_PROPANOATE_METABOLISM | -3.70 | 0.00E+00 |
| WP_MITOCHONDRIAL_LONG_CHAIN_FATTY_ACID_BETAOXIDATION | -3.65 | 0.00E+00 |
| WP_MITOCHONDRIAL_COMPLEX_IV_ASSEMBLY | -3.63 | 0.00E+00 |
| REACTOME_GLYOXYLATE_METABOLISM_AND_GLYCINE_DEGRADATION | -3.62 | 0.00E+00 |
| KEGG_FATTY_ACID_METABOLISM | -3.55 | 0.00E+00 |
| REACTOME_MITOCHONDRIAL_BIOGENESIS | -3.48 | 0.00E+00 |
| REACTOME_PYRUVATE_METABOLISM | -3.31 | 0.00E+00 |
| KEGG_PEROXISOME | -3.30 | 0.00E+00 |
| REACTOME_REGULATION_OF_PYRUVATE_DEHYDROGENASE_PDH_COMPLEX | -3.17 | 0.00E+00 |
| REACTOME_PEROXISOMAL_PROTEIN_IMPORT | -3.17 | 0.00E+00 |
| WP_AMINO_ACID_METABOLISM | -3.16 | 0.00E+00 |
| WP_MITOCHONDRIAL_COMPLEX_III_ASSEMBLY | -3.12 | 0.00E+00 |
| REACTOME_MITOCHONDRIAL_FATTY_ACID_BETA_OXIDATION_OF_SATURATED_FATTY_ACIDS | -3.10 | 0.00E+00 |
| KEGG_PYRUVATE_METABOLISM | -3.04 | 0.00E+00 |
| BIOCARTA_KREB_PATHWAY | -3.01 | 0.00E+00 |
| REACTOME_CYTOPROTECTION_BY_HMOX1 | -2.99 | 1.14E-05 |
| REACTOME_UBIQUINOL_BIOSYNTHESIS | -2.91 | 4.57E-05 |
| REACTOME_TRANSCRIPTIONAL_ACTIVATION_OF_MITOCHONDRIAL_BIOGENESIS | -2.89 | 1.00E-04 |
| REACTOME_MITOCHONDRIAL_FATTY_ACID_BETA_OXIDATION_OF_UNSATURATED_FATTY_ACIDS | -2.87 | 1.20E-04 |
| REACTOME_SULFUR_AMINO_ACID_METABOLISM | -2.85 | 1.39E-04 |
| REACTOME_DEFECTS_IN_VITAMIN_AND_COFACTOR_METABOLISM | -2.84 | 1.36E-04 |
| REACTOME_SIGNALING_BY_RETINOIC_ACID | -2.81 | 1.44E-04 |
| KEGG_BUTANOATE_METABOLISM | -2.81 | 1.41E-04 |
| WP_CEREBRAL_ORGANIC_ACIDURIAS_INCLUDING_DISEASES | -2.79 | 1.59E-04 |
| HALLMARK_BILE_ACID_METABOLISM | -2.79 | 1.56E-04 |
| REACTOME_TRANSLATION | -2.78 | 1.81E-04 |
| REACTOME_DEFECTS_IN_COBALAMIN_B12_METABOLISM | -2.72 | 2.73E-04 |
| REACTOME_BETA_OXIDATION_OF_OCTANOYL_COA_TO_HEXANOYL_COA | -2.72 | 2.68E-04 |
| REACTOME_PINK1_PRKN_MEDIATED_MITOPHAGY | -2.72 | 2.81E-04 |
| REACTOME_MITOCHONDRIAL_TRNA_AMINOACYLATION | -2.70 | 3.49E-04 |
| BIOCARTA_ETC_PATHWAY | -2.70 | 3.43E-04 |
| REACTOME_DEGRADATION_OF_CYSTEINE_AND_HOMOCYSTEINE | -2.68 | 3.73E-04 |
| REACTOME_TP53_REGULATES_METABOLIC_GENES | -2.67 | 3.84E-04 |
| REACTOME_RRNA_MODIFICATION_IN_THE_MITOCHONDRION | -2.67 | 3.86E-04 |
| REACTOME_BETA_OXIDATION_OF_LAUROYL_COA_TO_DECANOYL_COA_COA | -2.66 | 4.29E-04 |
| REACTOME_MITOCHONDRIAL_IRON_SULFUR_CLUSTER_BIOGENESIS | -2.64 | 5.29E-04 |
| REACTOME_BETA_OXIDATION_OF_HEXANOYL_COA_TO_BUTANOYL_COA | -2.62 | 6.01E-04 |
| KEGG_BETA_ALANINE_METABOLISM | -2.61 | 5.93E-04 |
| KEGG_TRYPTOPHAN_METABOLISM | -2.61 | 6.38E-04 |
| WP_RIBOFLAVIN_AND_COQ_DISORDERS | -2.60 | 6.52E-04 |
| REACTOME_PEROXISOMAL_LIPID_METABOLISM | -2.59 | 7.40E-04 |
| REACTOME_CRISTAE_FORMATION | -2.57 | 9.01E-04 |
| WP_MITOCHONDRIAL_COMPLEX_II_ASSEMBLY | -2.54 | 1.16E-03 |
| REACTOME_CARNITINE_METABOLISM | -2.52 | 1.35E-03 |
| KEGG_DRUG_METABOLISM_CYTOCHROME_P450 | -2.49 | 1.51E-03 |
| REACTOME_BETA_OXIDATION_OF_DECANOYL_COA_TO_OCTANOYL_COA_COA | -2.49 | 1.58E-03 |
| BIOCARTA_PPARG_PATHWAY | -2.49 | 1.58E-03 |
| WP_VITAMIN_B12_DISORDERS | -2.49 | 1.56E-03 |
| KEGG_PORPHYRIN_AND_CHLOROPHYLL_METABOLISM | -2.48 | 1.62E-03 |
| REACTOME_MITOCHONDRIAL_CALCIUM_ION_TRANSPORT | -2.47 | 1.84E-03 |
| KEGG_AMINOACYL_TRNA_BIOSYNTHESIS | -2.46 | 1.82E-03 |
| WP_KREBS_CYCLE_DISORDERS | -2.46 | 1.82E-03 |
| WP_MITOCHONDRIAL_BETAOXIDATION | -2.45 | 1.99E-03 |
| WP_TCA_CYCLE_AND_DEFICIENCY_OF_PYRUVATE_DEHYDROGENASE_COMPLEX_PDHC | -2.44 | 2.16E-03 |
| KEGG_GLYCOLYSIS_GLUCONEOGENESIS | -2.39 | 3.22E-03 |
| KEGG_METABOLISM_OF_XENOBIOTICS_BY_CYTOCHROME_P450 | -2.38 | 3.37E-03 |
| WP_PPAR_SIGNALING_PATHWAY | -2.37 | 3.60E-03 |
| REACTOME_PROPIONYL_COA_CATABOLISM | -2.36 | 3.85E-03 |
| KEGG_PPAR_SIGNALING_PATHWAY | -2.34 | 4.47E-03 |
| KEGG_CYSTEINE_AND_METHIONINE_METABOLISM | -2.34 | 4.47E-03 |
| REACTOME_PROCESSING_OF_CAPPED_INTRON_CONTAINING_PRE_MRNA | -2.34 | 4.44E-03 |
| REACTOME_SYNTHESIS_OF_PA | -2.33 | 4.70E-03 |
| WP_FATTY_ACID_BIOSYNTHESIS | -2.30 | 5.66E-03 |
| REACTOME_RRNA_PROCESSING_IN_THE_MITOCHONDRION | -2.30 | 5.62E-03 |
| KEGG_GLYCEROLIPID_METABOLISM | -2.28 | 6.58E-03 |
| WP_TRYPTOPHAN_METABOLISM | -2.27 | 6.80E-03 |
| REACTOME_COBALAMIN_CBL_METABOLISM | -2.27 | 6.83E-03 |
| REACTOME_COBALAMIN_CBL_VITAMIN_B12_TRANSPORT_AND_METABOLISM | -2.26 | 6.95E-03 |
| REACTOME_TRNA_AMINOACYLATION | -2.21 | 1.04E-02 |
| REACTOME_FOXO_MEDIATED_TRANSCRIPTION_OF_OXIDATIVE_STRESS_METABOLIC_AND_NEURONAL_GENES | -2.20 | 1.05E-02 |
| REACTOME_HATS_ACETYLATE_HISTONES | -2.20 | 1.08E-02 |
| REACTOME_MITOPHAGY | -2.18 | 1.18E-02 |
| HALLMARK_PEROXISOME | -2.18 | 1.22E-02 |
| REACTOME_CARDIAC_CONDUCTION | -2.18 | 1.23E-02 |
| REACTOME_REGULATION_OF_LIPID_METABOLISM_BY_PPARALPHA | -2.17 | 1.26E-02 |
| REACTOME_BIOLOGICAL_OXIDATIONS | -2.17 | 1.25E-02 |
| WP_FAMILIAL_PARTIAL_LIPODYSTROPHY | -2.17 | 1.27E-02 |
| REACTOME_METABOLISM_OF_COFACTORS | -2.17 | 1.28E-02 |
| REACTOME_PHENYLALANINE_AND_TYROSINE_METABOLISM | -2.16 | 1.30E-02 |
| WP_ARSENIC_METABOLISM_AND_REACTIVE_OXYGEN_SPECIES_GENERATION | -2.15 | 1.40E-02 |
| REACTOME_ACYL_CHAIN_REMODELING_OF_DAG_AND_TAG | -2.14 | 1.43E-02 |
| REACTOME_MIRO_GTPASE_CYCLE | -2.14 | 1.42E-02 |
| WP_HEME_BIOSYNTHESIS | -2.14 | 1.46E-02 |
| WP_NUCLEAR_RECEPTORS | -2.13 | 1.50E-02 |
| WP_GLYCOLYSIS_AND_GLUCONEOGENESIS | -2.13 | 1.50E-02 |
| WP_HEMESYNTHESIS_DEFECTS_AND_PORPHYRIAS | -2.12 | 1.55E-02 |
| WP_METHIONINE_METABOLISM_LEADING_TO_SULFUR_AMINO_ACIDS_AND_RELATED_DISORDERS | -2.12 | 1.62E-02 |
| WP_UREA_CYCLE_AND_ASSOCIATED_PATHWAYS | -2.12 | 1.61E-02 |
| WP_CYSTEINE_AND_METHIONINE_CATABOLISM | -2.11 | 1.65E-02 |
| REACTOME_GLUCONEOGENESIS | -2.11 | 1.68E-02 |
| REACTOME_GLYCEROPHOSPHOLIPID_BIOSYNTHESIS | -2.09 | 1.84E-02 |
| BIOCARTA_AHSP_PATHWAY | -2.09 | 1.84E-02 |
| NEGATIVE FERROPTOSIS REGULATORS | -2.09 | 1.84E-02 |
| ALL FERROPTOSIS | -2.07 | 2.00E-02 |
| REACTOME_GLYCOGEN_BREAKDOWN_GLYCOGENOLYSIS | -2.07 | 2.10E-02 |
| REACTOME_METABOLISM_OF_AMINO_ACIDS_AND_DERIVATIVES | -2.06 | 2.11E-02 |
| REACTOME_FOXO_MEDIATED_TRANSCRIPTION | -2.06 | 2.14E-02 |
| WP_FLUOROACETIC_ACID_TOXICITY | -2.06 | 2.18E-02 |
| REACTOME_METABOLISM_OF_PORPHYRINS | -2.05 | 2.25E-02 |
| HALLMARK_HEME_METABOLISM | -2.02 | 2.65E-02 |
| KEGG_TYROSINE_METABOLISM | -2.02 | 2.73E-02 |
| REACTOME_SULFIDE_OXIDATION_TO_SULFATE | -1.99 | 3.18E-02 |
| REACTOME_MRNA_SPLICING | -1.98 | 3.47E-02 |
| WP_7OXOC_AND_7BETAHC_PATHWAYS | -1.98 | 3.47E-02 |
| KEGG_HISTIDINE_METABOLISM | -1.97 | 3.65E-02 |
| WP_ENERGY_METABOLISM | -1.97 | 3.62E-02 |
| REACTOME_BETA_OXIDATION_OF_VERY_LONG_CHAIN_FATTY_ACIDS | -1.96 | 3.65E-02 |
| REACTOME_PASSIVE_TRANSPORT_BY_AQUAPORINS | -1.95 | 3.86E-02 |
| REACTOME_AKT_PHOSPHORYLATES_TARGETS_IN_THE_NUCLEUS | -1.95 | 3.89E-02 |
| WP_VITAMIN_B6DEPENDENT_AND_RESPONSIVE_DISORDERS | -1.95 | 3.97E-02 |
| REACTOME_POTASSIUM_CHANNELS | -1.94 | 3.96E-02 |
| WP_BIOTIN_METABOLISM_INCLUDING_IMDS | -1.91 | 4.72E-02 |
| REACTOME_NUCLEAR_RECEPTOR_TRANSCRIPTION_PATHWAY | -1.91 | 4.89E-02 |
| WP_VALPROIC_ACID_PATHWAY | -1.90 | 4.89E-02 |

**Supplementary Table S2.** List of differentially altered gene sets in human cardiomyopathy patients (FDR < 0.05).

| **Gene Set Name** | **Normalized Enrichment Score (NES)** | **FDR** |
| --- | --- | --- |
| HALLMARK_MYC_TARGETS_V1 | 5.15 | 0.00E+00 |
| HALLMARK_OXIDATIVE_PHOSPHORYLATION | 4.73 | 0.00E+00 |
| HALLMARK_PROTEIN_SECRETION | 4.23 | 0.00E+00 |
| WP_VEGFAVEGFR2_SIGNALING_PATHWAY | 4.14 | 0.00E+00 |
| REACTOME_SIGNALING_BY_RECEPTOR_TYROSINE_KINASES | 4.04 | 0.00E+00 |
| REACTOME_PROTEIN_LOCALIZATION | 3.98 | 0.00E+00 |
| WP_CILIARY_LANDSCAPE | 3.94 | 0.00E+00 |
| REACTOME_DISEASES_OF_SIGNAL_TRANSDUCTION_BY_GROWTH_FACTOR_RECEPTORS_AND_SECOND_MESSENGERS | 3.70 | 0.00E+00 |
| KEGG_VALINE_LEUCINE_AND_ISOLEUCINE_DEGRADATION | 3.69 | 0.00E+00 |
| HALLMARK_UV_RESPONSE_DN | 3.60 | 0.00E+00 |
| REACTOME_ANTIGEN_PROCESSING_UBIQUITINATION_PROTEASOME_DEGRADATION | 3.56 | 0.00E+00 |
| REACTOME_PYRUVATE_METABOLISM_AND_CITRIC_ACID_TCA_CYCLE | 3.54 | 0.00E+00 |
| HALLMARK_MTORC1_SIGNALING | 3.53 | 0.00E+00 |
| REACTOME_CLASS_I_MHC_MEDIATED_ANTIGEN_PROCESSING_PRESENTATION | 3.52 | 0.00E+00 |
| KEGG_CITRATE_CYCLE_TCA_CYCLE | 3.43 | 0.00E+00 |
| REACTOME_TRANSLATION | 3.42 | 0.00E+00 |
| REACTOME_INTERFERON_SIGNALING | 3.41 | 0.00E+00 |
| HALLMARK_FATTY_ACID_METABOLISM | 3.40 | 0.00E+00 |
| REACTOME_BRANCHED_CHAIN_AMINO_ACID_CATABOLISM | 3.39 | 0.00E+00 |
| REACTOME_PROCESSING_OF_CAPPED_INTRON_CONTAINING_PRE_MRNA | 3.38 | 0.00E+00 |
| PID_FOXO_PATHWAY | 3.38 | 0.00E+00 |
| HALLMARK_INTERFERON_GAMMA_RESPONSE | 3.35 | 0.00E+00 |
| REACTOME_FC_EPSILON_RECEPTOR_FCERI_SIGNALING | 3.34 | 0.00E+00 |
| REACTOME_MRNA_SPLICING | 3.30 | 0.00E+00 |
| REACTOME_SARS_COV_INFECTIONS | 3.29 | 0.00E+00 |
| REACTOME_PEROXISOMAL_PROTEIN_IMPORT | 3.28 | 0.00E+00 |
| WP_EXERCISEINDUCED_CIRCADIAN_REGULATION | 3.27 | 0.00E+00 |
| REACTOME_CELL_CYCLE_MITOTIC | 3.27 | 0.00E+00 |
| HALLMARK_ANDROGEN_RESPONSE | 3.26 | 0.00E+00 |
| REACTOME_HIV_INFECTION | 3.26 | 0.00E+00 |
| REACTOME_INTRA_GOLGI_AND_RETROGRADE_GOLGI_TO_ER_TRAFFIC | 3.26 | 0.00E+00 |
| WP_LEUCINE_ISOLEUCINE_AND_VALINE_METABOLISM | 3.23 | 0.00E+00 |
| REACTOME_ANTIVIRAL_MECHANISM_BY_IFN_STIMULATED_GENES | 3.21 | 0.00E+00 |
| REACTOME_PROTEIN_UBIQUITINATION | 3.21 | 0.00E+00 |
| REACTOME_HSP90_CHAPERONE_CYCLE_FOR_STEROID_HORMONE_RECEPTORS_SHR_IN_THE_PRESENCE_OF_LIGAND | 3.19 | 0.00E+00 |
| REACTOME_CITRIC_ACID_CYCLE_TCA_CYCLE | 3.17 | 0.00E+00 |
| REACTOME_E3_UBIQUITIN_LIGASES_UBIQUITINATE_TARGET_PROTEINS | 3.17 | 0.00E+00 |
| WP_TCA_CYCLE_AKA_KREBS_OR_CITRIC_ACID_CYCLE | 3.15 | 0.00E+00 |
| PID_ERBB1_DOWNSTREAM_PATHWAY | 3.15 | 0.00E+00 |
| REACTOME_AUTOPHAGY | 3.13 | 0.00E+00 |
| HALLMARK_G2M_CHECKPOINT | 3.11 | 0.00E+00 |
| REACTOME_SARS_COV_2_INFECTION | 3.08 | 3.87E-05 |
| WP_AMINO_ACID_METABOLISM | 3.07 | 3.78E-05 |
| WP_CLOCKCONTROLLED_AUTOPHAGY_IN_BONE_METABOLISM | 3.07 | 3.69E-05 |
| REACTOME_TCR_SIGNALING | 3.05 | 3.61E-05 |
| HALLMARK_E2F_TARGETS | 3.04 | 3.53E-05 |
| KEGG_ANTIGEN_PROCESSING_AND_PRESENTATION | 3.03 | 3.46E-05 |
| PID_TELOMERASE_PATHWAY | 3.03 | 3.38E-05 |
| WP_TGFBETA_SIGNALING_PATHWAY | 3.00 | 6.56E-05 |
| REACTOME_FCERI_MEDIATED_MAPK_ACTIVATION | 2.98 | 6.43E-05 |
| HALLMARK_ALLOGRAFT_REJECTION | 2.98 | 6.31E-05 |
| KEGG_TYPE_I_DIABETES_MELLITUS | 2.98 | 6.19E-05 |
| REACTOME_NEDDYLATION | 2.95 | 6.07E-05 |
| REACTOME_ROLE_OF_LAT2_NTAL_LAB_ON_CALCIUM_MOBILIZATION | 2.94 | 5.96E-05 |
| HALLMARK_KRAS_SIGNALING_UP | 2.94 | 5.85E-05 |
| HALLMARK_IL2_STAT5_SIGNALING | 2.92 | 8.58E-05 |
| REACTOME_CELL_CYCLE_CHECKPOINTS | 2.90 | 1.40E-04 |
| REACTOME_CD22_MEDIATED_BCR_REGULATION | 2.89 | 1.38E-04 |
| HALLMARK_TNFA_SIGNALING_VIA_NFKB | 2.88 | 1.63E-04 |
| REACTOME_ANTIGEN_PRESENTATION_FOLDING_ASSEMBLY_AND_PEPTIDE_LOADING_OF_CLASS_I_MHC | 2.88 | 2.38E-04 |
| PID_CD8_TCR_DOWNSTREAM_PATHWAY | 2.85 | 2.61E-04 |
| KEGG_TGF_BETA_SIGNALING_PATHWAY | 2.85 | 2.56E-04 |
| REACTOME_CELLULAR_RESPONSE_TO_HEAT_STRESS | 2.85 | 2.52E-04 |
| REACTOME_BINDING_AND_UPTAKE_OF_LIGANDS_BY_SCAVENGER_RECEPTORS | 2.84 | 2.48E-04 |
| HALLMARK_ADIPOGENESIS | 2.83 | 2.69E-04 |
| REACTOME_FCGR_ACTIVATION | 2.83 | 2.65E-04 |
| REACTOME_THE_CITRIC_ACID_TCA_CYCLE_AND_RESPIRATORY_ELECTRON_TRANSPORT | 2.80 | 3.10E-04 |
| REACTOME_SIGNALING_BY_TGFB_FAMILY_MEMBERS | 2.79 | 3.76E-04 |
| REACTOME_ORGANELLE_BIOGENESIS_AND_MAINTENANCE | 2.79 | 3.94E-04 |
| WP_TRANSLATION_FACTORS | 2.78 | 3.89E-04 |
| REACTOME_GOLGI_TO_ER_RETROGRADE_TRANSPORT | 2.77 | 4.28E-04 |
| KEGG_GRAFT_VERSUS_HOST_DISEASE | 2.76 | 5.33E-04 |
| REACTOME_ANTIGEN_ACTIVATES_B_CELL_RECEPTOR_BCR_LEADING_TO_GENERATION_OF_SECOND_MESSENGERS | 2.76 | 5.47E-04 |
| REACTOME_SIGNALING_BY_TGF_BETA_RECEPTOR_COMPLEX | 2.76 | 5.40E-04 |
| REACTOME_FORMATION_OF_INCISION_COMPLEX_IN_GG_NER | 2.76 | 5.32E-04 |
| WP_TCA_CYCLE_AND_DEFICIENCY_OF_PYRUVATE_DEHYDROGENASE_COMPLEX_PDHC | 2.76 | 5.25E-04 |
| PID_PDGFRB_PATHWAY | 2.76 | 5.19E-04 |
| REACTOME_INTRACELLULAR_SIGNALING_BY_SECOND_MESSENGERS | 2.75 | 5.33E-04 |
| KEGG_PROPANOATE_METABOLISM | 2.73 | 5.87E-04 |
| KEGG_REGULATION_OF_ACTIN_CYTOSKELETON | 2.73 | 5.79E-04 |
| WP_KREBS_CYCLE_DISORDERS | 2.73 | 5.72E-04 |
| KEGG_BUTANOATE_METABOLISM | 2.72 | 6.23E-04 |
| KEGG_WNT_SIGNALING_PATHWAY | 2.72 | 6.35E-04 |
| REACTOME_FCGR3A_MEDIATED_IL10_SYNTHESIS | 2.71 | 6.27E-04 |
| KEGG_BETA_ALANINE_METABOLISM | 2.71 | 6.38E-04 |
| REACTOME_S_PHASE | 2.71 | 6.31E-04 |
| REACTOME_DEUBIQUITINATION | 2.70 | 6.42E-04 |
| BIOCARTA_PLCE_PATHWAY | 2.70 | 6.35E-04 |
| NABA_ECM_GLYCOPROTEINS | 2.69 | 7.17E-04 |
| KEGG_UBIQUITIN_MEDIATED_PROTEOLYSIS | 2.68 | 7.98E-04 |
| NABA_CORE_MATRISOME | 2.68 | 8.07E-04 |
| WP_NSP1_FROM_SARSCOV2_INHIBITS_TRANSLATION_INITIATION_IN_THE_HOST_CELL | 2.68 | 8.50E-04 |
| BIOCARTA_NFAT_PATHWAY | 2.66 | 9.25E-04 |
| REACTOME_FCERI_MEDIATED_CA_2_MOBILIZATION | 2.66 | 9.66E-04 |
| WP_STEROL_REGULATORY_ELEMENTBINDING_PROTEINS_SREBP_SIGNALING | 2.66 | 9.56E-04 |
| REACTOME_EXTRA_NUCLEAR_ESTROGEN_SIGNALING | 2.65 | 1.01E-03 |
| REACTOME_REGULATION_OF_HSF1_MEDIATED_HEAT_SHOCK_RESPONSE | 2.64 | 1.12E-03 |
| REACTOME_HSF1_ACTIVATION | 2.64 | 1.11E-03 |
| REACTOME_TRANSCRIPTIONAL_REGULATION_BY_TP53 | 2.63 | 1.22E-03 |
| BIOCARTA_EIF4_PATHWAY | 2.63 | 1.21E-03 |
| REACTOME_SIGNALING_BY_THE_B_CELL_RECEPTOR_BCR | 2.63 | 1.20E-03 |
| REACTOME_MITOTIC_G1_PHASE_AND_G1_S_TRANSITION | 2.63 | 1.20E-03 |
| REACTOME_COSTIMULATION_BY_THE_CD28_FAMILY | 2.62 | 1.22E-03 |
| REACTOME_HIV_LIFE_CYCLE | 2.62 | 1.21E-03 |
| WP_EGFEGFR_SIGNALING_PATHWAY | 2.62 | 1.24E-03 |
| KEGG_NATURAL_KILLER_CELL_MEDIATED_CYTOTOXICITY | 2.61 | 1.35E-03 |
| REACTOME_FCERI_MEDIATED_NF_KB_ACTIVATION | 2.61 | 1.38E-03 |
| WP_HAIR_FOLLICLE_DEVELOPMENT_CYTODIFFERENTIATION_PART_3_OF_3 | 2.61 | 1.37E-03 |
| REACTOME_GLYCOGEN_SYNTHESIS | 2.60 | 1.50E-03 |
| WP_CIRCADIAN_RHYTHM_GENES | 2.60 | 1.49E-03 |
| REACTOME_SCAVENGING_OF_HEME_FROM_PLASMA | 2.60 | 1.48E-03 |
| REACTOME_NEUTROPHIL_DEGRANULATION | 2.60 | 1.46E-03 |
| KEGG_PATHWAYS_IN_CANCER | 2.60 | 1.45E-03 |
| REACTOME_RAB_GERANYLGERANYLATION | 2.59 | 1.45E-03 |
| WP_MEASLES_VIRUS_INFECTION | 2.59 | 1.45E-03 |
| WP_BRAINDERIVED_NEUROTROPHIC_FACTOR_BDNF_SIGNALING_PATHWAY | 2.59 | 1.44E-03 |
| KEGG_ALLOGRAFT_REJECTION | 2.59 | 1.44E-03 |
| KEGG_CELL_ADHESION_MOLECULES_CAMS | 2.59 | 1.47E-03 |
| PID_AMB2_NEUTROPHILS_PATHWAY | 2.58 | 1.47E-03 |
| REACTOME_CREATION_OF_C4_AND_C2_ACTIVATORS | 2.58 | 1.51E-03 |
| REACTOME_NUCLEOTIDE_EXCISION_REPAIR | 2.58 | 1.54E-03 |
| REACTOME_ROLE_OF_PHOSPHOLIPIDS_IN_PHAGOCYTOSIS | 2.57 | 1.61E-03 |
| PID_MET_PATHWAY | 2.56 | 1.67E-03 |
| BIOCARTA_CTL_PATHWAY | 2.56 | 1.70E-03 |
| REACTOME_SIGNALING_BY_WNT | 2.56 | 1.73E-03 |
| REACTOME_ESR_MEDIATED_SIGNALING | 2.55 | 1.81E-03 |
| REACTOME_NEGATIVE_REGULATION_OF_THE_PI3K_AKT_NETWORK | 2.55 | 1.80E-03 |
| WP_BMP2WNT4FOXO1_PATHWAY_IN_PRIMARY_ENDOMETRIAL_STROMAL_CELL_DIFFERENTIATION | 2.55 | 1.84E-03 |
| KEGG_SPLICEOSOME | 2.54 | 1.98E-03 |
| WP_FOCAL_ADHESION_PI3KAKTMTORSIGNALING_PATHWAY | 2.53 | 2.12E-03 |
| WP_ALZHEIMERS_DISEASE | 2.52 | 2.25E-03 |
| REACTOME_SIGNALING_BY_NUCLEAR_RECEPTORS | 2.52 | 2.25E-03 |
| REACTOME_G_ALPHA_I_SIGNALLING_EVENTS | 2.52 | 2.23E-03 |
| WP_PI3KAKT_SIGNALING_PATHWAY | 2.52 | 2.23E-03 |
| KEGG_COLORECTAL_CANCER | 2.52 | 2.24E-03 |
| REACTOME_PLATELET_ACTIVATION_SIGNALING_AND_AGGREGATION | 2.52 | 2.22E-03 |
| BIOCARTA_CTLA4_PATHWAY | 2.51 | 2.24E-03 |
| BIOCARTA_ALK_PATHWAY | 2.51 | 2.42E-03 |
| BIOCARTA_STATHMIN_PATHWAY | 2.50 | 2.41E-03 |
| BIOCARTA_TCYTOTOXIC_PATHWAY | 2.50 | 2.45E-03 |
| WP_ENDODERM_DIFFERENTIATION | 2.50 | 2.48E-03 |
| BIOCARTA_PRION_PATHWAY | 2.50 | 2.50E-03 |
| WP_EPITHELIAL_TO_MESENCHYMAL_TRANSITION_IN_COLORECTAL_CANCER | 2.50 | 2.51E-03 |
| WP_CHROMOSOMAL_AND_MICROSATELLITE_INSTABILITY_IN_COLORECTAL_CANCER | 2.49 | 2.62E-03 |
| REACTOME_TRANS_GOLGI_NETWORK_VESICLE_BUDDING | 2.48 | 2.75E-03 |
| REACTOME_CELL_SURFACE_INTERACTIONS_AT_THE_VASCULAR_WALL | 2.48 | 2.79E-03 |
| WP_MRNA_PROCESSING | 2.48 | 2.79E-03 |
| WP_OVERVIEW_OF_PROINFLAMMATORY_AND_PROFIBROTIC_MEDIATORS | 2.48 | 2.84E-03 |
| REACTOME_UB_SPECIFIC_PROCESSING_PROTEASES | 2.48 | 2.83E-03 |
| REACTOME_SIGNALING_BY_INTERLEUKINS | 2.47 | 2.87E-03 |
| REACTOME_M_PHASE | 2.47 | 2.85E-03 |
| BIOCARTA_GCR_PATHWAY | 2.47 | 2.83E-03 |
| BIOCARTA_GSK3_PATHWAY | 2.47 | 2.84E-03 |
| REACTOME_EGR2_AND_SOX10_MEDIATED_INITIATION_OF_SCHWANN_CELL_MYELINATION | 2.46 | 3.01E-03 |
| WP_LEPTIN_SIGNALING_PATHWAY | 2.45 | 3.22E-03 |
| HALLMARK_COMPLEMENT | 2.45 | 3.30E-03 |
| REACTOME_FORMATION_OF_TC_NER_PRE_INCISION_COMPLEX | 2.45 | 3.29E-03 |
| WP_ALZHEIMERS_DISEASE_AND_MIRNA_EFFECTS | 2.45 | 3.33E-03 |
| WP_GLYCOGEN_SYNTHESIS_AND_DEGRADATION | 2.44 | 3.44E-03 |
| REACTOME_TGF_BETA_RECEPTOR_SIGNALING_ACTIVATES_SMADS | 2.44 | 3.47E-03 |
| WP_PHYSICOCHEMICAL_FEATURES_AND_TOXICITYASSOCIATED_PATHWAYS | 2.44 | 3.57E-03 |
| REACTOME_GOLGI_ASSOCIATED_VESICLE_BIOGENESIS | 2.44 | 3.56E-03 |
| REACTOME_METABOLISM_OF_AMINO_ACIDS_AND_DERIVATIVES | 2.43 | 3.60E-03 |
| PID_P75_NTR_PATHWAY | 2.43 | 3.63E-03 |
| WP_GASTRIN_SIGNALING_PATHWAY | 2.43 | 3.76E-03 |
| REACTOME_IMMUNOREGULATORY_INTERACTIONS_BETWEEN_A_LYMPHOID_AND_A_NON_LYMPHOID_CELL | 2.42 | 3.94E-03 |
| WP_T_CELL_RECEPTOR_AND_COSTIMULATORY_SIGNALING | 2.42 | 3.99E-03 |
| WP_TCELL_RECEPTOR_SIGNALING_PATHWAY | 2.42 | 4.03E-03 |
| HALLMARK_INTERFERON_ALPHA_RESPONSE | 2.41 | 4.24E-03 |
| WP_MALIGNANT_PLEURAL_MESOTHELIOMA | 2.40 | 4.44E-03 |
| WP_NETWORK_MAP_OF_SARSCOV2_SIGNALING_PATHWAY | 2.40 | 4.69E-03 |
| KEGG_PYRUVATE_METABOLISM | 2.39 | 4.79E-03 |
| REACTOME_TRNA_AMINOACYLATION | 2.39 | 4.85E-03 |
| REACTOME_E2F_MEDIATED_REGULATION_OF_DNA_REPLICATION | 2.39 | 4.83E-03 |
| REACTOME_TRANSLESION_SYNTHESIS_BY_POLH | 2.39 | 4.82E-03 |
| REACTOME_TRANSCRIPTION_COUPLED_NUCLEOTIDE_EXCISION_REPAIR_TC_NER | 2.39 | 4.94E-03 |
| BIOCARTA_TEL_PATHWAY | 2.39 | 4.94E-03 |
| PID_IL12_STAT4_PATHWAY | 2.38 | 5.25E-03 |
| WP_GLIOBLASTOMA_SIGNALING_PATHWAYS | 2.38 | 5.28E-03 |
| REACTOME_MITOTIC_METAPHASE_AND_ANAPHASE | 2.38 | 5.29E-03 |
| PID_NFAT_3PATHWAY | 2.38 | 5.29E-03 |
| REACTOME_NEGATIVE_REGULATORS_OF_DDX58_IFIH1_SIGNALING | 2.38 | 5.31E-03 |
| REACTOME_TCF_DEPENDENT_SIGNALING_IN_RESPONSE_TO_WNT | 2.38 | 5.28E-03 |
| KEGG_PEROXISOME | 2.37 | 5.40E-03 |
| PID_NEPHRIN_NEPH1_PATHWAY | 2.37 | 5.39E-03 |
| REACTOME_MAPK_FAMILY_SIGNALING_CASCADES | 2.37 | 5.50E-03 |
| WP_REGULATION_OF_WNT_BCATENIN_SIGNALING_BY_SMALL_MOLECULE_COMPOUNDS | 2.37 | 5.57E-03 |
| WP_TYPE_II_INTERFERON_SIGNALING | 2.36 | 5.73E-03 |
| REACTOME_EARLY_SARS_COV_2_INFECTION_EVENTS | 2.36 | 5.79E-03 |
| REACTOME_SYNTHESIS_OF_ACTIVE_UBIQUITIN_ROLES_OF_E1_AND_E2_ENZYMES | 2.36 | 5.89E-03 |
| HALLMARK_EPITHELIAL_MESENCHYMAL_TRANSITION | 2.36 | 5.87E-03 |
| HALLMARK_APOPTOSIS | 2.36 | 5.89E-03 |
| PID_MTOR_4PATHWAY | 2.35 | 5.97E-03 |
| REACTOME_CTLA4_INHIBITORY_SIGNALING | 2.35 | 5.99E-03 |
| REACTOME_CLASS_I_PEROXISOMAL_MEMBRANE_PROTEIN_IMPORT | 2.35 | 6.00E-03 |
| BIOCARTA_BAD_PATHWAY | 2.35 | 6.07E-03 |
| BIOCARTA_THELPER_PATHWAY | 2.35 | 6.04E-03 |
| BIOCARTA_EIF_PATHWAY | 2.35 | 6.02E-03 |
| REACTOME_MITOCHONDRIAL_TRNA_AMINOACYLATION | 2.35 | 6.00E-03 |
| REACTOME_GLOBAL_GENOME_NUCLEOTIDE_EXCISION_REPAIR_GG_NER | 2.35 | 6.14E-03 |
| REACTOME_TRANSCRIPTION_OF_THE_HIV_GENOME | 2.34 | 6.14E-03 |
| KEGG_VIRAL_MYOCARDITIS | 2.34 | 6.18E-03 |
| REACTOME_SUMO_IS_CONJUGATED_TO_E1_UBA2_SAE1 | 2.34 | 6.22E-03 |
| REACTOME_G2_M_CHECKPOINTS | 2.34 | 6.23E-03 |
| REACTOME_MITOCHONDRIAL_TRANSLATION | 2.34 | 6.23E-03 |
| HALLMARK_PEROXISOME | 2.34 | 6.21E-03 |
| WP_ANDROGEN_RECEPTOR_SIGNALING_PATHWAY | 2.34 | 6.25E-03 |
| REACTOME_MRNA_CAPPING | 2.34 | 6.25E-03 |
| BIOCARTA_CREB_PATHWAY | 2.34 | 6.23E-03 |
| WP_EBOLA_VIRUS_INFECTION_IN_HOST | 2.33 | 6.41E-03 |
| REACTOME_SIGNALING_BY_WNT_IN_CANCER | 2.33 | 6.51E-03 |
| KEGG_PROTEIN_EXPORT | 2.33 | 6.57E-03 |
| BIOCARTA_PAR1_PATHWAY | 2.32 | 6.76E-03 |
| REACTOME_DUAL_INCISION_IN_TC_NER | 2.32 | 6.79E-03 |
| REACTOME_PYRUVATE_METABOLISM | 2.32 | 7.03E-03 |
| REACTOME_MHC_CLASS_II_ANTIGEN_PRESENTATION | 2.32 | 7.01E-03 |
| REACTOME_DISASSEMBLY_OF_THE_DESTRUCTION_COMPLEX_AND_RECRUITMENT_OF_AXIN_TO_THE_MEMBRANE | 2.32 | 6.97E-03 |
| BIOCARTA_IGF1MTOR_PATHWAY | 2.31 | 7.06E-03 |
| REACTOME_METABOLISM_OF_COFACTORS | 2.31 | 7.03E-03 |
| BIOCARTA_CSK_PATHWAY | 2.31 | 7.16E-03 |
| REACTOME_METABOLISM_OF_VITAMINS_AND_COFACTORS | 2.31 | 7.14E-03 |
| HALLMARK_UNFOLDED_PROTEIN_RESPONSE | 2.30 | 7.58E-03 |
| REACTOME_DEADENYLATION_DEPENDENT_MRNA_DECAY | 2.30 | 7.59E-03 |
| REACTOME_PROGRAMMED_CELL_DEATH | 2.30 | 7.63E-03 |
| KEGG_T_CELL_RECEPTOR_SIGNALING_PATHWAY | 2.30 | 7.66E-03 |
| WP_EMBRYONIC_STEM_CELL_PLURIPOTENCY_PATHWAYS | 2.30 | 7.63E-03 |
| PID_AR_TF_PATHWAY | 2.30 | 7.69E-03 |
| WP_DEVELOPMENT_OF_URETERIC_COLLECTION_SYSTEM | 2.30 | 7.67E-03 |
| WP_PHOTODYNAMIC_THERAPYINDUCED_UNFOLDED_PROTEIN_RESPONSE | 2.30 | 7.64E-03 |
| REACTOME_RNA_POLYMERASE_II_PRE_TRANSCRIPTION_EVENTS | 2.29 | 7.84E-03 |
| REACTOME_MAPK_TARGETS_NUCLEAR_EVENTS_MEDIATED_BY_MAP_KINASES | 2.29 | 7.86E-03 |
| BIOCARTA_MTOR_PATHWAY | 2.29 | 7.86E-03 |
| PID_TGFBR_PATHWAY | 2.29 | 7.94E-03 |
| REACTOME_HIV_TRANSCRIPTION_INITIATION | 2.28 | 8.28E-03 |
| WP_TRANSLATION_INHIBITORS_IN_CHRONICALLY_ACTIVATED_PDGFRA_CELLS | 2.27 | 8.81E-03 |
| KEGG_AUTOIMMUNE_THYROID_DISEASE | 2.27 | 8.87E-03 |
| REACTOME_PIWI_INTERACTING_RNA_PIRNA_BIOGENESIS | 2.27 | 9.02E-03 |
| REACTOME_CLEC7A_DECTIN_1_SIGNALING | 2.27 | 9.02E-03 |
| REACTOME_DEADENYLATION_OF_MRNA | 2.27 | 9.00E-03 |
| WP_RETINOBLASTOMA_GENE_IN_CANCER | 2.26 | 9.03E-03 |
| WP_7Q1123_COPY_NUMBER_VARIATION_SYNDROME | 2.26 | 9.26E-03 |
| REACTOME_INTERFERON_ALPHA_BETA_SIGNALING | 2.25 | 9.60E-03 |
| PID_IL2_PI3K_PATHWAY | 2.25 | 9.71E-03 |
| REACTOME_SARS_COV_2_TARGETS_HOST_INTRACELLULAR_SIGNALLING_AND_REGULATORY_PATHWAYS | 2.25 | 9.71E-03 |
| REACTOME_CILIUM_ASSEMBLY | 2.25 | 9.85E-03 |
| REACTOME_SIGNALING_BY_INSULIN_RECEPTOR | 2.24 | 1.01E-02 |
| BIOCARTA_LYMPHOCYTE_PATHWAY | 2.24 | 1.03E-02 |
| REACTOME_SARS_COV_1_INFECTION | 2.24 | 1.03E-02 |
| REACTOME_MYD88_INDEPENDENT_TLR4_CASCADE | 2.24 | 1.04E-02 |
| BIOCARTA_PPARA_PATHWAY | 2.23 | 1.05E-02 |
| REACTOME_OAS_ANTIVIRAL_RESPONSE | 2.23 | 1.06E-02 |
| REACTOME_MITOTIC_SPINDLE_CHECKPOINT | 2.23 | 1.09E-02 |
| WP_G1_TO_S_CELL_CYCLE_CONTROL | 2.22 | 1.12E-02 |
| HALLMARK_GLYCOLYSIS | 2.22 | 1.13E-02 |
| REACTOME_HOST_INTERACTIONS_OF_HIV_FACTORS | 2.22 | 1.13E-02 |
| REACTOME_PEROXISOMAL_LIPID_METABOLISM | 2.22 | 1.13E-02 |
| WP_SPLICING_FACTOR_NOVA_REGULATED_SYNAPTIC_PROTEINS | 2.22 | 1.13E-02 |
| REACTOME_CIRCADIAN_CLOCK | 2.22 | 1.15E-02 |
| KEGG_CELL_CYCLE | 2.22 | 1.15E-02 |
| REACTOME_PARASITE_INFECTION | 2.21 | 1.16E-02 |
| REACTOME_SIGNALING_BY_MET | 2.21 | 1.16E-02 |
| REACTOME_MTOR_SIGNALLING | 2.21 | 1.16E-02 |
| REACTOME_MITOTIC_PROMETAPHASE | 2.21 | 1.18E-02 |
| REACTOME_APOPTOSIS | 2.21 | 1.18E-02 |
| WP_LIPID_METABOLISM_PATHWAY | 2.21 | 1.20E-02 |
| PID_INTEGRIN5_PATHWAY | 2.20 | 1.21E-02 |
| WP_HEPATITIS_B_INFECTION | 2.20 | 1.21E-02 |
| REACTOME_RESOLUTION_OF_SISTER_CHROMATID_COHESION | 2.20 | 1.22E-02 |
| REACTOME_TOLL_LIKE_RECEPTOR_CASCADES | 2.20 | 1.24E-02 |
| REACTOME_MITOCHONDRIAL_BIOGENESIS | 2.20 | 1.25E-02 |
| REACTOME_SIGNALING_BY_FGFR | 2.20 | 1.24E-02 |
| REACTOME_TRANSPORT_TO_THE_GOLGI_AND_SUBSEQUENT_MODIFICATION | 2.20 | 1.25E-02 |
| REACTOME_SELECTIVE_AUTOPHAGY | 2.20 | 1.25E-02 |
| WP_WNT_SIGNALING | 2.19 | 1.25E-02 |
| REACTOME_TRIGLYCERIDE_CATABOLISM | 2.19 | 1.32E-02 |
| WP_CELL_CYCLE | 2.18 | 1.36E-02 |
| PID_S1P_S1P2_PATHWAY | 2.18 | 1.35E-02 |
| REACTOME_INTERFERON_GAMMA_SIGNALING | 2.18 | 1.36E-02 |
| WP_BMP_SIGNALING_IN_EYELID_DEVELOPMENT | 2.18 | 1.36E-02 |
| KEGG_TRYPTOPHAN_METABOLISM | 2.18 | 1.37E-02 |
| WP_WNT_SIGNALING_PATHWAY_AND_PLURIPOTENCY | 2.18 | 1.36E-02 |
| PID_ATF2_PATHWAY | 2.18 | 1.37E-02 |
| REACTOME_SIGNALING_BY_ALK_IN_CANCER | 2.17 | 1.42E-02 |
| WP_TCELL_ANTIGEN_RECEPTOR_TCR_PATHWAY_DURING_STAPHYLOCOCCUS_AUREUS_INFECTION | 2.17 | 1.42E-02 |
| REACTOME_TOLL_LIKE_RECEPTOR_9_TLR9_CASCADE | 2.17 | 1.43E-02 |
| WP_THYROID_STIMULATING_HORMONE_TSH_SIGNALING_PATHWAY | 2.17 | 1.43E-02 |
| WP_OSTEOBLAST_DIFFERENTIATION_AND_RELATED_DISEASES | 2.17 | 1.44E-02 |
| REACTOME_MET_ACTIVATES_PI3K_AKT_SIGNALING | 2.17 | 1.43E-02 |
| REACTOME_ASPARAGINE_N_LINKED_GLYCOSYLATION | 2.17 | 1.44E-02 |
| KEGG_NEUROTROPHIN_SIGNALING_PATHWAY | 2.17 | 1.45E-02 |
| HALLMARK_BILE_ACID_METABOLISM | 2.16 | 1.46E-02 |
| PID_IL12_2PATHWAY | 2.16 | 1.49E-02 |
| WP_ENDOPLASMIC_RETICULUM_STRESS_RESPONSE_IN_CORONAVIRUS_INFECTION | 2.16 | 1.50E-02 |
| KEGG_RNA_DEGRADATION | 2.16 | 1.51E-02 |
| PID_IL2_1PATHWAY | 2.16 | 1.52E-02 |
| REACTOME_SARS_COV_2_HOST_INTERACTIONS | 2.16 | 1.52E-02 |
| REACTOME_SYNTHESIS_SECRETION_AND_INACTIVATION_OF_GLUCAGON_LIKE_PEPTIDE_1_GLP_1 | 2.16 | 1.53E-02 |
| REACTOME_RHO_GTPASE_EFFECTORS | 2.15 | 1.56E-02 |
| WP_PATHOGENESIS_OF_SARSCOV2_MEDIATED_BY_NSP9NSP10_COMPLEX | 2.14 | 1.66E-02 |
| REACTOME_FOXO_MEDIATED_TRANSCRIPTION | 2.13 | 1.74E-02 |
| REACTOME_NETRIN_1_SIGNALING | 2.13 | 1.74E-02 |
| REACTOME_IRON_UPTAKE_AND_TRANSPORT | 2.13 | 1.74E-02 |
| BIOCARTA_CK1_PATHWAY | 2.13 | 1.80E-02 |
| REACTOME_REGULATION_OF_PLK1_ACTIVITY_AT_G2_M_TRANSITION | 2.13 | 1.80E-02 |
| REACTOME_SIGNALING_BY_FGFR2 | 2.13 | 1.81E-02 |
| WP_MITOCHONDRIAL_COMPLEX_II_ASSEMBLY | 2.12 | 1.84E-02 |
| REACTOME_METABOLISM_OF_CARBOHYDRATES | 2.12 | 1.87E-02 |
| PID_WNT_SIGNALING_PATHWAY | 2.12 | 1.87E-02 |
| WP_CALCIUM_REGULATION_IN_CARDIAC_CELLS | 2.12 | 1.86E-02 |
| BIOCARTA_TCRA_PATHWAY | 2.12 | 1.87E-02 |
| PID_BETA_CATENIN_NUC_PATHWAY | 2.12 | 1.88E-02 |
| REACTOME_REGULATION_OF_MRNA_STABILITY_BY_PROTEINS_THAT_BIND_AU_RICH_ELEMENTS | 2.12 | 1.91E-02 |
| REACTOME_TRANSCRIPTIONAL_ACTIVITY_OF_SMAD2_SMAD3_SMAD4_HETEROTRIMER | 2.11 | 1.93E-02 |
| REACTOME_CARGO_TRAFFICKING_TO_THE_PERICILIARY_MEMBRANE | 2.11 | 1.94E-02 |
| REACTOME_COPI_MEDIATED_ANTEROGRADE_TRANSPORT | 2.11 | 1.95E-02 |
| WP_REGULATION_OF_ACTIN_CYTOSKELETON | 2.11 | 1.94E-02 |
| REACTOME_DNA_REPAIR | 2.11 | 1.94E-02 |
| REACTOME_GENERATION_OF_SECOND_MESSENGER_MOLECULES | 2.11 | 1.94E-02 |
| REACTOME_REGULATION_OF_PTEN_STABILITY_AND_ACTIVITY | 2.11 | 1.97E-02 |
| BIOCARTA_MONOCYTE_PATHWAY | 2.10 | 2.01E-02 |
| BIOCARTA_IGF1R_PATHWAY | 2.10 | 2.01E-02 |
| REACTOME_CHK1_CHK2_CDS1_MEDIATED_INACTIVATION_OF_CYCLIN_B_CDK1_COMPLEX | 2.10 | 2.01E-02 |
| WP_WNT_SIGNALING_PATHWAY | 2.10 | 2.03E-02 |
| BIOCARTA_TCAPOPTOSIS_PATHWAY | 2.10 | 2.05E-02 |
| WP_ALLOGRAFT_REJECTION | 2.10 | 2.08E-02 |
| BIOCARTA_CCR5_PATHWAY | 2.09 | 2.08E-02 |
| PID_HIF1_TFPATHWAY | 2.09 | 2.08E-02 |
| REACTOME_ENDOSOMAL_VACUOLAR_PATHWAY | 2.09 | 2.09E-02 |
| BIOCARTA_SUMO_PATHWAY | 2.09 | 2.09E-02 |
| WP_SUDDEN_INFANT_DEATH_SYNDROME_SIDS_SUSCEPTIBILITY_PATHWAYS | 2.09 | 2.10E-02 |
| REACTOME_ACTIVATION_OF_BH3_ONLY_PROTEINS | 2.09 | 2.09E-02 |
| REACTOME_MTORC1_MEDIATED_SIGNALLING | 2.09 | 2.11E-02 |
| REACTOME_SIGNALING_BY_ERBB4 | 2.09 | 2.11E-02 |
| HALLMARK_NOTCH_SIGNALING | 2.09 | 2.12E-02 |
| WP_GENES_RELATED_TO_PRIMARY_CILIUM_DEVELOPMENT_BASED_ON_CRISPR | 2.09 | 2.13E-02 |
| REACTOME_SPHINGOLIPID_DE_NOVO_BIOSYNTHESIS | 2.09 | 2.13E-02 |
| REACTOME_RETROGRADE_TRANSPORT_AT_THE_TRANS_GOLGI_NETWORK | 2.08 | 2.16E-02 |
| REACTOME_G2_M_DNA_DAMAGE_CHECKPOINT | 2.08 | 2.17E-02 |
| PID_THROMBIN_PAR4_PATHWAY | 2.08 | 2.19E-02 |
| REACTOME_DNA_STRAND_ELONGATION | 2.08 | 2.20E-02 |
| PID_AR_PATHWAY | 2.08 | 2.24E-02 |
| REACTOME_OPIOID_SIGNALLING | 2.07 | 2.25E-02 |
| PID_BCR_5PATHWAY | 2.07 | 2.26E-02 |
| REACTOME_SIGNAL_AMPLIFICATION | 2.07 | 2.26E-02 |
| REACTOME_ACTIVATION_OF_THE_MRNA_UPON_BINDING_OF_THE_CAP_BINDING_COMPLEX_AND_EIFS_AND_SUBSEQUENT_BINDING_TO_43S | 2.07 | 2.26E-02 |
| REACTOME_SIGNALING_BY_HIPPO | 2.07 | 2.26E-02 |
| REACTOME_PEPTIDE_HORMONE_METABOLISM | 2.07 | 2.28E-02 |
| PID_IL1_PATHWAY | 2.07 | 2.28E-02 |
| REACTOME_SEPARATION_OF_SISTER_CHROMATIDS | 2.07 | 2.27E-02 |
| REACTOME_RAC2_GTPASE_CYCLE | 2.07 | 2.29E-02 |
| REACTOME_SIGNALING_BY_ERBB2 | 2.06 | 2.35E-02 |
| HALLMARK_PI3K_AKT_MTOR_SIGNALING | 2.06 | 2.35E-02 |
| REACTOME_SIGNALING_BY_RNF43_MUTANTS | 2.06 | 2.37E-02 |
| PID_AURORA_B_PATHWAY | 2.06 | 2.37E-02 |
| HALLMARK_CHOLESTEROL_HOMEOSTASIS | 2.06 | 2.37E-02 |
| REACTOME_BETA_CATENIN_PHOSPHORYLATION_CASCADE | 2.06 | 2.38E-02 |
| REACTOME_PI_3K_CASCADE_FGFR4 | 2.05 | 2.46E-02 |
| REACTOME_COPI_INDEPENDENT_GOLGI_TO_ER_RETROGRADE_TRAFFIC | 2.05 | 2.47E-02 |
| WP_MAMMARY_GLAND_DEVELOPMENT_PATHWAY_EMBRYONIC_DEVELOPMENT_STAGE_1_OF_4 | 2.05 | 2.49E-02 |
| REACTOME_MEIOTIC_RECOMBINATION | 2.05 | 2.49E-02 |
| WP_NETRINUNC5B_SIGNALING_PATHWAY | 2.05 | 2.50E-02 |
| WP_PARKINUBIQUITIN_PROTEASOMAL_SYSTEM_PATHWAY | 2.05 | 2.52E-02 |
| REACTOME_DOWNREGULATION_OF_SMAD2_3_SMAD4_TRANSCRIPTIONAL_ACTIVITY | 2.05 | 2.51E-02 |
| REACTOME_AMYLOID_FIBER_FORMATION | 2.05 | 2.51E-02 |
| WP_SARSCOV2_INNATE_IMMUNITY_EVASION_AND_CELLSPECIFIC_IMMUNE_RESPONSE | 2.05 | 2.50E-02 |
| WP_MBDNF_AND_PROBDNF_REGULATION_OF_GABA_NEUROTRANSMISSION | 2.05 | 2.52E-02 |
| WP_NAD_METABOLISM_IN_ONCOGENEINDUCED_SENESCENCE_AND_MITOCHONDRIAL_DYSFUNCTIONASSOCIATED_SENESCENCE | 2.04 | 2.52E-02 |
| PID_LYMPH_ANGIOGENESIS_PATHWAY | 2.04 | 2.66E-02 |
| PID_HDAC_CLASSII_PATHWAY | 2.03 | 2.69E-02 |
| REACTOME_SIGNALING_BY_FGFR2_IIIA_TM | 2.03 | 2.69E-02 |
| WP_DEVELOPMENT_AND_HETEROGENEITY_OF_THE_ILC_FAMILY | 2.03 | 2.72E-02 |
| PID_FGF_PATHWAY | 2.03 | 2.77E-02 |
| REACTOME_RND3_GTPASE_CYCLE | 2.03 | 2.77E-02 |
| REACTOME_COOPERATION_OF_PDCL_PHLP1_AND_TRIC_CCT_IN_G_PROTEIN_BETA_FOLDING | 2.03 | 2.76E-02 |
| WP_EXTRAFOLLICULAR_B_CELL_ACTIVATION_BY_SARSCOV2 | 2.03 | 2.76E-02 |
| REACTOME_TRANSLATION_OF_REPLICASE_AND_ASSEMBLY_OF_THE_REPLICATION_TRANSCRIPTION_COMPLEX | 2.02 | 2.84E-02 |
| BIOCARTA_TCR_PATHWAY | 2.02 | 2.86E-02 |
| WP_DNA_DAMAGE_RESPONSE_ONLY_ATM_DEPENDENT | 2.02 | 2.86E-02 |
| REACTOME_BBSOME_MEDIATED_CARGO_TARGETING_TO_CILIUM | 2.02 | 2.87E-02 |
| REACTOME_PINK1_PRKN_MEDIATED_MITOPHAGY | 2.01 | 2.91E-02 |
| REACTOME_INTERLEUKIN_17_SIGNALING | 2.01 | 2.92E-02 |
| PID_CD8_TCR_PATHWAY | 2.01 | 2.92E-02 |
| WP_BREAST_CANCER_PATHWAY | 2.01 | 2.93E-02 |
| KEGG_LIMONENE_AND_PINENE_DEGRADATION | 2.01 | 2.94E-02 |
| REACTOME_ACTIVATED_NTRK2_SIGNALS_THROUGH_PI3K | 2.01 | 2.95E-02 |
| REACTOME_PTEN_REGULATION | 2.01 | 2.97E-02 |
| BIOCARTA_NO1_PATHWAY | 2.01 | 2.98E-02 |
| REACTOME_CHEMOKINE_RECEPTORS_BIND_CHEMOKINES | 2.01 | 2.98E-02 |
| BIOCARTA_PTEN_PATHWAY | 2.01 | 3.00E-02 |
| KEGG_EPITHELIAL_CELL_SIGNALING_IN_HELICOBACTER_PYLORI_INFECTION | 2.00 | 3.01E-02 |
| REACTOME_ATTENUATION_PHASE | 2.00 | 3.01E-02 |
| REACTOME_HCMV_INFECTION | 2.00 | 3.01E-02 |
| WP_B_CELL_RECEPTOR_SIGNALING_PATHWAY | 2.00 | 3.00E-02 |
| KEGG_PANCREATIC_CANCER | 2.00 | 3.03E-02 |
| REACTOME_RHO_GTPASES_ACTIVATE_PKNS | 2.00 | 3.03E-02 |
| KEGG_TOLL_LIKE_RECEPTOR_SIGNALING_PATHWAY | 2.00 | 3.03E-02 |
| REACTOME_ONCOGENIC_MAPK_SIGNALING | 2.00 | 3.04E-02 |
| REACTOME_MET_ACTIVATES_PTPN11 | 2.00 | 3.05E-02 |
| WP_DYRK1A | 2.00 | 3.06E-02 |
| WP_VASOPRESSINREGULATED_WATER_REABSORPTION | 2.00 | 3.08E-02 |
| REACTOME_REELIN_SIGNALLING_PATHWAY | 2.00 | 3.08E-02 |
| KEGG_NUCLEOTIDE_EXCISION_REPAIR | 1.99 | 3.09E-02 |
| BIOCARTA_ETC_PATHWAY | 1.99 | 3.09E-02 |
| REACTOME_RAB_GEFS_EXCHANGE_GTP_FOR_GDP_ON_RABS | 1.99 | 3.10E-02 |
| WP_GLYCOLYSIS_AND_GLUCONEOGENESIS | 1.99 | 3.13E-02 |
| REACTOME_FCGAMMA_RECEPTOR_FCGR_DEPENDENT_PHAGOCYTOSIS | 1.99 | 3.14E-02 |
| REACTOME_CHROMOSOME_MAINTENANCE | 1.99 | 3.16E-02 |
| REACTOME_CLATHRIN_MEDIATED_ENDOCYTOSIS | 1.99 | 3.17E-02 |
| REACTOME_PP2A_MEDIATED_DEPHOSPHORYLATION_OF_KEY_METABOLIC_FACTORS | 1.99 | 3.17E-02 |
| REACTOME_CYTOCHROME_C_MEDIATED_APOPTOTIC_RESPONSE | 1.99 | 3.17E-02 |
| WP_METABOLIC_REPROGRAMMING_IN_PANCREATIC_CANCER | 1.99 | 3.17E-02 |
| WP_HEAD_AND_NECK_SQUAMOUS_CELL_CARCINOMA | 1.99 | 3.18E-02 |
| REACTOME_CYCLIN_A_CDK2_ASSOCIATED_EVENTS_AT_S_PHASE_ENTRY | 1.99 | 3.20E-02 |
| WP_TGFBETA_RECEPTOR_SIGNALING | 1.99 | 3.19E-02 |
| BIOCARTA_KREB_PATHWAY | 1.98 | 3.24E-02 |
| WP_BDNFTRKB_SIGNALING | 1.98 | 3.26E-02 |
| REACTOME_IRS_MEDIATED_SIGNALLING | 1.98 | 3.28E-02 |
| REACTOME_ER_TO_GOLGI_ANTEROGRADE_TRANSPORT | 1.98 | 3.30E-02 |
| WP_MECP2_AND_ASSOCIATED_RETT_SYNDROME | 1.98 | 3.32E-02 |
| REACTOME_RHOBTB1_GTPASE_CYCLE | 1.98 | 3.33E-02 |
| BIOCARTA_EDG1_PATHWAY | 1.97 | 3.34E-02 |
| REACTOME_SIGNALING_BY_FGFR4 | 1.97 | 3.37E-02 |
| REACTOME_INHIBITION_OF_REPLICATION_INITIATION_OF_DAMAGED_DNA_BY_RB1_E2F1 | 1.97 | 3.39E-02 |
| KEGG_VASOPRESSIN_REGULATED_WATER_REABSORPTION | 1.97 | 3.41E-02 |
| REACTOME_POTENTIAL_THERAPEUTICS_FOR_SARS | 1.97 | 3.41E-02 |
| BIOCARTA_RAB_PATHWAY | 1.97 | 3.42E-02 |
| REACTOME_RHO_GTPASE_CYCLE | 1.97 | 3.43E-02 |
| KEGG_ALANINE_ASPARTATE_AND_GLUTAMATE_METABOLISM | 1.97 | 3.42E-02 |
| PID_VEGFR1_PATHWAY | 1.97 | 3.42E-02 |
| REACTOME_SIGNALING_BY_VEGF | 1.97 | 3.42E-02 |
| WP_3Q29_COPY_NUMBER_VARIATION_SYNDROME | 1.97 | 3.42E-02 |
| REACTOME_ENERGY_DEPENDENT_REGULATION_OF_MTOR_BY_LKB1_AMPK | 1.97 | 3.42E-02 |
| KEGG_INSULIN_SIGNALING_PATHWAY | 1.97 | 3.42E-02 |
| REACTOME_REGULATION_OF_CHOLESTEROL_BIOSYNTHESIS_BY_SREBP_SREBF | 1.97 | 3.42E-02 |
| REACTOME_METHIONINE_SALVAGE_PATHWAY | 1.97 | 3.42E-02 |
| WP_MESODERMAL_COMMITMENT_PATHWAY | 1.96 | 3.60E-02 |
| WP_DISRUPTION_OF_POSTSYNAPTIC_SIGNALING_BY_CNV | 1.96 | 3.59E-02 |
| KEGG_REGULATION_OF_AUTOPHAGY | 1.96 | 3.59E-02 |
| REACTOME_DNA_REPLICATION | 1.96 | 3.59E-02 |
| WP_2586_ARYL_HYDROCARBON_RECEPTOR_PATHWAY | 1.95 | 3.61E-02 |
| REACTOME_INCRETIN_SYNTHESIS_SECRETION_AND_INACTIVATION | 1.95 | 3.73E-02 |
| REACTOME_SARS_COV_2_ACTIVATES_MODULATES_INNATE_AND_ADAPTIVE_IMMUNE_RESPONSES | 1.95 | 3.75E-02 |
| KEGG_FATTY_ACID_METABOLISM | 1.95 | 3.75E-02 |
| REACTOME_REGULATION_OF_TP53_ACTIVITY_THROUGH_PHOSPHORYLATION | 1.95 | 3.79E-02 |
| WP_NEOVASCULARISATION_PROCESSES | 1.94 | 3.78E-02 |
| WP_AUTOPHAGY | 1.94 | 3.80E-02 |
| KEGG_OOCYTE_MEIOSIS | 1.94 | 3.81E-02 |
| REACTOME_NEGATIVE_REGULATION_OF_MAPK_PATHWAY | 1.94 | 3.80E-02 |
| REACTOME_CONSTITUTIVE_SIGNALING_BY_ABERRANT_PI3K_IN_CANCER | 1.94 | 3.80E-02 |
| PID_BMP_PATHWAY | 1.94 | 3.86E-02 |
| PID_AVB3_INTEGRIN_PATHWAY | 1.94 | 3.87E-02 |
| WP_ENDOCHONDRAL_OSSIFICATION_WITH_SKELETAL_DYSPLASIAS | 1.94 | 3.86E-02 |
| WP_TOLLLIKE_RECEPTOR_SIGNALING_PATHWAY | 1.94 | 3.86E-02 |
| PID_ATR_PATHWAY | 1.94 | 3.86E-02 |
| WP_APOPTOSISRELATED_NETWORK_DUE_TO_ALTERED_NOTCH3_IN_OVARIAN_CANCER | 1.94 | 3.88E-02 |
| BIOCARTA_CHREBP_PATHWAY | 1.94 | 3.88E-02 |
| WP_NAD_METABOLISM_SIRTUINS_AND_AGING | 1.94 | 3.88E-02 |
| REACTOME_TRANSLOCATION_OF_SLC2A4_GLUT4_TO_THE_PLASMA_MEMBRANE | 1.94 | 3.89E-02 |
| PID_EPO_PATHWAY | 1.94 | 3.88E-02 |
| KEGG_ARGININE_AND_PROLINE_METABOLISM | 1.94 | 3.87E-02 |
| PID_ERBB1_INTERNALIZATION_PATHWAY | 1.94 | 3.88E-02 |
| REACTOME_SIGNALING_BY_TYPE_1_INSULIN_LIKE_GROWTH_FACTOR_1_RECEPTOR_IGF1R | 1.93 | 3.88E-02 |
| HALLMARK_HEME_METABOLISM | 1.93 | 3.87E-02 |
| REACTOME_REGULATION_OF_FZD_BY_UBIQUITINATION | 1.93 | 3.94E-02 |
| PID_NFAT_TFPATHWAY | 1.93 | 3.94E-02 |
| WP_TH17_CELL_DIFFERENTIATION_PATHWAY | 1.93 | 4.03E-02 |
| PID_CXCR4_PATHWAY | 1.93 | 4.04E-02 |
| REACTOME_INTEGRATION_OF_ENERGY_METABOLISM | 1.93 | 4.04E-02 |
| BIOCARTA_TFF_PATHWAY | 1.93 | 4.05E-02 |
| REACTOME_THROMBOXANE_SIGNALLING_THROUGH_TP_RECEPTOR | 1.92 | 4.05E-02 |
| BIOCARTA_IL17_PATHWAY | 1.92 | 4.05E-02 |
| PID_FAK_PATHWAY | 1.92 | 4.05E-02 |
| WP_ENDOCHONDRAL_OSSIFICATION | 1.92 | 4.06E-02 |
| WP_ACUTE_VIRAL_MYOCARDITIS | 1.92 | 4.17E-02 |
| REACTOME_SIGNALING_BY_CTNNB1_PHOSPHO_SITE_MUTANTS | 1.92 | 4.19E-02 |
| WP_RANKLRANK_SIGNALING_PATHWAY | 1.92 | 4.19E-02 |
| REACTOME_MITOPHAGY | 1.92 | 4.21E-02 |
| PID_ARF6_TRAFFICKING_PATHWAY | 1.92 | 4.21E-02 |
| REACTOME_MRNA_DECAY_BY_5_TO_3_EXORIBONUCLEASE | 1.92 | 4.21E-02 |
| REACTOME_DOWNSTREAM_SIGNALING_EVENTS_OF_B_CELL_RECEPTOR_BCR | 1.92 | 4.22E-02 |
| REACTOME_BETA_OXIDATION_OF_DECANOYL_COA_TO_OCTANOYL_COA_COA | 1.92 | 4.21E-02 |
| PID_ERBB4_PATHWAY | 1.91 | 4.20E-02 |
| REACTOME_INSULIN_PROCESSING | 1.91 | 4.22E-02 |
| KEGG_B_CELL_RECEPTOR_SIGNALING_PATHWAY | 1.91 | 4.24E-02 |
| WP_PANCREATIC_ADENOCARCINOMA_PATHWAY | 1.91 | 4.25E-02 |
| WP_TGFBETA_RECEPTOR_SIGNALING_IN_SKELETAL_DYSPLASIAS | 1.91 | 4.28E-02 |
| REACTOME_PEXOPHAGY | 1.91 | 4.29E-02 |
| REACTOME_FORMATION_OF_RNA_POL_II_ELONGATION_COMPLEX | 1.91 | 4.34E-02 |
| KEGG_FOCAL_ADHESION | 1.91 | 4.34E-02 |
| WP_CHOLESTEROL_METABOLISM_WITH_BLOCH_AND_KANDUTSCHRUSSELL_PATHWAYS | 1.91 | 4.33E-02 |
| REACTOME_THROMBIN_SIGNALLING_THROUGH_PROTEINASE_ACTIVATED_RECEPTORS_PARS | 1.90 | 4.37E-02 |
| PID_IL5_PATHWAY | 1.90 | 4.36E-02 |
| REACTOME_VEGFR2_MEDIATED_VASCULAR_PERMEABILITY | 1.90 | 4.35E-02 |
| KEGG_ARRHYTHMOGENIC_RIGHT_VENTRICULAR_CARDIOMYOPATHY_ARVC | 1.90 | 4.36E-02 |
| REACTOME_SIGNALING_BY_FGFR_IN_DISEASE | 1.90 | 4.37E-02 |
| REACTOME_ESTROGEN_DEPENDENT_NUCLEAR_EVENTS_DOWNSTREAM_OF_ESR_MEMBRANE_SIGNALING | 1.90 | 4.37E-02 |
| PID_IL6_7_PATHWAY | 1.90 | 4.41E-02 |
| WP_SPHINGOLIPID_METABOLISM_IN_SENESCENCE | 1.90 | 4.42E-02 |
| REACTOME_ENDOSOMAL_SORTING_COMPLEX_REQUIRED_FOR_TRANSPORT_ESCRT | 1.90 | 4.43E-02 |
| REACTOME_MET_ACTIVATES_RAP1_AND_RAC1 | 1.90 | 4.43E-02 |
| PID_RET_PATHWAY | 1.90 | 4.43E-02 |
| REACTOME_SIGNALING_BY_GPCR | 1.90 | 4.42E-02 |
| PID_ANGIOPOIETIN_RECEPTOR_PATHWAY | 1.90 | 4.42E-02 |
| REACTOME_TP53_REGULATES_METABOLIC_GENES | 1.90 | 4.43E-02 |
| REACTOME_DNA_REPLICATION_PRE_INITIATION | 1.90 | 4.44E-02 |
| REACTOME_SIGNALING_BY_FGFR2_IN_DISEASE | 1.90 | 4.44E-02 |
| BIOCARTA_CELLCYCLE_PATHWAY | 1.90 | 4.44E-02 |
| REACTOME_SPHINGOLIPID_METABOLISM | 1.90 | 4.44E-02 |
| PID_INTEGRIN_A4B1_PATHWAY | 1.89 | 4.48E-02 |
| WP_IL6_SIGNALING_PATHWAY | 1.89 | 4.47E-02 |
| WP_CILIOPATHIES | 1.89 | 4.58E-02 |
| REACTOME_SIGNALING_BY_FGFR1 | 1.89 | 4.59E-02 |
| BIOCARTA_SET_PATHWAY | 1.89 | 4.59E-02 |
| WP_NUCLEOTIDE_EXCISION_REPAIR | 1.89 | 4.64E-02 |
| WP_DNA_REPAIR_PATHWAYS_FULL_NETWORK | 1.89 | 4.64E-02 |
| WP_MELANOMA | 1.89 | 4.64E-02 |
| REACTOME_GENE_SILENCING_BY_RNA | 1.88 | 4.66E-02 |
| BIOCARTA_ECM_PATHWAY | 1.88 | 4.67E-02 |
| WP_TARGET_OF_RAPAMYCIN_SIGNALING | 1.88 | 4.68E-02 |
| KEGG_LEUKOCYTE_TRANSENDOTHELIAL_MIGRATION | 1.88 | 4.79E-02 |
| REACTOME_SUMO_IS_TRANSFERRED_FROM_E1_TO_E2_UBE2I_UBC9 | 1.87 | 4.88E-02 |
| WP_SPINAL_CORD_INJURY | 1.87 | 4.89E-02 |
| KEGG_ECM_RECEPTOR_INTERACTION | 1.87 | 4.91E-02 |
| BIOCARTA_AGPCR_PATHWAY | 1.87 | 4.93E-02 |
| REACTOME_DUAL_INCISION_IN_GG_NER | 1.87 | 4.94E-02 |
| REACTOME_GLYCOGEN_METABOLISM | 1.87 | 4.94E-02 |
| REACTOME_NEGATIVE_REGULATION_OF_MET_ACTIVITY | 1.87 | 4.94E-02 |
| REACTOME_INSULIN_RECEPTOR_SIGNALLING_CASCADE | 1.87 | 4.96E-02 |
| KEGG_CHEMOKINE_SIGNALING_PATHWAY | 1.87 | 4.99E-02 |
| REACTOME_TRANSLESION_SYNTHESIS_BY_POLK | 1.87 | 5.00E-02 |

**Supplementary Table S3.** List of concordant differentially altered gene sets in *Gpx4 KO* mouse hearts and human cardiomyopathy (FDR < 0.05).

| **Gene Set Name** | ***Gpx4* KO** | | **Human Cardiomyopathy** | |
| --- | --- | --- | --- | --- |
|  | **Normalized Enrichment Score (NES)** | **FDR** | **Normalized Enrichment Score (NES)** | **FDR** |
| HALLMARK_MYC_TARGETS_V1 | 3.21 | 0 | 5.15 | 0 |
| HALLMARK_PROTEIN_SECRETION | 2.91 | 0 | 4.23 | 0 |
| [REACTOME_SIGNALING_BY_RECEPTOR_TYROSINE_KINASES](http://www.gsea-msigdb.org/gsea/msigdb/human/geneset/REACTOME_SIGNALING_BY_RECEPTOR_TYROSINE_KINASES) | 5.33 | 0 | 4.04 | 0 |
| [HALLMARK_MTORC1_SIGNALING](http://www.gsea-msigdb.org/gsea/msigdb/human/geneset/HALLMARK_MTORC1_SIGNALING) | 3.35 | 0 | 3.53 | 0 |
| REACTOME_INTERFERON_SIGNALING | 3.04 | 0 | 3.41 | 0 |
| [HALLMARK_INTERFERON_GAMMA_RESPONSE](http://www.gsea-msigdb.org/gsea/msigdb/human/geneset/HALLMARK_INTERFERON_GAMMA_RESPONSE) | 3.7 | 0 | 3.35 | 0 |
| [REACTOME_SARS_COV_INFECTIONS](http://www.gsea-msigdb.org/gsea/msigdb/human/geneset/REACTOME_SARS_COV_INFECTIONS) | 3.48 | 0 | 3.29 | 0 |
| [REACTOME_CELL_CYCLE_MITOTIC](http://www.gsea-msigdb.org/gsea/msigdb/human/geneset/REACTOME_CELL_CYCLE_MITOTIC) | 4.18 | 0 | 3.27 | 0 |
| [HALLMARK_ANDROGEN_RESPONSE](http://www.gsea-msigdb.org/gsea/msigdb/human/geneset/HALLMARK_ANDROGEN_RESPONSE) | 3.82 | 0 | 3.26 | 0 |
| REACTOME_INTRA_GOLGI_AND_RETROGRADE_GOLGI_TO_ER_TRAFFIC | 3.17 | 0 | 3.26 | 0 |
| REACTOME_HIV_INFECTION | 2.93 | 0 | 3.26 | 0 |
| [HALLMARK_G2M_CHECKPOINT](http://www.gsea-msigdb.org/gsea/msigdb/human/geneset/HALLMARK_G2M_CHECKPOINT) | 3.83 | 0 | 3.11 | 0 |
| REACTOME_SARS_COV_2_INFECTION | 2.79 | 0 | 3.08 | 0 |
| REACTOME_TCR_SIGNALING | 2.72 | 0 | 3.05 | 0 |
| [HALLMARK_E2F_TARGETS](http://www.gsea-msigdb.org/gsea/msigdb/human/geneset/HALLMARK_E2F_TARGETS) | 3.9 | 0 | 3.04 | 0 |
| WP_TGFBETA_SIGNALING_PATHWAY | 2.84 | 0 | 3 | 0 |
| [HALLMARK_ALLOGRAFT_REJECTION](http://www.gsea-msigdb.org/gsea/msigdb/human/geneset/HALLMARK_ALLOGRAFT_REJECTION) | 5.14 | 0 | 2.98 | 0 |
| [HALLMARK_KRAS_SIGNALING_UP](http://www.gsea-msigdb.org/gsea/msigdb/human/geneset/HALLMARK_KRAS_SIGNALING_UP) | 4.13 | 0 | 2.94 | 0 |
| [HALLMARK_IL2_STAT5_SIGNALING](http://www.gsea-msigdb.org/gsea/msigdb/human/geneset/HALLMARK_IL2_STAT5_SIGNALING) | 4.48 | 0 | 2.92 | 0 |
| [REACTOME_CELL_CYCLE_CHECKPOINTS](http://www.gsea-msigdb.org/gsea/msigdb/human/geneset/REACTOME_CELL_CYCLE_CHECKPOINTS) | 3.61 | 0 | 2.9 | 0 |
| [HALLMARK_TNFA_SIGNALING_VIA_NFKB](http://www.gsea-msigdb.org/gsea/msigdb/human/geneset/HALLMARK_TNFA_SIGNALING_VIA_NFKB) | 6.63 | 0 | 2.88 | 0 |
| REACTOME_GOLGI_TO_ER_RETROGRADE_TRANSPORT | 3.09 | 0 | 2.77 | 0 |
| [PID_PDGFRB_PATHWAY](http://www.gsea-msigdb.org/gsea/msigdb/human/geneset/PID_PDGFRB_PATHWAY) | 3.56 | 0 | 2.76 | 0.001 |
| [KEGG_REGULATION_OF_ACTIN_CYTOSKELETON](http://www.gsea-msigdb.org/gsea/msigdb/human/geneset/KEGG_REGULATION_OF_ACTIN_CYTOSKELETON) | 4.82 | 0 | 2.73 | 0.001 |
| REACTOME_S_PHASE | 2.89 | 0 | 2.71 | 0.001 |
| NABA_ECM_GLYCOPROTEINS | 2.86 | 0 | 2.69 | 0.001 |
| [NABA_CORE_MATRISOME](http://www.gsea-msigdb.org/gsea/msigdb/human/geneset/NABA_CORE_MATRISOME) | 3.91 | 0 | 2.68 | 0.001 |
| [REACTOME_MITOTIC_G1_PHASE_AND_G1_S_TRANSITION](http://www.gsea-msigdb.org/gsea/msigdb/human/geneset/REACTOME_MITOTIC_G1_PHASE_AND_G1_S_TRANSITION) | 3.65 | 0 | 2.63 | 0.001 |
| KEGG_NATURAL_KILLER_CELL_MEDIATED_CYTOTOXICITY | 2.79 | 0 | 2.61 | 0.001 |
| [REACTOME_NEUTROPHIL_DEGRANULATION](http://www.gsea-msigdb.org/gsea/msigdb/human/geneset/REACTOME_NEUTROPHIL_DEGRANULATION) | 6.12 | 0 | 2.6 | 0.001 |
| KEGG_PATHWAYS_IN_CANCER | 3.07 | 0 | 2.6 | 0.001 |
| [KEGG_CELL_ADHESION_MOLECULES_CAMS](http://www.gsea-msigdb.org/gsea/msigdb/human/geneset/KEGG_CELL_ADHESION_MOLECULES_CAMS) | 3.49 | 0 | 2.59 | 0.001 |
| [WP_MEASLES_VIRUS_INFECTION](http://www.gsea-msigdb.org/gsea/msigdb/human/geneset/WP_MEASLES_VIRUS_INFECTION) | 3.39 | 0 | 2.59 | 0.001 |
| [PID_AMB2_NEUTROPHILS_PATHWAY](http://www.gsea-msigdb.org/gsea/msigdb/human/geneset/PID_AMB2_NEUTROPHILS_PATHWAY) | 3.56 | 0 | 2.58 | 0.001 |
| [REACTOME_PLATELET_ACTIVATION_SIGNALING_AND_AGGREGATION](http://www.gsea-msigdb.org/gsea/msigdb/human/geneset/REACTOME_PLATELET_ACTIVATION_SIGNALING_AND_AGGREGATION) | 3.78 | 0 | 2.52 | 0.002 |
| WP_PI3KAKT_SIGNALING_PATHWAY | 3.1 | 0 | 2.52 | 0.002 |
| [REACTOME_CELL_SURFACE_INTERACTIONS_AT_THE_VASCULAR_WALL](http://www.gsea-msigdb.org/gsea/msigdb/human/geneset/REACTOME_CELL_SURFACE_INTERACTIONS_AT_THE_VASCULAR_WALL) | 4.31 | 0 | 2.48 | 0.003 |
| [WP_OVERVIEW_OF_PROINFLAMMATORY_AND_PROFIBROTIC_MEDIATORS](http://www.gsea-msigdb.org/gsea/msigdb/human/geneset/WP_OVERVIEW_OF_PROINFLAMMATORY_AND_PROFIBROTIC_MEDIATORS) | 3.3 | 0 | 2.48 | 0.003 |
| [REACTOME_SIGNALING_BY_INTERLEUKINS](http://www.gsea-msigdb.org/gsea/msigdb/human/geneset/REACTOME_SIGNALING_BY_INTERLEUKINS) | 5.22 | 0 | 2.47 | 0.003 |
| REACTOME_M_PHASE | 2.99 | 0 | 2.47 | 0.003 |
| [HALLMARK_COMPLEMENT](http://www.gsea-msigdb.org/gsea/msigdb/human/geneset/HALLMARK_COMPLEMENT) | 3.89 | 0 | 2.45 | 0.003 |
| [REACTOME_IMMUNOREGULATORY_INTERACTIONS_BETWEEN_A_LYMPHOID_AND_A_NON_LYMPHOID_CELL](http://www.gsea-msigdb.org/gsea/msigdb/human/geneset/REACTOME_IMMUNOREGULATORY_INTERACTIONS_BETWEEN_A_LYMPHOID_AND_A_NON_LYMPHOID_CELL) | 4.4 | 0 | 2.42 | 0.004 |
| WP_MALIGNANT_PLEURAL_MESOTHELIOMA | 3.23 | 0 | 2.4 | 0.004 |
| [WP_NETWORK_MAP_OF_SARSCOV2_SIGNALING_PATHWAY](http://www.gsea-msigdb.org/gsea/msigdb/human/geneset/WP_NETWORK_MAP_OF_SARSCOV2_SIGNALING_PATHWAY) | 3.71 | 0 | 2.4 | 0.005 |
| [REACTOME_MITOTIC_METAPHASE_AND_ANAPHASE](http://www.gsea-msigdb.org/gsea/msigdb/human/geneset/REACTOME_MITOTIC_METAPHASE_AND_ANAPHASE) | 3.38 | 0 | 2.38 | 0.005 |
| REACTOME_MAPK_FAMILY_SIGNALING_CASCADES | 2.8 | 0 | 2.37 | 0.006 |
| [HALLMARK_EPITHELIAL_MESENCHYMAL_TRANSITION](http://www.gsea-msigdb.org/gsea/msigdb/human/geneset/HALLMARK_EPITHELIAL_MESENCHYMAL_TRANSITION) | 6.98 | 0 | 2.36 | 0.006 |
| [HALLMARK_APOPTOSIS](http://www.gsea-msigdb.org/gsea/msigdb/human/geneset/HALLMARK_APOPTOSIS) | 3.32 | 0 | 2.36 | 0.006 |
| REACTOME_G2_M_CHECKPOINTS | 2.9 | 0 | 2.34 | 0.006 |
| [WP_EBOLA_VIRUS_INFECTION_IN_HOST](http://www.gsea-msigdb.org/gsea/msigdb/human/geneset/WP_EBOLA_VIRUS_INFECTION_IN_HOST) | 3.44 | 0 | 2.33 | 0.006 |
| REACTOME_MHC_CLASS_II_ANTIGEN_PRESENTATION | 3.22 | 0 | 2.32 | 0.007 |
| KEGG_T_CELL_RECEPTOR_SIGNALING_PATHWAY | 2.83 | 0 | 2.3 | 0.008 |
| REACTOME_PROGRAMMED_CELL_DEATH | 2.69 | 0 | 2.3 | 0.008 |
| REACTOME_CLEC7A_DECTIN_1_SIGNALING | 2.89 | 0 | 2.27 | 0.009 |
| [WP_RETINOBLASTOMA_GENE_IN_CANCER](http://www.gsea-msigdb.org/gsea/msigdb/human/geneset/WP_RETINOBLASTOMA_GENE_IN_CANCER) | 3.66 | 0 | 2.26 | 0.009 |
| REACTOME_INTERFERON_ALPHA_BETA_SIGNALING | 2.77 | 0 | 2.25 | 0.01 |
| BIOCARTA_LYMPHOCYTE_PATHWAY | 2.63 | 0 | 2.24 | 0.01 |
| REACTOME_SARS_COV_1_INFECTION | 2.61 | 0 | 2.24 | 0.01 |
| REACTOME_MITOTIC_SPINDLE_CHECKPOINT | 2.66 | 0 | 2.23 | 0.011 |
| WP_G1_TO_S_CELL_CYCLE_CONTROL | 3.21 | 0 | 2.22 | 0.011 |
| REACTOME_HOST_INTERACTIONS_OF_HIV_FACTORS | 3.16 | 0 | 2.22 | 0.011 |
| KEGG_CELL_CYCLE | 3.11 | 0 | 2.22 | 0.012 |
| [REACTOME_PARASITE_INFECTION](http://www.gsea-msigdb.org/gsea/msigdb/human/geneset/REACTOME_PARASITE_INFECTION) | 3.44 | 0 | 2.21 | 0.012 |
| REACTOME_SIGNALING_BY_MET | 3.11 | 0 | 2.21 | 0.012 |
| REACTOME_RESOLUTION_OF_SISTER_CHROMATID_COHESION | 3.27 | 0 | 2.2 | 0.012 |
| WP_HEPATITIS_B_INFECTION | 3.17 | 0 | 2.2 | 0.012 |
| REACTOME_TOLL_LIKE_RECEPTOR_CASCADES | 2.71 | 0 | 2.2 | 0.012 |
| WP_CELL_CYCLE | 3.11 | 0 | 2.18 | 0.014 |
| PID_ATF2_PATHWAY | 2.85 | 0 | 2.18 | 0.014 |
| [REACTOME_ASPARAGINE_N_LINKED_GLYCOSYLATION](http://www.gsea-msigdb.org/gsea/msigdb/human/geneset/REACTOME_ASPARAGINE_N_LINKED_GLYCOSYLATION) | 3.32 | 0 | 2.17 | 0.014 |
| [REACTOME_RHO_GTPASE_EFFECTORS](http://www.gsea-msigdb.org/gsea/msigdb/human/geneset/REACTOME_RHO_GTPASE_EFFECTORS) | 4.9 | 0 | 2.15 | 0.016 |
| [WP_REGULATION_OF_ACTIN_CYTOSKELETON](http://www.gsea-msigdb.org/gsea/msigdb/human/geneset/WP_REGULATION_OF_ACTIN_CYTOSKELETON) | 3.47 | 0 | 2.11 | 0.019 |
| BIOCARTA_MONOCYTE_PATHWAY | 2.98 | 0 | 2.1 | 0.02 |
| [REACTOME_RAC2_GTPASE_CYCLE](http://www.gsea-msigdb.org/gsea/msigdb/human/geneset/REACTOME_RAC2_GTPASE_CYCLE) | 3.51 | 0 | 2.07 | 0.023 |
| REACTOME_SEPARATION_OF_SISTER_CHROMATIDS | 3.19 | 0 | 2.07 | 0.023 |
| PID_AURORA_B_PATHWAY | 3.13 | 0 | 2.06 | 0.024 |
| HALLMARK_CHOLESTEROL_HOMEOSTASIS | 2.95 | 0 | 2.06 | 0.024 |
| REACTOME_CHEMOKINE_RECEPTORS_BIND_CHEMOKINES | 3.17 | 0 | 2.01 | 0.03 |
| [KEGG_TOLL_LIKE_RECEPTOR_SIGNALING_PATHWAY](http://www.gsea-msigdb.org/gsea/msigdb/human/geneset/KEGG_TOLL_LIKE_RECEPTOR_SIGNALING_PATHWAY) | 3.56 | 0 | 2 | 0.03 |
| KEGG_EPITHELIAL_CELL_SIGNALING_IN_HELICOBACTER_PYLORI_INFECTION | 2.71 | 0 | 2 | 0.03 |
| [REACTOME_FCGAMMA_RECEPTOR_FCGR_DEPENDENT_PHAGOCYTOSIS](http://www.gsea-msigdb.org/gsea/msigdb/human/geneset/REACTOME_FCGAMMA_RECEPTOR_FCGR_DEPENDENT_PHAGOCYTOSIS) | 3.4 | 0 | 1.99 | 0.031 |
| REACTOME_CLATHRIN_MEDIATED_ENDOCYTOSIS | 2.67 | 0 | 1.99 | 0.032 |
| [REACTOME_RHO_GTPASE_CYCLE](http://www.gsea-msigdb.org/gsea/msigdb/human/geneset/REACTOME_RHO_GTPASE_CYCLE) | 3.49 | 0 | 1.97 | 0.034 |
| REACTOME_SIGNALING_BY_VEGF | 2.65 | 0 | 1.97 | 0.034 |
| REACTOME_DNA_REPLICATION | 2.65 | 0 | 1.96 | 0.036 |
| [PID_AVB3_INTEGRIN_PATHWAY](http://www.gsea-msigdb.org/gsea/msigdb/human/geneset/PID_AVB3_INTEGRIN_PATHWAY) | 3.61 | 0 | 1.94 | 0.039 |
| [WP_TOLLLIKE_RECEPTOR_SIGNALING_PATHWAY](http://www.gsea-msigdb.org/gsea/msigdb/human/geneset/WP_TOLLLIKE_RECEPTOR_SIGNALING_PATHWAY) | 3.57 | 0 | 1.94 | 0.039 |
| [PID_CXCR4_PATHWAY](http://www.gsea-msigdb.org/gsea/msigdb/human/geneset/PID_CXCR4_PATHWAY) | 4.06 | 0 | 1.93 | 0.04 |
| REACTOME_DOWNSTREAM_SIGNALING_EVENTS_OF_B_CELL_RECEPTOR_BCR | 2.69 | 0 | 1.92 | 0.042 |
| [KEGG_FOCAL_ADHESION](http://www.gsea-msigdb.org/gsea/msigdb/human/geneset/KEGG_FOCAL_ADHESION) | 4.43 | 0 | 1.91 | 0.043 |
| WP_PANCREATIC_ADENOCARCINOMA_PATHWAY | 2.85 | 0 | 1.91 | 0.043 |
| WP_TGFBETA_RECEPTOR_SIGNALING_IN_SKELETAL_DYSPLASIAS | 2.63 | 0 | 1.91 | 0.043 |
| BIOCARTA_CELLCYCLE_PATHWAY | 3.04 | 0 | 1.9 | 0.044 |
| REACTOME_SIGNALING_BY_GPCR | 2.69 | 0 | 1.9 | 0.044 |
| REACTOME_SPHINGOLIPID_METABOLISM | 2.65 | 0 | 1.9 | 0.044 |
| PID_INTEGRIN_A4B1_PATHWAY | 3.27 | 0 | 1.89 | 0.045 |
| [KEGG_LEUKOCYTE_TRANSENDOTHELIAL_MIGRATION](http://www.gsea-msigdb.org/gsea/msigdb/human/geneset/KEGG_LEUKOCYTE_TRANSENDOTHELIAL_MIGRATION) | 4.13 | 0 | 1.88 | 0.048 |
| [KEGG_ECM_RECEPTOR_INTERACTION](http://www.gsea-msigdb.org/gsea/msigdb/human/geneset/KEGG_ECM_RECEPTOR_INTERACTION) | 3.39 | 0 | 1.87 | 0.049 |
| [WP_SPINAL_CORD_INJURY](http://www.gsea-msigdb.org/gsea/msigdb/human/geneset/WP_SPINAL_CORD_INJURY) | 3.37 | 0 | 1.87 | 0.049 |
| HALLMARK_UV_RESPONSE_DN | 2.47 | 0.001 | 3.6 | 0 |
| REACTOME_FC_EPSILON_RECEPTOR_FCERI_SIGNALING | 2.51 | 0.001 | 3.34 | 0 |
| PID_ERBB1_DOWNSTREAM_PATHWAY | 2.6 | 0.001 | 3.15 | 0 |
| REACTOME_SIGNALING_BY_THE_B_CELL_RECEPTOR_BCR | 2.57 | 0.001 | 2.63 | 0.001 |
| WP_TYPE_II_INTERFERON_SIGNALING | 2.61 | 0.001 | 2.36 | 0.006 |
| REACTOME_MITOTIC_PROMETAPHASE | 2.59 | 0.001 | 2.21 | 0.012 |
| PID_INTEGRIN5_PATHWAY | 2.55 | 0.001 | 2.2 | 0.012 |
| REACTOME_SARS_COV_2_HOST_INTERACTIONS | 2.53 | 0.001 | 2.16 | 0.015 |
| WP_SARSCOV2_INNATE_IMMUNITY_EVASION_AND_CELLSPECIFIC_IMMUNE_RESPONSE | 2.52 | 0.001 | 2.05 | 0.025 |
| WP_TGFBETA_RECEPTOR_SIGNALING | 2.5 | 0.001 | 1.99 | 0.032 |
| WP_APOPTOSISRELATED_NETWORK_DUE_TO_ALTERED_NOTCH3_IN_OVARIAN_CANCER | 2.46 | 0.001 | 1.94 | 0.039 |
| PID_NFAT_TFPATHWAY | 2.56 | 0.001 | 1.93 | 0.039 |
| KEGG_B_CELL_RECEPTOR_SIGNALING_PATHWAY | 2.46 | 0.001 | 1.91 | 0.042 |
| WP_SPHINGOLIPID_METABOLISM_IN_SENESCENCE | 2.46 | 0.001 | 1.9 | 0.044 |
| REACTOME_DISEASES_OF_SIGNAL_TRANSDUCTION_BY_GROWTH_FACTOR_RECEPTORS_AND_SECOND_MESSENGERS | 2.4 | 0.002 | 3.7 | 0 |
| REACTOME_ANTIVIRAL_MECHANISM_BY_IFN_STIMULATED_GENES | 2.39 | 0.002 | 3.21 | 0 |
| WP_EGFEGFR_SIGNALING_PATHWAY | 2.4 | 0.002 | 2.62 | 0.001 |
| WP_FOCAL_ADHESION_PI3KAKTMTORSIGNALING_PATHWAY | 2.41 | 0.002 | 2.53 | 0.002 |
| WP_TCELL_RECEPTOR_SIGNALING_PATHWAY | 2.43 | 0.002 | 2.42 | 0.004 |
| REACTOME_REGULATION_OF_MRNA_STABILITY_BY_PROTEINS_THAT_BIND_AU_RICH_ELEMENTS | 2.42 | 0.002 | 2.12 | 0.019 |
| PID_CD8_TCR_PATHWAY | 2.38 | 0.002 | 2.01 | 0.029 |
| REACTOME_SIGNALING_BY_TGFB_FAMILY_MEMBERS | 2.33 | 0.003 | 2.79 | 0 |
| REACTOME_SIGNALING_BY_TGF_BETA_RECEPTOR_COMPLEX | 2.34 | 0.003 | 2.76 | 0.001 |
| WP_GLIOBLASTOMA_SIGNALING_PATHWAYS | 2.35 | 0.003 | 2.38 | 0.005 |
| REACTOME_APOPTOSIS | 2.36 | 0.003 | 2.21 | 0.012 |
| PID_IL12_2PATHWAY | 2.34 | 0.003 | 2.16 | 0.015 |
| WP_B_CELL_RECEPTOR_SIGNALING_PATHWAY | 2.33 | 0.003 | 2 | 0.03 |
| REACTOME_CYCLIN_A_CDK2_ASSOCIATED_EVENTS_AT_S_PHASE_ENTRY | 2.34 | 0.003 | 1.99 | 0.032 |
| REACTOME_DNA_REPLICATION_PRE_INITIATION | 2.37 | 0.003 | 1.9 | 0.044 |
| REACTOME_FCERI_MEDIATED_NF_KB_ACTIVATION | 2.31 | 0.004 | 2.61 | 0.001 |
| HALLMARK_GLYCOLYSIS | 2.28 | 0.004 | 2.22 | 0.011 |
| REACTOME_METABOLISM_OF_CARBOHYDRATES | 2.29 | 0.004 | 2.12 | 0.019 |
| REACTOME_SARS_COV_2_ACTIVATES_MODULATES_INNATE_AND_ADAPTIVE_IMMUNE_RESPONSES | 2.3 | 0.004 | 1.95 | 0.037 |
| PID_ERBB1_INTERNALIZATION_PATHWAY | 2.28 | 0.004 | 1.94 | 0.039 |
| REACTOME_TRANSPORT_TO_THE_GOLGI_AND_SUBSEQUENT_MODIFICATION | 2.28 | 0.005 | 2.2 | 0.013 |
| REACTOME_ONCOGENIC_MAPK_SIGNALING | 2.27 | 0.005 | 2 | 0.03 |
| REACTOME_TRANS_GOLGI_NETWORK_VESICLE_BUDDING | 2.24 | 0.006 | 2.48 | 0.003 |
| REACTOME_GENERATION_OF_SECOND_MESSENGER_MOLECULES | 2.24 | 0.006 | 2.11 | 0.019 |
| WP_PHYSICOCHEMICAL_FEATURES_AND_TOXICITYASSOCIATED_PATHWAYS | 2.21 | 0.007 | 2.44 | 0.004 |
| WP_TCELL_ANTIGEN_RECEPTOR_TCR_PATHWAY_DURING_STAPHYLOCOCCUS_AUREUS_INFECTION | 2.21 | 0.007 | 2.17 | 0.014 |
| REACTOME_BINDING_AND_UPTAKE_OF_LIGANDS_BY_SCAVENGER_RECEPTORS | 2.19 | 0.008 | 2.84 | 0 |
| REACTOME_DEUBIQUITINATION | 2.17 | 0.008 | 2.7 | 0.001 |
| BIOCARTA_CCR5_PATHWAY | 2.17 | 0.008 | 2.09 | 0.021 |
| REACTOME_POTENTIAL_THERAPEUTICS_FOR_SARS | 2.17 | 0.008 | 1.97 | 0.034 |
| PID_ARF6_TRAFFICKING_PATHWAY | 2.17 | 0.008 | 1.92 | 0.042 |
| REACTOME_UB_SPECIFIC_PROCESSING_PROTEASES | 2.16 | 0.009 | 2.48 | 0.003 |
| HALLMARK_UNFOLDED_PROTEIN_RESPONSE | 2.16 | 0.009 | 2.3 | 0.008 |
| PID_FAK_PATHWAY | 2.16 | 0.009 | 1.92 | 0.04 |
| REACTOME_EXTRA_NUCLEAR_ESTROGEN_SIGNALING | 2.14 | 0.01 | 2.65 | 0.001 |
| REACTOME_HIV_LIFE_CYCLE | 2.14 | 0.01 | 2.62 | 0.001 |
| REACTOME_SIGNALING_BY_WNT | 2.13 | 0.01 | 2.56 | 0.002 |
| WP_ALLOGRAFT_REJECTION | 2.13 | 0.01 | 2.1 | 0.021 |
| REACTOME_SIGNALING_BY_HIPPO | 2.14 | 0.01 | 2.07 | 0.023 |
| KEGG_PANCREATIC_CANCER | 2.11 | 0.011 | 2 | 0.03 |
| KEGG_ANTIGEN_PROCESSING_AND_PRESENTATION | 2.09 | 0.012 | 3.03 | 0 |
| PID_FGF_PATHWAY | 2.11 | 0.012 | 2.03 | 0.028 |
| REACTOME_TRANSLOCATION_OF_SLC2A4_GLUT4_TO_THE_PLASMA_MEMBRANE | 2.11 | 0.012 | 1.94 | 0.039 |
| WP_ACUTE_VIRAL_MYOCARDITIS | 2.09 | 0.013 | 1.92 | 0.042 |
| REACTOME_G_ALPHA_I_SIGNALLING_EVENTS | 2.07 | 0.014 | 2.52 | 0.002 |
| REACTOME_TGF_BETA_RECEPTOR_SIGNALING_ACTIVATES_SMADS | 2.07 | 0.014 | 2.44 | 0.003 |
| KEGG_NEUROTROPHIN_SIGNALING_PATHWAY | 2.06 | 0.014 | 2.17 | 0.015 |
| WP_PARKINUBIQUITIN_PROTEASOMAL_SYSTEM_PATHWAY | 2.06 | 0.015 | 2.05 | 0.025 |
| KEGG_VIRAL_MYOCARDITIS | 2.04 | 0.016 | 2.34 | 0.006 |
| WP_EPITHELIAL_TO_MESENCHYMAL_TRANSITION_IN_COLORECTAL_CANCER | 2.02 | 0.017 | 2.5 | 0.003 |
| WP_CHROMOSOMAL_AND_MICROSATELLITE_INSTABILITY_IN_COLORECTAL_CANCER | 2.01 | 0.018 | 2.49 | 0.003 |
| REACTOME_HCMV_INFECTION | 2.01 | 0.018 | 2 | 0.03 |
| REACTOME_CONSTITUTIVE_SIGNALING_BY_ABERRANT_PI3K_IN_CANCER | 2.01 | 0.018 | 1.94 | 0.038 |
| REACTOME_ROLE_OF_LAT2_NTAL_LAB_ON_CALCIUM_MOBILIZATION | 2 | 0.019 | 2.94 | 0 |
| KEGG_TGF_BETA_SIGNALING_PATHWAY | 2 | 0.019 | 2.85 | 0 |
| REACTOME_GOLGI_ASSOCIATED_VESICLE_BIOGENESIS | 2 | 0.019 | 2.44 | 0.004 |
| REACTOME_ANTIGEN_ACTIVATES_B_CELL_RECEPTOR_BCR_LEADING_TO_GENERATION_OF_SECOND_MESSENGERS | 1.97 | 0.022 | 2.76 | 0.001 |
| BIOCARTA_PAR1_PATHWAY | 1.97 | 0.022 | 2.32 | 0.007 |
| WP_THYROID_STIMULATING_HORMONE_TSH_SIGNALING_PATHWAY | 1.97 | 0.022 | 2.17 | 0.014 |
| WP_PATHOGENESIS_OF_SARSCOV2_MEDIATED_BY_NSP9NSP10_COMPLEX | 1.97 | 0.022 | 2.14 | 0.017 |
| REACTOME_ACTIVATION_OF_THE_MRNA_UPON_BINDING_OF_THE_CAP_BINDING_COMPLEX_AND_EIFS_AND_SUBSEQUENT_BINDING_TO_43S | 1.98 | 0.022 | 2.07 | 0.023 |
| KEGG_OOCYTE_MEIOSIS | 1.97 | 0.022 | 1.94 | 0.038 |
| REACTOME_COPI_MEDIATED_ANTEROGRADE_TRANSPORT | 1.97 | 0.023 | 2.11 | 0.019 |
| REACTOME_REGULATION_OF_PTEN_STABILITY_AND_ACTIVITY | 1.96 | 0.023 | 2.11 | 0.02 |
| WP_MAMMARY_GLAND_DEVELOPMENT_PATHWAY_EMBRYONIC_DEVELOPMENT_STAGE_1_OF_4 | 1.96 | 0.023 | 2.05 | 0.025 |
| REACTOME_INTRACELLULAR_SIGNALING_BY_SECOND_MESSENGERS | 1.95 | 0.024 | 2.75 | 0.001 |
| WP_GASTRIN_SIGNALING_PATHWAY | 1.95 | 0.024 | 2.43 | 0.004 |
| PID_TGFBR_PATHWAY | 1.95 | 0.024 | 2.29 | 0.008 |
| HALLMARK_PI3K_AKT_MTOR_SIGNALING | 1.95 | 0.024 | 2.06 | 0.023 |
| KEGG_SPLICEOSOME | 1.93 | 0.026 | 2.54 | 0.002 |
| BIOCARTA_BAD_PATHWAY | 1.93 | 0.026 | 2.35 | 0.006 |
| PID_BCR_5PATHWAY | 1.93 | 0.026 | 2.07 | 0.023 |
| BIOCARTA_TCR_PATHWAY | 1.93 | 0.027 | 2.02 | 0.029 |
| WP_MELANOMA | 1.93 | 0.027 | 1.89 | 0.046 |
| REACTOME_IRON_UPTAKE_AND_TRANSPORT | 1.92 | 0.028 | 2.13 | 0.017 |
| KEGG_COLORECTAL_CANCER | 1.91 | 0.029 | 2.52 | 0.002 |
| BIOCARTA_PRION_PATHWAY | 1.9 | 0.029 | 2.5 | 0.002 |
| REACTOME_ER_TO_GOLGI_ANTEROGRADE_TRANSPORT | 1.89 | 0.031 | 1.98 | 0.033 |
| REACTOME_SIGNALING_BY_ALK_IN_CANCER | 1.89 | 0.032 | 2.17 | 0.014 |
| REACTOME_NEGATIVE_REGULATION_OF_THE_PI3K_AKT_NETWORK | 1.88 | 0.033 | 2.55 | 0.002 |
| BIOCARTA_GSK3_PATHWAY | 1.88 | 0.033 | 2.47 | 0.003 |
| BIOCARTA_TCYTOTOXIC_PATHWAY | 1.87 | 0.035 | 2.5 | 0.002 |
| PID_ATR_PATHWAY | 1.86 | 0.036 | 1.94 | 0.039 |
| PID_CD8_TCR_DOWNSTREAM_PATHWAY | 1.85 | 0.037 | 2.85 | 0 |
| REACTOME_CHK1_CHK2_CDS1_MEDIATED_INACTIVATION_OF_CYCLIN_B_CDK1_COMPLEX | 1.85 | 0.037 | 2.1 | 0.02 |
| REACTOME_E2F_MEDIATED_REGULATION_OF_DNA_REPLICATION | 1.85 | 0.038 | 2.39 | 0.005 |
| BIOCARTA_THELPER_PATHWAY | 1.84 | 0.038 | 2.35 | 0.006 |
| REACTOME_INHIBITION_OF_REPLICATION_INITIATION_OF_DAMAGED_DNA_BY_RB1_E2F1 | 1.84 | 0.038 | 1.97 | 0.034 |
| REACTOME_FCERI_MEDIATED_CA_2_MOBILIZATION | 1.84 | 0.039 | 2.66 | 0.001 |
| BIOCARTA_GCR_PATHWAY | 1.83 | 0.041 | 2.47 | 0.003 |
| REACTOME_NEGATIVE_REGULATION_OF_MET_ACTIVITY | 1.82 | 0.042 | 1.87 | 0.049 |
| WP_HEAD_AND_NECK_SQUAMOUS_CELL_CARCINOMA | 1.81 | 0.044 | 1.99 | 0.032 |
| BIOCARTA_TEL_PATHWAY | 1.81 | 0.045 | 2.39 | 0.005 |
| WP_WNT_SIGNALING | 1.8 | 0.046 | 2.19 | 0.013 |
| PID_S1P_S1P2_PATHWAY | 1.8 | 0.046 | 2.18 | 0.014 |
| REACTOME_PTEN_REGULATION | 1.79 | 0.047 | 2.01 | 0.03 |
| KEGG_WNT_SIGNALING_PATHWAY | 1.79 | 0.048 | 2.72 | 0.001 |
| REACTOME_SPHINGOLIPID_DE_NOVO_BIOSYNTHESIS | 1.79 | 0.048 | 2.09 | 0.021 |

**Supplementary Table S4.** List of discordant differentially altered gene sets in *Gpx4 KO* mouse hearts and human cardiomyopathy (FDR < 0.05).

| **Gene Set Name** | ***Gpx4* KO** | | **Human Cardiomyopathy** | |
| --- | --- | --- | --- | --- |
|  | **Normalized Enrichment Score (NES)** | **FDR** | **Normalized Enrichment Score (NES)** | **FDR** |
| [HALLMARK_OXIDATIVE_PHOSPHORYLATION](http://www.gsea-msigdb.org/gsea/msigdb/human/geneset/HALLMARK_OXIDATIVE_PHOSPHORYLATION) | -9.42 | 0 | 4.73 | 0 |
| [REACTOME_THE_CITRIC_ACID_TCA_CYCLE_AND_RESPIRATORY_ELECTRON_TRANSPORT](http://www.gsea-msigdb.org/gsea/msigdb/human/geneset/REACTOME_THE_CITRIC_ACID_TCA_CYCLE_AND_RESPIRATORY_ELECTRON_TRANSPORT) | -9.08 | 0 | 2.8 | 0 |
| [HALLMARK_ADIPOGENESIS](http://www.gsea-msigdb.org/gsea/msigdb/human/geneset/HALLMARK_ADIPOGENESIS) | -5.41 | 0 | 2.83 | 0 |
| [REACTOME_PROTEIN_LOCALIZATION](http://www.gsea-msigdb.org/gsea/msigdb/human/geneset/REACTOME_PROTEIN_LOCALIZATION) | -5.25 | 0 | 3.98 | 0 |
| [HALLMARK_FATTY_ACID_METABOLISM](http://www.gsea-msigdb.org/gsea/msigdb/human/geneset/HALLMARK_FATTY_ACID_METABOLISM) | -5.16 | 0 | 3.4 | 0 |
| [REACTOME_PYRUVATE_METABOLISM_AND_CITRIC_ACID_TCA_CYCLE](http://www.gsea-msigdb.org/gsea/msigdb/human/geneset/REACTOME_PYRUVATE_METABOLISM_AND_CITRIC_ACID_TCA_CYCLE) | -5.07 | 0 | 3.54 | 0 |
| [REACTOME_CITRIC_ACID_CYCLE_TCA_CYCLE](http://www.gsea-msigdb.org/gsea/msigdb/human/geneset/REACTOME_CITRIC_ACID_CYCLE_TCA_CYCLE) | -4.72 | 0 | 3.17 | 0 |
| [KEGG_VALINE_LEUCINE_AND_ISOLEUCINE_DEGRADATION](http://www.gsea-msigdb.org/gsea/msigdb/human/geneset/KEGG_VALINE_LEUCINE_AND_ISOLEUCINE_DEGRADATION) | -4.65 | 0 | 3.69 | 0 |
| [WP_LEUCINE_ISOLEUCINE_AND_VALINE_METABOLISM](http://www.gsea-msigdb.org/gsea/msigdb/human/geneset/WP_LEUCINE_ISOLEUCINE_AND_VALINE_METABOLISM) | -4.35 | 0 | 3.23 | 0 |
| [WP_TCA_CYCLE_AKA_KREBS_OR_CITRIC_ACID_CYCLE](http://www.gsea-msigdb.org/gsea/msigdb/human/geneset/WP_TCA_CYCLE_AKA_KREBS_OR_CITRIC_ACID_CYCLE) | -4.28 | 0 | 3.15 | 0 |
| [REACTOME_BRANCHED_CHAIN_AMINO_ACID_CATABOLISM](http://www.gsea-msigdb.org/gsea/msigdb/human/geneset/REACTOME_BRANCHED_CHAIN_AMINO_ACID_CATABOLISM) | -4 | 0 | 3.39 | 0 |
| [KEGG_CITRATE_CYCLE_TCA_CYCLE](http://www.gsea-msigdb.org/gsea/msigdb/human/geneset/KEGG_CITRATE_CYCLE_TCA_CYCLE) | -3.98 | 0 | 3.43 | 0 |
| [REACTOME_PEROXISOMAL_PROTEIN_IMPORT](http://www.gsea-msigdb.org/gsea/msigdb/human/geneset/REACTOME_PEROXISOMAL_PROTEIN_IMPORT) | -3.17 | 0 | 3.28 | 0 |
| [WP_AMINO_ACID_METABOLISM](http://www.gsea-msigdb.org/gsea/msigdb/human/geneset/WP_AMINO_ACID_METABOLISM) | -3.16 | 0 | 3.07 | 0 |
| [REACTOME_TRANSLATION](http://www.gsea-msigdb.org/gsea/msigdb/human/geneset/REACTOME_TRANSLATION) | -2.78 | 0 | 3.42 | 0 |
| [REACTOME_PROCESSING_OF_CAPPED_INTRON_CONTAINING_PRE_MRNA](http://www.gsea-msigdb.org/gsea/msigdb/human/geneset/REACTOME_PROCESSING_OF_CAPPED_INTRON_CONTAINING_PRE_MRNA) | -2.34 | 0.004 | 3.38 | 0 |
| REACTOME_MRNA_SPLICING | -1.98 | 0.035 | 3.3 | 0 |
| [KEGG_PROPANOATE_METABOLISM](http://www.gsea-msigdb.org/gsea/msigdb/human/geneset/KEGG_PROPANOATE_METABOLISM) | -3.7 | 0 | 2.73 | 0.001 |
| [KEGG_BUTANOATE_METABOLISM](http://www.gsea-msigdb.org/gsea/msigdb/human/geneset/KEGG_BUTANOATE_METABOLISM) | -2.81 | 0 | 2.72 | 0.001 |
| [KEGG_BETA_ALANINE_METABOLISM](http://www.gsea-msigdb.org/gsea/msigdb/human/geneset/KEGG_BETA_ALANINE_METABOLISM) | -2.61 | 0.001 | 2.71 | 0.001 |
| [WP_KREBS_CYCLE_DISORDERS](http://www.gsea-msigdb.org/gsea/msigdb/human/geneset/WP_KREBS_CYCLE_DISORDERS) | -2.46 | 0.002 | 2.73 | 0.001 |
| [WP_TCA_CYCLE_AND_DEFICIENCY_OF_PYRUVATE_DEHYDROGENASE_COMPLEX_PDHC](http://www.gsea-msigdb.org/gsea/msigdb/human/geneset/WP_TCA_CYCLE_AND_DEFICIENCY_OF_PYRUVATE_DEHYDROGENASE_COMPLEX_PDHC) | -2.44 | 0.002 | 2.76 | 0.001 |
| REACTOME_METABOLISM_OF_AMINO_ACIDS_AND_DERIVATIVES | -2.06 | 0.021 | 2.43 | 0.004 |
| [KEGG_PEROXISOME](http://www.gsea-msigdb.org/gsea/msigdb/human/geneset/KEGG_PEROXISOME) | -3.3 | 0 | 2.37 | 0.005 |
| [KEGG_PYRUVATE_METABOLISM](http://www.gsea-msigdb.org/gsea/msigdb/human/geneset/KEGG_PYRUVATE_METABOLISM) | -3.04 | 0 | 2.39 | 0.005 |
| [REACTOME_TRNA_AMINOACYLATION](http://www.gsea-msigdb.org/gsea/msigdb/human/geneset/REACTOME_TRNA_AMINOACYLATION) | -2.21 | 0.01 | 2.39 | 0.005 |
| [REACTOME_MITOCHONDRIAL_TRANSLATION](http://www.gsea-msigdb.org/gsea/msigdb/human/geneset/REACTOME_MITOCHONDRIAL_TRANSLATION) | -6.75 | 0 | 2.34 | 0.006 |
| [REACTOME_MITOCHONDRIAL_TRNA_AMINOACYLATION](http://www.gsea-msigdb.org/gsea/msigdb/human/geneset/REACTOME_MITOCHONDRIAL_TRNA_AMINOACYLATION) | -2.7 | 0 | 2.35 | 0.006 |
| HALLMARK_PEROXISOME | -2.18 | 0.012 | 2.34 | 0.006 |
| [REACTOME_PYRUVATE_METABOLISM](http://www.gsea-msigdb.org/gsea/msigdb/human/geneset/REACTOME_PYRUVATE_METABOLISM) | -3.31 | 0 | 2.32 | 0.007 |
| REACTOME_METABOLISM_OF_COFACTORS | -2.17 | 0.013 | 2.31 | 0.007 |
| [REACTOME_PEROXISOMAL_LIPID_METABOLISM](http://www.gsea-msigdb.org/gsea/msigdb/human/geneset/REACTOME_PEROXISOMAL_LIPID_METABOLISM) | -2.59 | 0.001 | 2.22 | 0.011 |
| [REACTOME_MITOCHONDRIAL_BIOGENESIS](http://www.gsea-msigdb.org/gsea/msigdb/human/geneset/REACTOME_MITOCHONDRIAL_BIOGENESIS) | -3.48 | 0 | 2.2 | 0.012 |
| [KEGG_TRYPTOPHAN_METABOLISM](http://www.gsea-msigdb.org/gsea/msigdb/human/geneset/KEGG_TRYPTOPHAN_METABOLISM) | -2.61 | 0.001 | 2.18 | 0.014 |
| [HALLMARK_BILE_ACID_METABOLISM](http://www.gsea-msigdb.org/gsea/msigdb/human/geneset/HALLMARK_BILE_ACID_METABOLISM) | -2.79 | 0 | 2.16 | 0.015 |
| REACTOME_FOXO_MEDIATED_TRANSCRIPTION | -2.06 | 0.021 | 2.13 | 0.017 |
| [WP_MITOCHONDRIAL_COMPLEX_II_ASSEMBLY](http://www.gsea-msigdb.org/gsea/msigdb/human/geneset/WP_MITOCHONDRIAL_COMPLEX_II_ASSEMBLY) | -2.54 | 0.001 | 2.12 | 0.018 |
| [REACTOME_PINK1_PRKN_MEDIATED_MITOPHAGY](http://www.gsea-msigdb.org/gsea/msigdb/human/geneset/REACTOME_PINK1_PRKN_MEDIATED_MITOPHAGY) | -2.72 | 0 | 2.01 | 0.029 |
| [BIOCARTA_ETC_PATHWAY](http://www.gsea-msigdb.org/gsea/msigdb/human/geneset/BIOCARTA_ETC_PATHWAY) | -2.7 | 0 | 1.99 | 0.031 |
| WP_GLYCOLYSIS_AND_GLUCONEOGENESIS | -2.13 | 0.015 | 1.99 | 0.031 |
| [BIOCARTA_KREB_PATHWAY](http://www.gsea-msigdb.org/gsea/msigdb/human/geneset/BIOCARTA_KREB_PATHWAY) | -3.01 | 0 | 1.98 | 0.032 |
| [KEGG_FATTY_ACID_METABOLISM](http://www.gsea-msigdb.org/gsea/msigdb/human/geneset/KEGG_FATTY_ACID_METABOLISM) | -3.55 | 0 | 1.95 | 0.037 |
| HALLMARK_HEME_METABOLISM | -2.02 | 0.027 | 1.93 | 0.039 |
| [REACTOME_BETA_OXIDATION_OF_DECANOYL_COA_TO_OCTANOYL_COA_COA](http://www.gsea-msigdb.org/gsea/msigdb/human/geneset/REACTOME_BETA_OXIDATION_OF_DECANOYL_COA_TO_OCTANOYL_COA_COA) | -2.49 | 0.002 | 1.92 | 0.042 |
| REACTOME_MITOPHAGY | -2.18 | 0.012 | 1.92 | 0.042 |
| [REACTOME_TP53_REGULATES_METABOLIC_GENES](http://www.gsea-msigdb.org/gsea/msigdb/human/geneset/REACTOME_TP53_REGULATES_METABOLIC_GENES) | -2.67 | 0 | 1.9 | 0.044 |

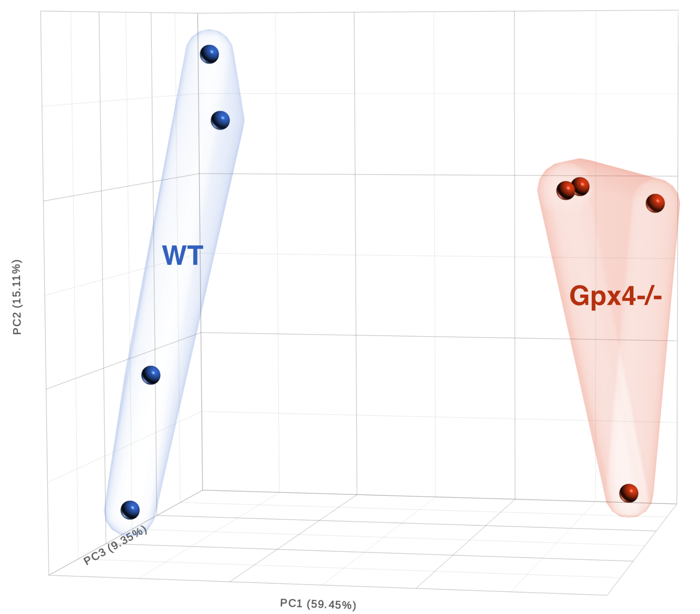

**Supplementary Figure S1. Transcriptomic variation between *Gpx4 KO* and WT control mouse hearts.** PCA plot across the cardiomyocyte transcriptome in *Gpx4*^-/-^ samples (n=4, red spheres) and WT controls (n=4, blue spheres).

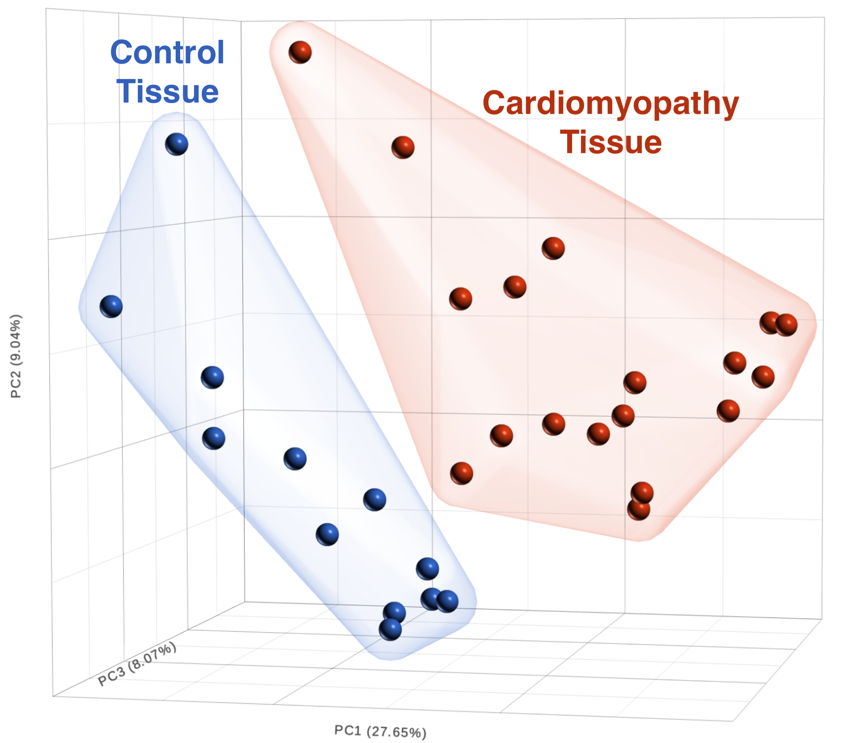

**Supplementary Figure S2. Transcriptomic variation between patients with diagnosed cardiomyopathy and controls.** PCA plot of the entire transcriptome in the cardiac tissue of patients diagnosed with cardiomyopathy (n=18, red spheres) and controls (n=12, blue spheres), displaying clear separation between the two groups.
